# Contrasting walking styles map to discrete neural substrates in the mouse brainstem

**DOI:** 10.1101/2023.04.19.537568

**Authors:** Audrey Worley, Alana Kirby, Sophie Luks, Tamara Samardzic, Brian Ellison, Lauren Broom, Alban Latremoliere, Veronique G VanderHorst

**Author notes:** Corresponding and lead author: Veronique VanderHorst MD PhD Center for Life Sciences, BIDMC, 3 Blackfan Circle, Suite 709, Boston, MA 02115.

## Abstract

Walking is a slow gait which is particularly adaptable to meet internal or external needs and is prone to maladaptive alterations that lead to gait disorders. Alterations can affect speed, but also style (the way one walks). While slowed speed may signify the presence of a problem, style represents the hallmark essential for clinical classification of gait disorders. However, it has been challenging to objectively capture key stylistic features while uncovering neural substrates driving these features. Here we revealed brainstem hotspots that drive strikingly different walking styles by employing an unbiased mapping assay that combines quantitative walking signatures with focal, cell type specific activation. We found that activation of inhibitory neurons that mapped to the ventromedial caudal pons induced slow motion-like style. Activation of excitatory neurons that mapped to the ventromedial upper medulla induced shuffle-like style. Contrasting shifts in walking signatures distinguished these styles. Activation of inhibitory and excitatory neurons outside these territories or of serotonergic neurons modulated walking speed, but without walking signature shifts. Consistent with their contrasting modulatory actions, hotspots for slow-motion and shuffle-like gaits preferentially innervated different substrates. These findings lay the basis for new avenues to study mechanisms underlying (mal)adaptive walking styles and gait disorders.

**Graphical abstract:** 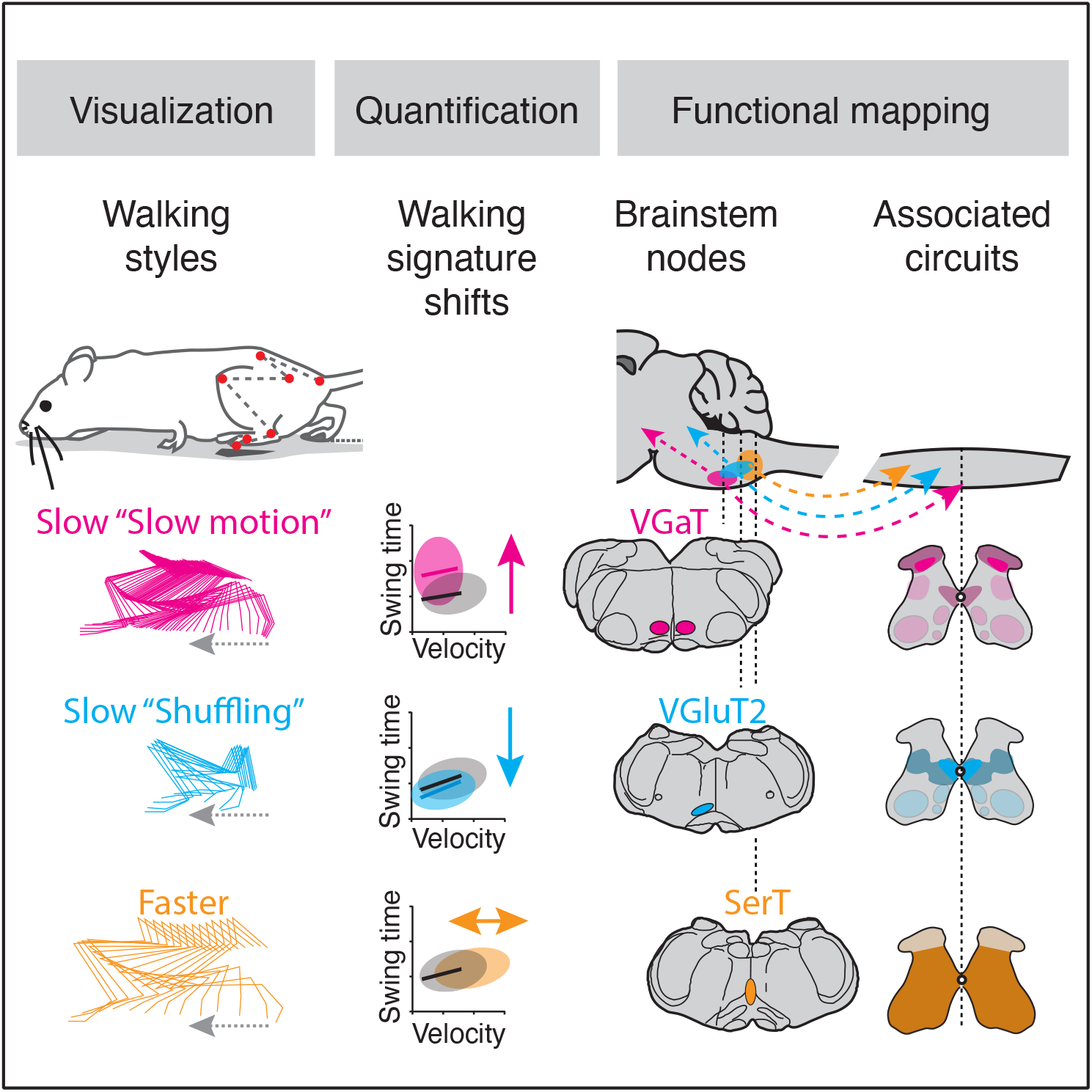

## Introduction

Walking is a slow gait which is particularly adaptable and is prone to maladaptive alterations that lead to gait disorders. Slowed walking speed is a cardinal marker of gait disorders in neurological conditions, systemic disease and aging (1–4). However, speed alone is too non-specific to disentangle underlying pathophysiology. Rather, characteristic differences in walking style (a. k. a. “phenomenology”) form the key for classification of gait disorders (5). For example, slowing with small shuffling steps is a feature of parkinsonian gait, while slowing with erratic steps would be classified as ataxic, and both these slowed styles look distinct from normal slowed walking as during window shopping. Classification of gait disorders is built on centuries of clinical observations (6–8), but objective classification of walking styles has remained challenging. Furthermore, insights into neural substrates that drive their key features are limited both in human subject and pre-clinical research, limiting the development of more targeted therapeutic approaches.

Multiple specialized CNS circuits govern gait in general and while insight in its network organization is growing, the neural basis for variations in style remain elusive. Conceptually, to adjust to external or internal demands, central pattern generator networks in the spinal cord that set basic locomotor patterns, including walking, function in a flexible way (9–12) guided by peripheral and supraspinal input (13–16). Recent work has demonstrated that a diverse set of cell-type specific supraspinal systems that reside in the brainstem mediate halting or initiation of movement, or modulate gait speed including via frequency changes and transitions between walking, trotting and running patterns (17–24). While these studies point to the importance of the brainstem in gait control, they were not designed to elicit, detect or quantify walking styles.

We set out to modulate neurons in the medial reticular formation (mRF) near the pontomedullary junction in the brainstem and test their role in walking styles mice. This area contains a large population of neurons that directly innervate the spinal cord, including lumbar levels important for hindlimb control (25). Thus, this region, among other sites that may modulate gait (26), is uniquely positioned to directly modulate walking style. We chose a reversible chemogenetic approach employing the hM3Dq receptor, which acts via canonical Gq protein signaling to amplify intrinsic neuronal activity when activated by clozapine-N-oxide (CNO)(27) or its metabolites (28). Where relevant we complemented this with a temporally precise optogenetic approach to probe gait phase-specificity (29). We separated roles of inhibitory, excitatory and serotonergic classes, which have all been implicated in locomotion, motor or tone control (18, 20, 23, 30–33), by transfecting the mRF with hM3Dq in vesicular GABA transporter (VGaT)-, vesicular glutamate transporter 2 (VGluT2)-ires-cre knock-in mice (34) or serotonin transporter (SerT)-cre transgenic mice (35).

Quantification of walking styles has been challenging as walking metrics invariably change when walking speed changes, even in the absence of style changes (Fig. 1), and as mice tend to trot and run rather than walk under standard testing conditions (36, 37). We developed and validated a quantitative “walking signature” assay (36, 38) to separate changes in walking style from changes in speed. Spatiotemporal walking metrics depicted as a function of speed (Fig. 1) adhere to stable signatures during normal speed modulation, while signature shifts of one or several gait metrics signify style changes in disease and aging models (38, 39). When applied to hindlimbs, walking signatures are translatable between quadruped and biped human models (36, 38). This readout, complemented with EMG analyses and motor tests, enabled us to classify and quantify walking styles.

**Figure 1:**
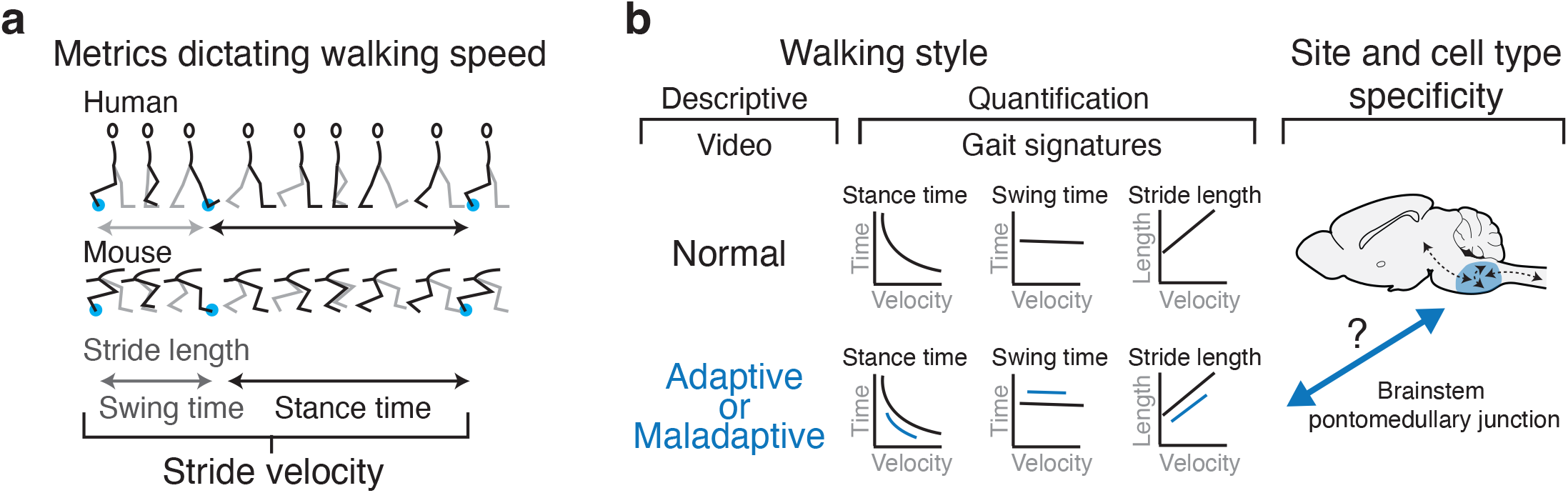
Which neural substrates modulate walking style? **a:** Schematic of key spatial-temporal footfall metrics that dictate walking speed in humans and mice. **b:** Gait signatures of metrics that dictate walking speed can serve to distinguish normal from (mal)adaptive walking styles. A key question remains which neural substrates modulate each of these signatures. In this study the focus is on substrates near the pontomedullary junction in the brainstem.

Finally, we probed the localization principle, a classical concept in neuroscience of a spatial organization into functional compartments (8), which remains controversial for the mRF. We dispersed small hM3Dq transfection sites across the mRF, and grouped and mapped these based upon type and magnitude of walking style. This functional assay contrasts with standard approaches that employ standardized anatomical boundaries as the basis for analyses. Such boundaries are notoriously ambiguous in the mRF and depend upon morphology (25, 40) to which functionally, molecularly or neurochemically distinct entities may not adhere (18, 25, 41, 42). This design uncovered spatially restricted mRF hotspots that differentially modulated walking style, the circuit specificity of which we further verified.

## Results

### Chemogenetic mRF modulation elicited different walking styles

To determine whether enhanced activation of inhibitory, excitatory or serotonergic mRF neurons induced different walking styles, we injected small volumes (median 35nl) of AAV8-DIO-hM3Dq-mCherry uni- or bilaterally into subregions of the mRF of VGaT-ires-cre, VGluT2-ires-cre and SerT-cre mice (Fig. 2a). Following transfection, we video-captured self-paced over-ground walking under baseline (saline i.p.) and activation conditions (0.3mg/kg CNO i.p; Figure 2-Figure supplement 1; Table S1 for dose and time titration) using a walkway that incites mice to walk rather than run (36, 38).

**Figure 2:**
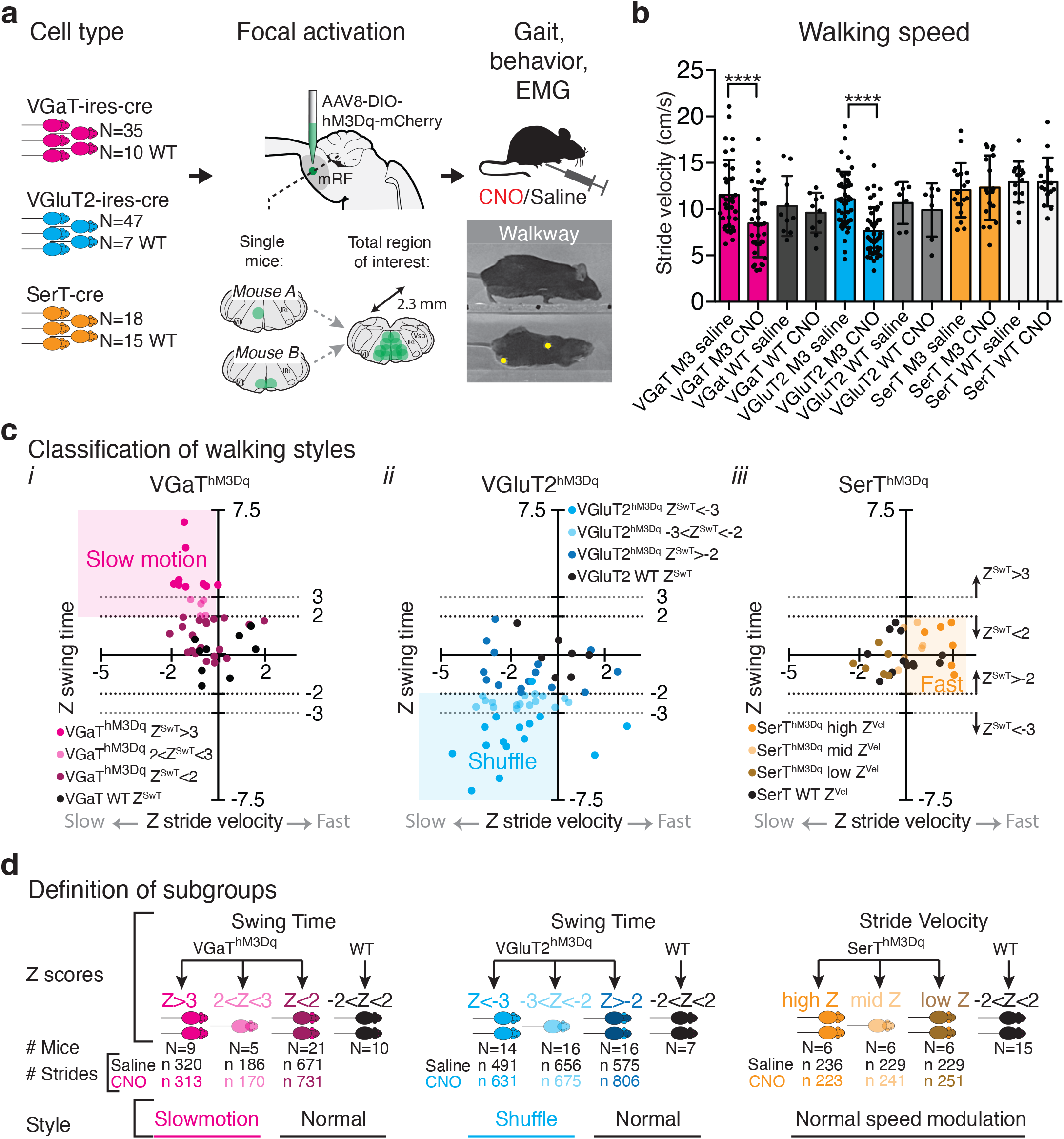
Chemogenetic activation of VGaT, VGluT2 or SerT mRF neurons elicited contrasting walking styles. **a:** Small sites with inhibitory (VGaT), excitatory (VGluT2) or serotonergic (SerT) brainstem neurons were selectively transfected with hM3Dq, at the group level covering the entire region of interest. Walking gait, motor behavior and electrophysiological data were collected in control (saline) and activation (CNO) conditions. **b:** Average stride velocity decreased following hM3Dq activation in experimental VGaT and VGluT2 cohorts, but not their controls (2-tailed paired t-test; Table S2; error bars represent standard deviation). **c:** hM3Dq activation induced slow motion and shuffle-like styles or faster walking (Videos 1-3) in subsets of experimental mice. Z-scores for stride velocity and swing time formed the basis for classification of style in each experimental and control mouse. Each dot represents one mouse. **d:** Z scores were applied to divide each cohort into sub-groups for in depth gait, motor and EMG analyses, and for transfection site and circuit mapping. See Figure 2-Figure supplement 1 for dose and time response titration.

We found that mRF activation decreased speed in both VGaT and VGluT2 cohorts (Fig. 2b; Table S2). We observed a “slow motion-like” (*slomo*) style with slow, long strides in the VGaT cohort (Video S1), and a “shuffling-like” (*shuffle*) style with quick, short strides in the VGluT2 cohort (Video S2). Importantly, these styles only involved a subset of mice in each cohort.

To objectively separate these slowed walking styles from each other and from slowed walking without style changes, we employed Z-scores for hindlimb swing time (time a limb is off the ground; Z^SwT^) and stride velocity (Z^StrV^) that for each mouse reflected the change between its baseline and CNO conditions relative to variation in WT littermates. Scatterplots of these scores (Fig. 2c) revealed that *slomo* and *shuffle* styles were distinct from normal walking based upon extreme values of Z^SwT^, positive for *slomo* style and negative for *shuffle* style. In contrast, activation of SerT neurons did not reveal an unusual style and did not change group-wise average stride velocity (Fig. 1b) or swing time (Z^SwT^) scores (Fig. 2c). However, in a subset of mice stride velocity (Z^StrV^) increased (Video S3; Fig. 2b), a feature that did not stand out in VGaT and VGLuT2 cohorts. In WT controls, CNO did not perturb gait metrics (Fig. 2c; black dots). CNO in experimental mice did not induce a state of immobility.

These findings indicate that the mRF is capable of driving different walking styles in a cell type specific way. Furthermore, transfection site rather than size alone may drive heterogeneity in gait style effect size as we used small AAV volumes distributed across a larger region of interest (Fig. 2c). To test this, we employed relevant Z-scores to define “positive” and “negative” sub-groups within each cell type specific cohort (Fig. 2f). Within each sub-group we then studied gait patterns, walking signatures, and muscle activation patterns. We then mapped “positive” and “negative” cell type specific transfection sites within each cohort and contrasted “positive” sites between cohorts.

### *Slomo* walking is defined by upward shifts in select gait signatures

To determine which gait features distinguish *slomo* style from normal (slow) walking, we first verified the gait pattern as defined by footfalls of all 4 limbs (Fig. 3a). In the Z^SwT^>3 (*slomo*) group footfall patterns were consistent with walking gait both at baseline and with VGaT^hM3Dq^ activation.

**Figure 3:**
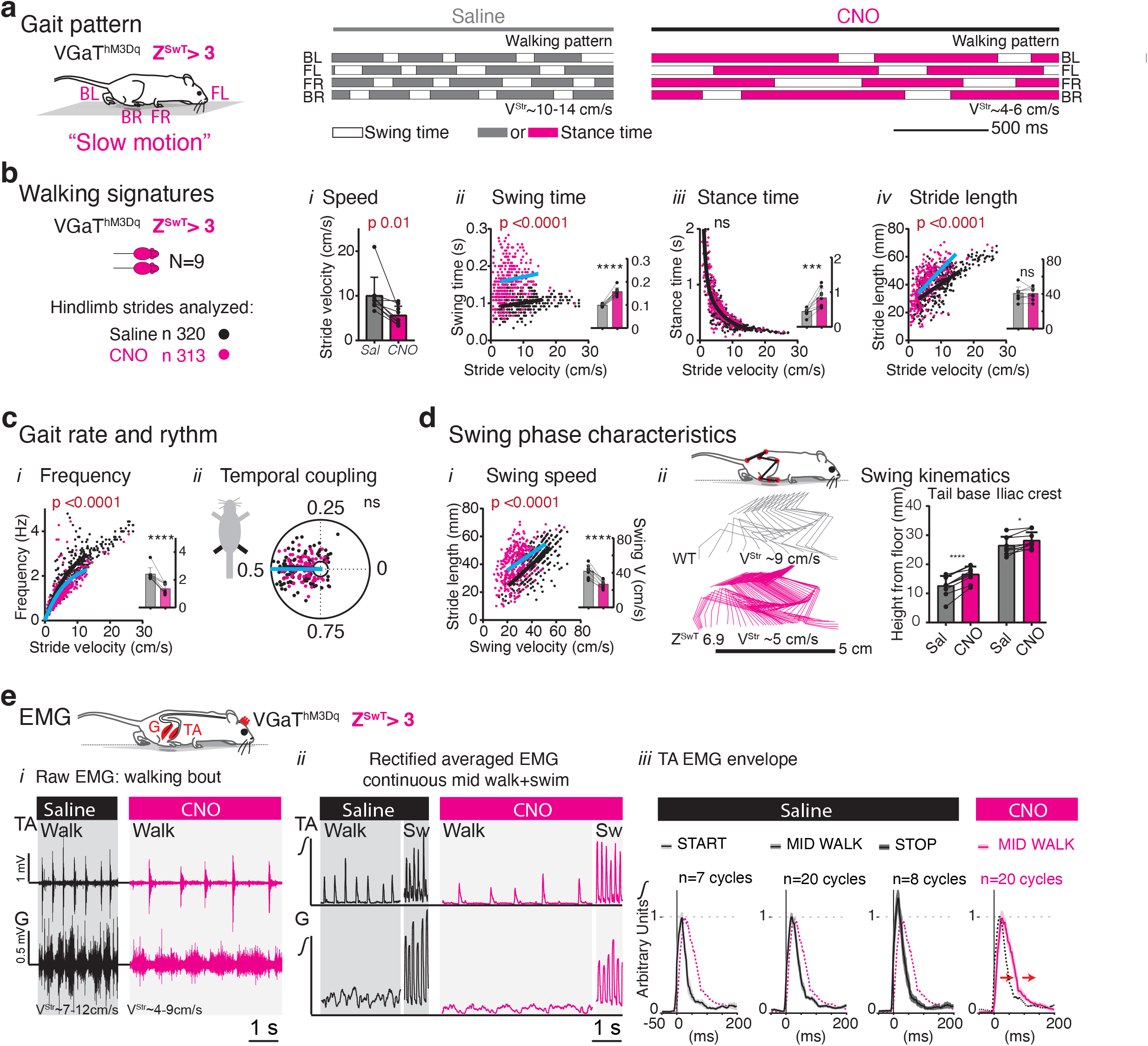
*Slomo* walking style was characterized by walking signature shifts and changes in EMG metrics that dictate the swing phase. VGaT^hM3Dq^ *slomo* subgroup (Z^SwT^>3; See Figure 3-Figure supplement 1 for Z^SwT^<2 *non-slomo* group). **a:** Footfall patterns in saline and CNO conditions represent walking gait. **b-d:** Scatter plots depict gait metrics as a function of stride-to-stride velocity in saline (black) or CNO (color) conditions. Each dot represents one hindlimb stride. For strides that fell within walking range, a sum of squares F-test was used to assess whether saline and CNO datasets shared regression lines (i.e. walking signatures; number of analyzed strides listed in a; α 0.001; black: baseline or baseline and CNO conditions are shared; color: signature shift). Bar graphs of averaged data that was not corrected for speed illustrate inter-animal homogeneity (2-tailed paired t-test; α 0.05; bars: SD). Polar plot shows temporal hindlimb coupling (radial axis 3-16cm/s; Watson and Williams test; α 0.05). Stick figures of the left 5^th^ metatarsal, ankle, knee, trochanter, iliac crest and tail base from representative cases illustrate the trajectory of the hindlimb during the swing phase. Table S3 for statistical details. **e:** (i-iii) Raw and rectified smoothened EMG during walking and swimming of a representative mouse (Z^SwT^=3.7; bilateral ventral mRF transfection; left EMG). See Figure 3-Figure supplement 2 for group EMG data; Table S4 for statistical details. (iv) EMG envelope, averaged from multiple TA bursts (+/- SEM) of start, steady walking and stop strides in the control condition (black) and steady walking with CNO (magenta). Note delayed ramp up and ramp down with CNO. Data are normalized to baseline TA bursts during continuous walking and aligned to TA burst onset.

Next, we examined hindlimb footfall metrics that together dictate speed, i.e. stride length, swing and stance time and their derivatives. As these metrics change as a function of walking speed alone (Fig. 1) and as speed changed in all experimental sub-groups, we had to account for speed related changes. We employed walking “signatures” to distinguish changes in speed alone (i.e. metrics abide by baseline signatures) from style changes, which are disproportionate to speed-related changes alone (i.e. signature shifts away from baseline) (36). In each sub-group, we compiled stride-to-stride metrics from baseline and CNO conditions as a function of stride velocity. Data within walking speed range was fitted to the simplest metric-specific regression model to determine whether baseline and CNO data fitted to a shared curve using an extra sum of squares F test. The results show (Fig. 3b-d) that decreased walking speed following VGaT^hM3Dq^ activation in the Z^SwT^>3 (*slomo*) group was accompanied by select shifts in gait signatures. As expected, swing time increased with an upward signature shift between baseline and CNO. In contrast, while stance time increased, it did so in proportion to decreased stride velocity (no shift). Stride length was significantly longer than expected for speed (shift). Stride frequency, derived from swing and stance time, decreased out of proportion to slowed speed, driven by increased swing time. Temporal coupling of the hindlimbs did not change, as shown by polar plots (Fig. 3c ii), in line with preserved footfall patterns. Averaged gait metrics of each mouse, which represent data irrespective of changing speed, showed that changes were homogeneous within subgroup (inserts Fig. 3b-d). We reproduced these results in the mild subgroup (2<Z^SwT^<3; Table S3). In contrast, in the Z^SwT^<2 (*non-slomo*) group, stride velocity decreased, but gait metrics followed normal speed-dependent rules without signature shifts (Figure 3-Figure supplement 1). In WT control mice, CNO did not affect stride velocity or perturb signatures (Table S3). Next, as prolonged swing time drove *slomo* style, we assessed the swing phase in more detail. In the Z^SwT^>3 (*slomo*) group (Fig. 3d), but not the *non-slomo* group (Figure 3-Figure supplement 1), swing speed decreased for a given stride length, as also illustrated by a higher number of sticks in representative stick figures derived from hindlimb positions in subsequent video frames. Finally, we quantified simple postural changes to assess whether these are part of *slomo* style or whether these occur independently. At the start of the swing phase the base of the tail and less so iliac crest were carried higher following VGaT^hM3Dq^ activation than at baseline in both *slomo* and Z^SwT^<2 (*non-slomo*) sub-cohorts (Fig. 3d; Figure 3-Figure supplement 1), implicating that these postural features were not a specific component of *slomo* gait.

### Activation of the anterior tibial muscle is prolonged during *slomo* walking

To assess whether prolonged swing phase that characterizes *slomo* style is due to changes in timing of hindlimb muscle activation and/or an imbalance in extensor and flexor hindlimb muscle activity, we employed electromyography (EMG) of Gastrocnemius (G) and Anterior Tibial (TA) muscles in a subset of VGaT^hM3Dq^ and control mice. During the swing phase of baseline walking, ankle dorsiflexor TA is active and plantar flexor G is silent, with TA burst duration being relatively unperturbed by changing velocity (43). During *slomo* walking, raw and averaged EMG showed a striking increase in cycle duration (swing plus stance time) with preserved TA-G pattern (Fig. 3e). TA burst duration during steady walking was prolonged in the Z^SwT^>2 (*slomo*) mice with CNO compared to baseline (EMG envelope of n=20 strides each; Fig. 3e) with a slower ramp on and ramp off. The TA EMG envelope during steady *slomo* walking did not resemble EMG envelopes of (per definition slow) starting and stopping strides at baseline. Group analyses confirmed prolonged TA burst duration in the Z^SwT^>2 (*slomo*) group, but not Z<2 and WT groups, with CNO compared to baseline (Figure 3-Figure supplement 2; Table S4) and showed that Z-scores for TA burst duration correlated with Z^SwT^ from gait analyses (Figure 3-Figure supplement 2, Table S5). These characteristics applied to other hind- and forelimb muscles (Figure 3-Figure supplement 2). As for effects on EMG amplitude, average TA peak burst amplitude was variable but did not change in any Z^SwT^-defined subgroup whereas TA interburst amplitude decreased in both high (*slomo*) and low Z^SwT^ groups but not WT (Figure 3-Figure supplement 2; Table S4). Decreased interburst amplitude in the CNO condition indicates that there was no co-activation of TA and G muscles in the Z^SwT^>2 (*slomo*) or other groups. As expected, Z-scores for TA amplitude did not correlate with Z^SwT^ (Figure 3-Figure supplement 2, Table S5). Altogether, *slomo* style is mediated by prolonged muscle burst activity with preserved temporal coordination between flexors and extensors and this is modulated independently from EMG interburst amplitude (tone).

### Do *slomo* mice have deficits in other (loco)motor tasks?

As specific neural substrates control different locomotor modes (12) we recorded TA and G EMG activity during swimming, which is characterized by a higher cycle frequency than walking. During swimming, TA cycle frequency decreased slightly in the Z^SwT^>2 (*slomo*) group with CNO compared to baseline (Table S4), but TA burst or interburst duration alone did not change significantly between conditions in any Z^SwT^ subgroup. We also tested for each Z^SwT^-defined subgroup whether VGaT^hM3Dq^ activation affected learned motor tasks, balance, and motor planning using rotarod, beam, and horizontal ladder, respectively (Table S6). With CNO, latency to drop off a rotarod was shortened in both Z^SwT^>2 and Z^SwT^<2 groups compared to saline, i.e. not specific for *slomo* mice. Activation did not affect errors on beam or ladder in Z^SwT^>2 (*slomo*) or Z^SwT^<2 groups, though *slomo* mice moved slowly with CNO during these tests (Video S1). Thus, while slowed movements can affect other motor tasks, performance of these tasks is not affected when the task can be performed at preferred pace.

### Shifts in gait signatures that define *Shuffle* style contrast those of *Slomo* walking

To study walking and motor function in VGluT2^hM3Dq^ subgroups we applied the same paradigms as in the VGaT cohort. During *shuffle* gait, footfall patterns remained consistent with walking (Fig. 4a). VGluT2^hM3Dq^ activation in the Z^SwT^<-3 (*shuffle*) group caused shortening of swing time, stance time and stride length out of proportion to slowed speed as demonstrated by downward shifts of their respective walking signatures (Fig. 4b). Due shortening of both swing and stance times, stride frequency (cadence) was higher than expected for stride velocity. Polar plots revealed that hindlimb coupling was more variable following VGluT2^hM3Dq^ activation than at baseline in the Z^SwT^<-3 (*shuffle*) group (Fig. 4c; Table S8), in line with asymmetries seen in footfall pattern (Fig. 4a). During the swing phase, stride length shortened with preserved swing speed (Fig. 4d), in contrast to *slomo* mice with stable stride length but decreased swing speed. Postural changes including lowered tail base and iliac crest accompanied *shuffle* gait in Z^SwT^<-3 sub-group. In the Z^SwT^>-2 (*non-shuffle*) group (Figure 4-Figure supplement 1; Table S8), walking speed decreased, but without changes in swing and stance time signatures or posture, and with subtle changes in stride length signature and hindlimb coupling. In WT littermates, CNO did not affect velocity or gait signatures (Table S8). In none of the Z^SwT^-defined experimental or control groups (Table S9) CNO activation changed performance on rotarod, beam, or ladder. Summarizing, changes in walking signatures and postural changes that characterize *shuffle* gait are distinct from similarly slowed *slomo* gait.

**Figure 4.**
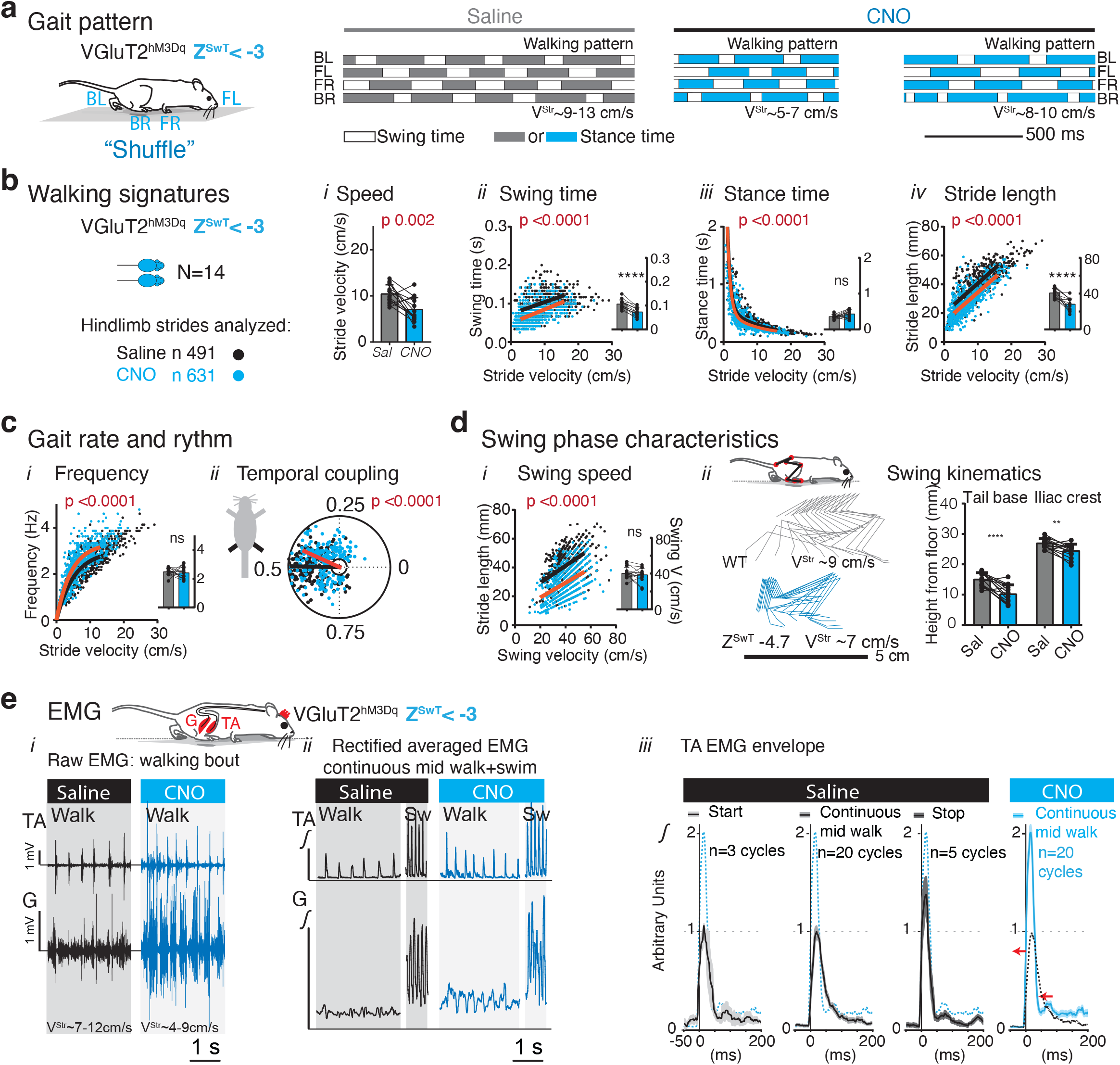
: *Shuffle* style was characterized by gait signature shifts and EMG changes that are distinct from *slomo* style. VGluT2^hM3Dq^ *shuffle* subgroup (Z^SwT^<-3 See Figure 4-Figure supplement 1 for Z^SwT^>-2 group). **a**-**e:** See Legend Fig. 3. Representative mouse in (**e** i-iii) with Z^SwT^=-4.7; EMG ipsilateral to unilateral transfection. See Figure 4-Figure supplement 2 for group EMG data; Tables S8 and S10 for statistical details.

### Anterior tibial and gastrocnemius muscles co-activate during *shuffle* walking

EMG of TA and G muscles in *shuffle* mice (VGluT2^hM3Dq^ Z^SwT^<-2) revealed co-activation of TA and G muscles during the stance phase due to increased TA interburst activity (Fig. 4e; Table S10 for group data). TA EMG envelopes during steady walking demonstrated decreased TA burst duration with a more rapid ramp down with CNO compared to saline (Fig. 4e iii; Figure 4-Figure supplement 2; Table 10S for group data). This was less pronounced in the Z^SwT^>-2 (*non-shuffle*) group (Table 10S). In addition, in *shuffle* mice the TA envelope revealed a larger amplitude with CNO versus baseline, which is a characteristic of halting steps at baseline (Fig. 4e iii). However, peak TA burst amplitude increased significantly with CNO compared to saline in both Z^SwT^<-2 (*shuffle*) and Z^SwT^>-2 (*non-shuffle*) groups (Table S10). In line with this, TA Z^BurstAmplitude^ did not correlate with Z^SwT^ (Figure 4-Figure supplement 2; Table S5). This means that the increase in TA burst amplitude is not specific for *shuffle* mice. During swimming, neither TA burst and interburst duration nor cycle frequency changed between baseline and CNO in the Z^SwT^<-2 (*shuffle*) group. There were no significant changes in G EMG parameters at the group level, though G data was underpowered due to quality constraints (Table S10).

### Serotonergic activation modulates walking speed without shifts in walking signatures

In SerT^hM3Dq^ mice with high Z^StrV^ scores, indicative of increased speed, footfall patterns remained consistent with walking and slow trotting gaits in saline and CNO conditions (Fig. 5; Table S12). In high Z^StrV^ (*fast*) (Fig. 5) and low Z^StrV^ (*slow*) groups (Figure 5-Figure supplement 1), CNO activation changed spatial and temporal gait metrics that dictate speed in opposite directions, but in proportion to speed, i.e. without curve shifts. Left-right temporal hindlimb coupling and posture of iliac crest and tail base did not change following CNO activation in any Z^StrV^ subgroup (Table S12). In WT littermates, CNO did not affect speed or walking signatures (Table S12). EMG of TA and G muscles revealed a trend for increased peak amplitude and area under the curve of TA bursts during steady walking in the high Z^StrV^ (*fast*) group (Fig 5e; for group data Figure 5-Figure supplement 2 and Table S13), but without the changes in timing or co-activation seen in *slomo* or *shuffle* subgroups. Neither timing nor amplitude changed in the low Z^StrV^ *(slow)* group (Table S13). During swimming, TA EMG metrics did not change in high or low Z^StrV^ groups with CNO versus baseline, except for a trend towards increased cycle frequency in the high Z^StrV^ (*fast*) group (Table S13). CNO induced no changes in G EMG metrics or in performance on rotarod, beam, and ladder in any of the Z^StrV^-defined experimental or control groups (Table S14). These data indicate that SerT neurons modulate gait speed in accordance with normal gait signatures, i.e. without style modification.

**Figure 5:**
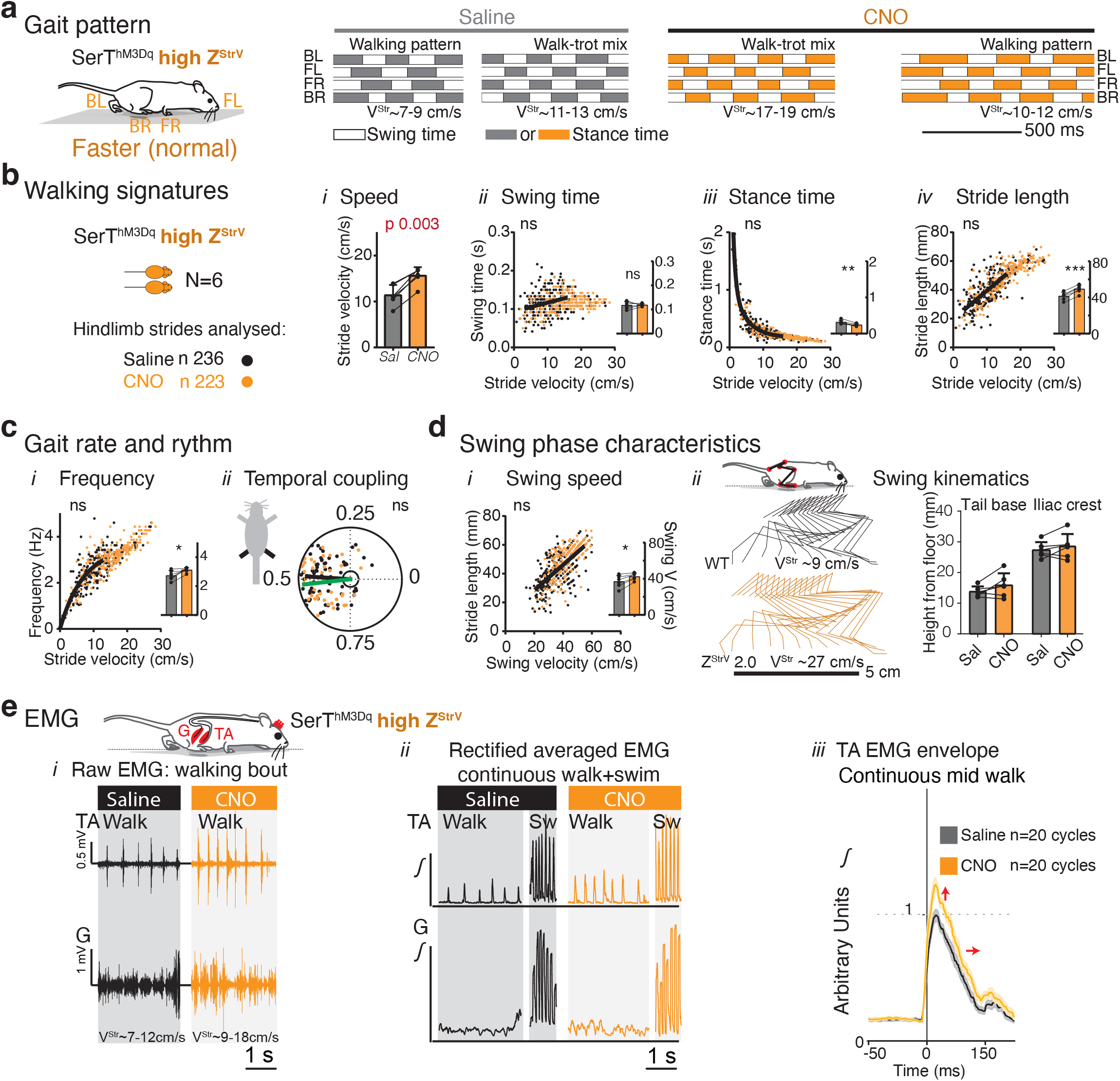
mRF SerT^hM3Dq^ activation modulated gait speed without style changes. SerT^hM3Dq^ subgroup with highest ranked Z^StrV^ (See Figure 5-Figure supplement 1 for lowest ranked Z^StrV^ group). **a**-**d**: see Legend Fig. 3. Representative mouse in (**e** i-iii) with Z^StrV^ 1.5, midline transfection in raphe obscurus; EMG: left leg. See Figure 5-Figure supplement 2 for group EMG data; Tables S12 and S13 for statistical details.

### Walking styles map to different mRF subregions

We showed that mRF VGaT^hM3Dq^ or VGluT2^hM3Dq^ activation elicited distinct walking styles, while SerT^hM3Dq^ activation sped up walking. However, the “positive” and “negative” subgroups indicated that this involved subsets of experimental mice. This was expected as AAV transfection targeted discrete, spatially distinct uni- and bilateral sites within a larger mRF region of interest. We took advantage of this heterogeneity to determine whether distinct sites drive each style and whether this required uni- or bilateral activation. To study this we compiled VGaT, VGluT2 and SerT hM3Dq-mCherry transfection sites for each Z^SwT^ or Z^StrV^ defined sub-group (Fig. 6; Figure 6-Figure supplement 1 for reconstruction methodology; Figure 6-Figure supplement 2 for detailed reconstruction).

**Figure 6:**
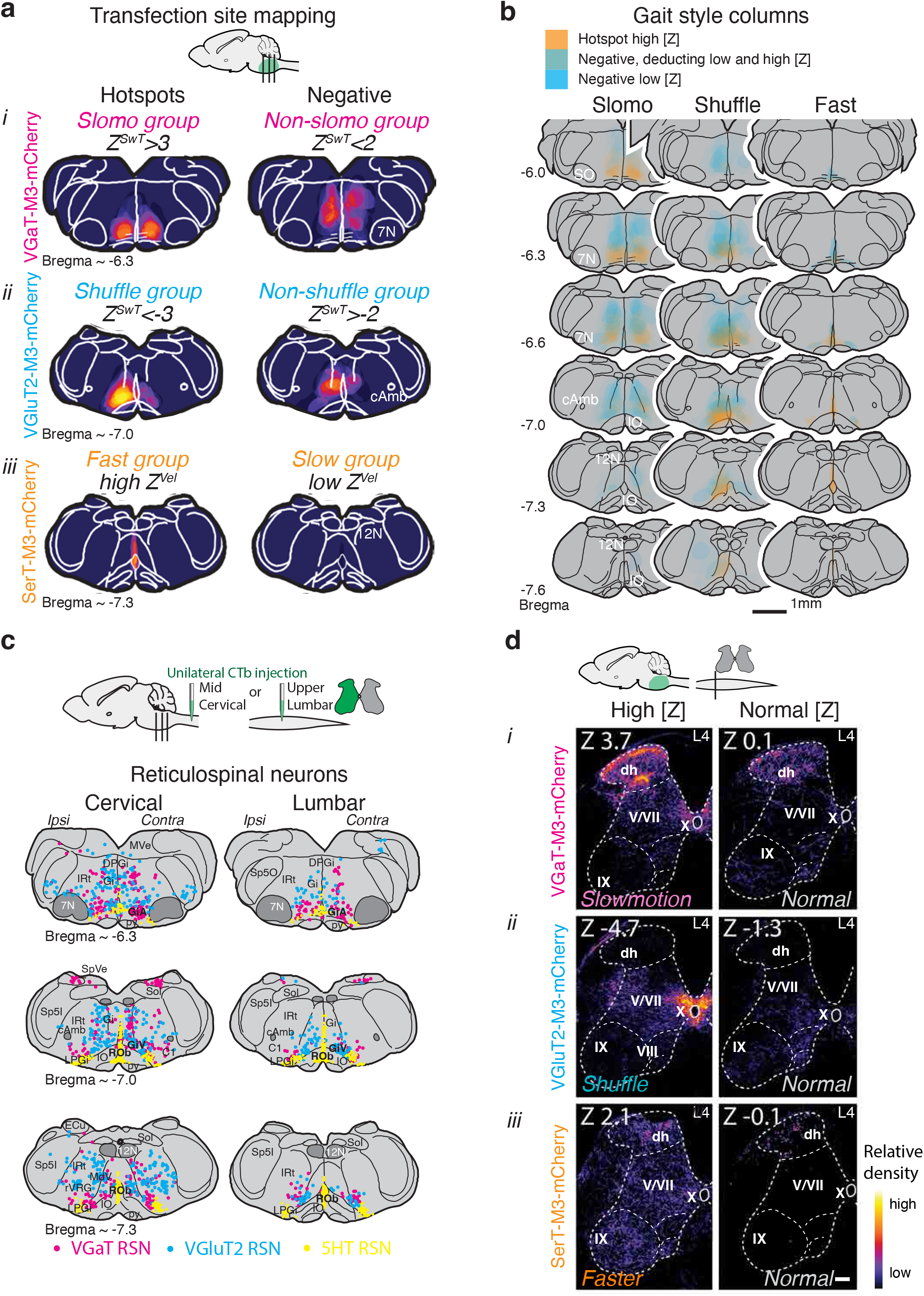
*Slomo*, *shuffle*, and *fast* walking styles localized to discrete sites in the mRF. **a:** *Slomo*, *shuffle* or *fast* walking styles map to distinct mRF levels and sites, as shown by gait Z score-grouped composite gradient based value maps of hM3Dq-mCherry transfection sites in VGaT, VGluT2 and SerT cohorts. See Figure 6-Figure supplement 1 for methodology; See Figure 6-Figure supplement 2 for full rostrocaudal compilations. **b:** Visual subtraction of negative (blue) from positive (orange) sub-group transfection sites further defines the extent of *slomo*, *shuffle*, and *fast* columns. **c:** Inhibitory and excitatory reticulospinal neurons (RSN) form clusters in the centers of *slomo* and *shuffle* hotspots, respectively, but these clusters represent only a small portion of the reticulospinal system. **d:** Projections from transfection sites of select *slomo*, *shuffle* or *fast* mice versus mice without these style changes illustrate differences in density and/or localization of projections to the lumbar enlargement. Bar=100µm. See Figure 6-Figure Supplement 3 for examples of ascending projections from hotspots and nearby sites.

In the VGaT cohort, Z^SwT^>3 (*slomo*) sites localized to the ventromedial caudal pons and were all bilateral. Bilateral Z^SwT^<2 (*non-slomo*) sites were located dorsally, or ventrally at more caudal levels. Unilateral sites, even when purposefully centered at the *slomo* hotspot, yielded Z^SwT^<2. Post-hoc analyses in unilateral cases did not reveal asymmetries in temporal (swing time) or spatial (stride length) metrics (Figure 3-Figure supplement 3; Table S7). As Z^SwT^ did not correlate with TA muscle amplitude changes, we expected the VGaT^hM3Dq^ region eliciting a drop in EMG amplitude to be distinct from the *slomo* Z^SwT^ hotspot. Indeed, while decreased Z^InterburstAmplitude^ mapped to the ventral mRF, this region extended caudally far beyond the *slomo* hotspot and involved both bilateral and unilateral sites (Figure 6-Figure supplement 2). This supports the notion that these distinct functions are mediated by different inhibitory neuronal populations which only partially overlap.

In the VGluT2^hM3Dq^ cohort, Z^SwT^<-3 (*shuffle*) sites also localized ventrally, but caudal to the VGaT^hM3Dq^ *slomo* hotspot. Z^SwT^>-2 (*non-shuffle*) sites localized more dorsally or laterally. In contrast to *slomo* gait, unilateral sites were sufficient to induce *shuffle* style. Post-hoc gait analyses to assess for asymmetries (Figure 4-Figure supplement 3; Table S11) revealed that activation of *uni*lateral VGluT2^hM3Dq^ Z^SwT^<-3 (*shuffle*) sites shifted key temporal (swing time) and spatial (stride length) gait signatures in a bilateral manner. Shifts in swing time signatures were larger *contra*-than ipsilateral to the transfection site, suggesting a crossed circuit mechanism, while stride length signatures changed symmetrically, pointing to a separate mechanism.

In the SerT cohort, transfection sites of high (*fast*) and low (*slow*) Z^StrV^ groups localized to caudal and rostral poles of the midline medullary raphe, respectively. As most transfection sites only involved the midline, we did not examine gait asymmetries.

Subtraction of transfection sites with and without change in walking style in each of the cell type specific cohorts (Fig. 6b) showed that dorsoventrally, ventromedially and rostrocaudally restricted columns drive each of the walking styles, with the centers of these columns being distinct from each other.

### Sites that drive contrasting walking styles differ in connectivity

The basis for differences in *slomo* and *shuffle* walking styles may lie in distinctions of pattern and/or density of efferent connections originating from their respective hotpots. While these may include local, ascending and descending circuits, to probe the concept of heterogeneity in projections we focused on spinal substrates as these govern baseline signatures and form the most direct route for modulation of style.

We first verified that *slomo* and *shuffle* hotspots project to the spinal cord and are predominantly VGaT^+^ and VGluT2^+^, respectively. Combined retrograde tracing with Cholera

Toxin subunit B (CTb) from the cervical or lumbar spinal cord with *in-situ* hybridization (ISH) revealed that both cervical and lumbar enlargements were innervated by the hotspots (Fig. 6c), which was expected as both hind- and forelimbs serve walking and share similarities in gait signatures (38). Of note, many reticulospinal neurons were present beyond the hotspots, involving sites that induced slowed walking but without changes in walking style.

We then screened hM3Dq-mCherry projections from mice with contrasting Z^SwT^ or Z^StrV^ scores and discrete transfection sites for heterogeneity in lumbar projections (Fig. 6d). VGaT^hM3Dq-mCherry^ projections from the Z^SwT^>3 (*slomo*) hotspot densely innervated dorsal horn and intermediate zone and less so ventral horn. VGluT2^hM3Dq-mCherry^ projections from the Z^SwT^<-3 (*shuffle*) hotspot most densely targeted the intermediate zone of the lumbar cord, especially area X, and also less so ventral horn. Sites negative for *slomo* and *shuffle* style involved lateral or medial ventral horn (both VGluT2 and VGaT) or superficial dorsal horn (VGaT), but without distinct pattern and density, in line with different sites of origin. SerT^hM3Dq-mCherry^ sites in the high (*fast*) Z^StrV^ group innervated the spinal ventral, intermediate and dorsal horn and the low (*slow*) Z^StrV^ group the dorsal horn only. From deduction it follows that the high Z^StrV^ effects are mediated through the ventral and intermediate gray. *Slomo* or *shuffle* hotspots in mice with a prominent style phenotype did not project heavily to mid- and forebrain sites in sharp contrast to sites that involved more dorsal mRF regions (Figure 6-Figure supplement 3). Thus, efferent connections from small hM3Dq-mCherry transfection sites that mediated different gait styles stood out from each other as well as from nearby sites.

### Hotspots for *slomo* and *shuffle* walking target distinct neuronal substrates

The site predilection of projections to the lumbar spinal cord from *slomo* and *shuffle* hotspots was unexpected as we had predicted primary involvement of ventral horn motoneuronal cell groups. To verify whether brainstem sites identified as hotspots indeed preferentially connected to dorsal horn (*slomo*), intermediate zone (*shuffle*) and ventral horn (*fast*), respectively, we injected CTb tracer unilaterally into these sites in mice with [high] Z scores, following completion of behavioral studies (Fig. 7a; Figure 7-Figure supplement 1). Retrograde CTb labeling of hM3Dq-mCherry transfected neurons in the hotspots confirmed ipsilateral connections to the respective spinal sub-regions. We further verified cell type specificity in the transfected hotspots and in transfected spinal boutons using ISH and IHC (Figure 7-Figure supplement 1).

**Figure 7:**
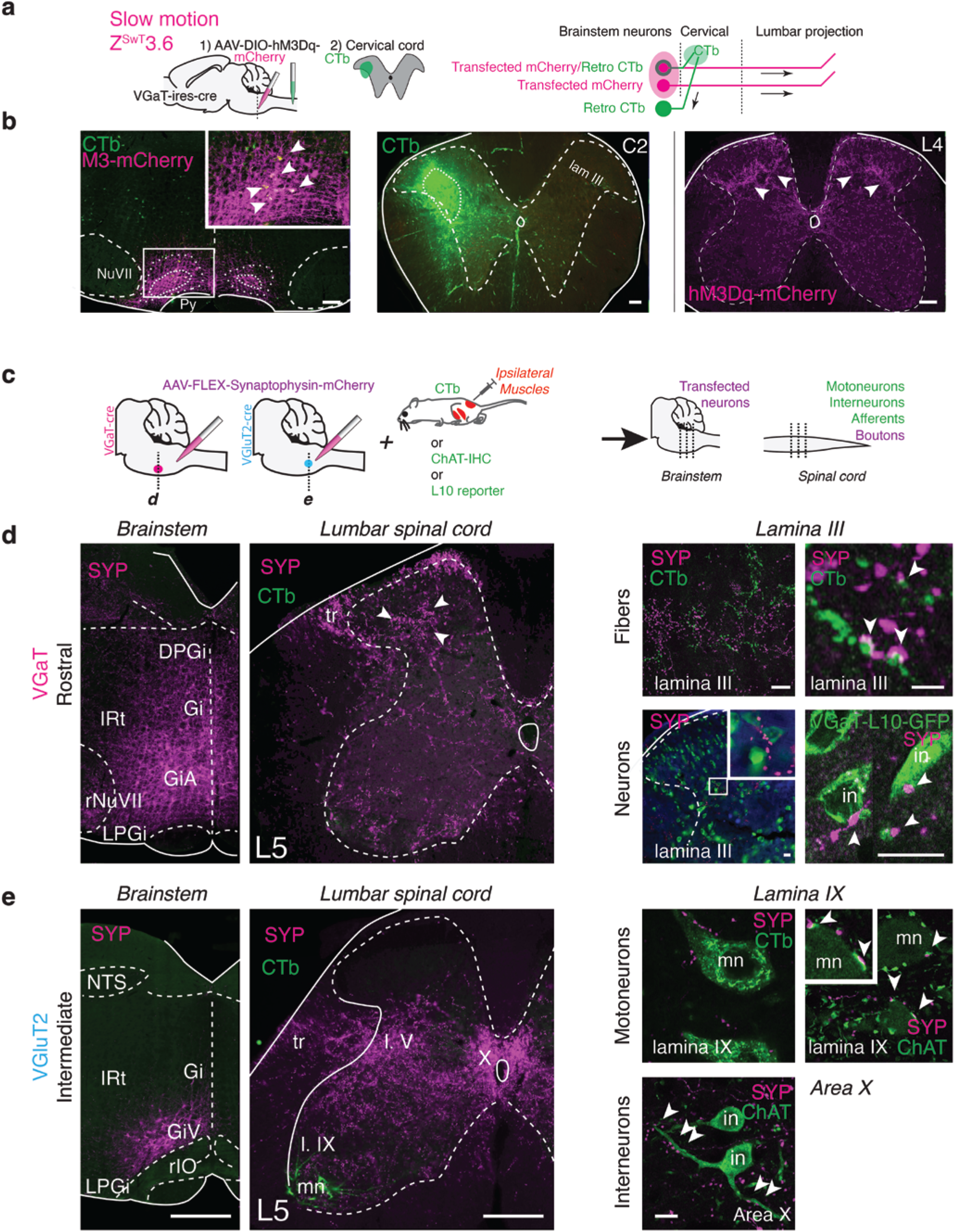
*Slomo* and *shuffle* hotspots preferentially targeted different substrates in the lumbar cord. **a:** Tracing paradigm to confirm connections from VGaT^hM3Dq-mCherry^ transfected neurons in the *slomo* hotspot to the deep dorsal horn (*slomo* mouse with Z^SwT^ 3.6). **b: Left:** Epifluorescence images of a bilateral transfection site (magenta; bar 250µm) and retrogradely labeled neurons following a CTb injection into the deep dorsal horn (green; middle). Arrowheads indicate CTb-labeled transfected neurons. **Right:** Anterograde hM3Dq-mCherry labeling from the brainstem transfection site confirms dense projections to lamina III in the lumbar cord (arrowheads; bar 100µm). See Figure 7-Figure Supplement 1 for validation of VGluT2 and SerT projections. **c:** Conditional synaptophysin (SYP) tracing from VGaT *slomo* and VGluT2 *shuffle* hotspots in combination with labeling of motoneurons, primary afferents and subtypes of interneurons to explore differentiation in target substrates. See Figure 7-Figure supplement 2 for contrasting projections from nearby mRF sites; See Figure 7-Figure supplement 3 and Table S15 for quantification of profiles. **d-e: left panels:** Images of transfection sites (bar= 500µm) and lumbar spinal projections in VGaT-ires-cre (d) and VGluT2-ires-cre (e) mice. Bar=200µm. Arrowheads: lamina III projections. Arrow: area X projections. Green: CTb labeled motoneurons. V: Lamina V. ***d, right panels:*** SYP^+^ labeling among CTb-labeled large diameter primary afferents (top) or VGaT-L10-GFP reporter interneurons (bottom) in lamina III (bar = 10µm). Terminal profiles (magenta) approximate labeled afferents (bar= 1µm) and VGaT-L10-GFP neurons in lamina III (bar=10µm). ***e, right panels:*** In area X (arrowheads, bottom panel) SYP+ boutons (magenta) heavily innervated ChAT^+^ interneurons. In lamina IX, SYP^+^ boutons (magenta) approximate proximal dendrites or somata of CTb labeled motoneurons (mn; left), but ChAT staining revealed that they mainly targeted large ChAT^+^ boutons (C-boutons; right, arrowheads) rather than motoneurons directly. Bar=10µm. rIO: rostral inferior olive; rVII: rostral facial nucleus; L5: 5th lumbar segment; NTS: nucleus of the solitary tract.

As motoneurons were not the primary spinal substrate for *slomo* and *shuffle* hotspots, we pursued a series of independent anterograde mapping experiments to probe alternative spinal target substrates. These studies also served to verify differences in density or distribution between gait style hotspots versus nearby ventral mRF sites (as shown in Figs. 6 and Figure 6-Figure supplement 2). We placed single 5-10nl injections of conditional AAV-synaptophysin(SYP)-mCherry into or adjacent to *slomo* and *shuffle* hotspots in VGaT-ires-cre, VGluT2-ires-cre or respective reporter L10-GFP mice (Fig. 7) and identified motoneurons, large fiber primary afferents, a subset of commissural interneurons, cholinergic interneurons, or cholinergic boutons using a combination of tracing, reporter and IHC tools (Fig. 7; Figure 7-Figure supplements 2 and 3; Table S15 for quantification). SYP^+^ projections to lumbar segments confirmed dense innervation of lamina III from the VGaT^+^ *slomo* brainstem region and of area X from sites involving the VGluT2^+^ *shuffle* site. VGaT SYP^+^ boutons in lamina III derived from the *slomo* hotspot apposed CTb-labeled large fiber primary (muscle) afferents and with L10-GFP^+^ VGaT reporter neurons (Fig. 7). VGluT2 SYP+ boutons in area X derived from the *shuffle* hotspot apposed cholinergic interneurons or L10-GFP^+^ VGluT2 reporter neurons that represented commissural interneurons as they were retrogradely labeled by CTb from the contralateral lumbar cord (Fig. 7; Figure 7-Figure supplement 1). As *shuffle* style changes in swing time were more obvious contralaterally in unilateral DREADD experiments, such a commissural (crossed) system may mediate this component of the *shuffle* phenotype. As for *off* sites, ventral mRF VGaT sites caudally or laterally overlapping the *slomo* hotspot (Figure 7-Figure supplement 1) projected more densely to lamina IX or superficial layers of the dorsal horn. Rostral ventral mRF VGluT2 sites neighboring the *shuffle* hotspot (Figure 7-Figure supplement 1) resulted in denser innervation of lamina IX, whereas more caudal ventral sites mainly innervated lateral laminae V-VII (Figure 7-Figure supplement 1). This latter pattern involving laminae V-VII was found from ventral sites along the rostrocaudal axis of the medullary mRF. Quantification of bouton-like profiles in lamina IX derived from different levels in the ventral mRF using confocal imaging confirmed that more caudal VGaT and more rostral VGluT2 sites more heavily innervated lamina IX than levels representing the centers of *slomo* and *shuffle* sites (Figure 7-Figure supplement 2;Table S15).

Additional features separated projections from hotspots versus neighboring sites. The few boutons in lamina IX derived from the *slomo* hotspot apposed motoneuronal dendrites and had an average diameter of 2.48µm+/-0.42µm. In contrast, ventral VGaT mRF sites lateral or caudal to the *slomo* hotspot densely innervated motoneuron somata with smaller bouton size (1.40µm +/-0.32µm). As for the *shuffle* hotspot, it innervated lamina IX sparsely with appositions onto large cholinergic (C) boutons rather than motoneurons. This in contrast to abundant lamina IX innervation from slightly more rostral mRF VGluT2 sites, with appositions directly onto motoneurons. This differential innervation of motoneurons and cholinergic boutons was confirmed with quantification at the group level (Figure 7-Figure supplement 2; Table S15). Thus, in line with their differential effects on walking style, VGaT *slomo* and VGluT2 *shuffle* hotspots and nearby “*off*” sites differentially projected to specialized spinal substrates, with a limited degree of overlap, supporting a functional-anatomical organization of walking style.

### May sensory gating contribute to prolongation of swing time in *slomo* walking?

The mapping findings of the *slomo* hotspot preferentially innervating the deep dorsal horn raised the question of whether sensory modulation may form the basis for *slomo* style. We probed this by systematically quantifying VGaT^hM3Dq-mCherry^ bouton-like profiles in 6 regions of the lumbar spinal gray matter in Z^SwT^>2 (*slomo*) and Z^SwT^<2 (*non-slomo*) groups using confocal imaging (Fig. 8; Table S16). Bouton counts per standardized volume were higher in *slomo* than *non-slomo* mice only in lamina III. In addition, Z^SwT^ correlated significantly and positively with bouton density in this region, but not in other regions. The correlation trend was negative for lamina IX. These results support the hypothesis that *slomo* style may involve modulation of sensory circuits in the dorsal horn rather than motoneurons in lamina IX. To test this further we assessed sensory responses (Fig. 8; Table S17) in VGaT^hM3Dq^ mice with Z^SwT^>2 and Z^SwT^<2. VGaT^hM3Dq^ activation in Z^SwT^>2 mice reduced the response rate to von Frey filaments that produce mild pressures of 1 and 2g. In contrast, responses to noxious heat (55°C) and mechanical stimuli (pinprick), which are mediated by nociceptors innervating lamina I/II, did not change, further corroborating anatomical data. Together, these data support that *slomo* style changes induced by VGaT^hM3Dq^ activation are not mediated by direct actions on motoneurons, but rather may involve inhibition of low threshold sensory input in the deep dorsal horn and intermediate zone.

**Figure 8:**
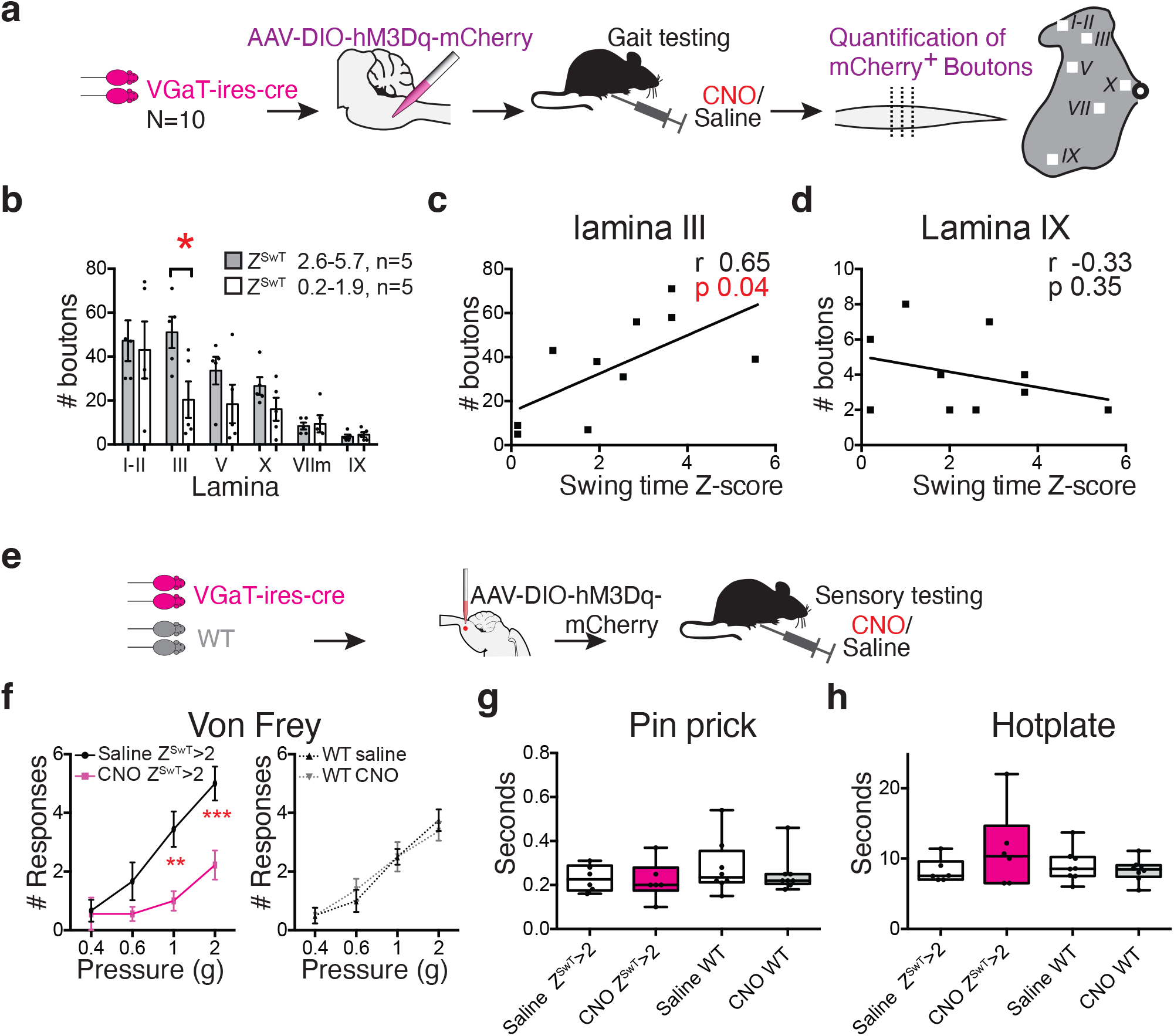
S*lomo* walking style was associated with modulation of low but not high threshold sensory stimuli. **a:** Experimental paradigm for quantification of bouton-like profiles in lamina III versus other regions in the lumbar cord derived from ventral mRF VGaT^hM3Dq-mCherry^ neurons. **b:** Average number of boutons from 3 confocal stacks per mouse in the superficial dorsal horn, lamina III, lateral lamina V, area X, laminae VIII or IX of 5 mice with high Z^SwT^ (*slomo*; gray bars) and 5 mice with low Z^SwT^ (white bars). In lamina III, high and low Z^SwT^ groups differ (p 0.016; two-way ANOVA, followed by Sidak’s multiple comparison test, Table S16; bars represent SD). **c:** In lamina III numbers of mCherry+ boutons correlated significantly and positively with Z^SwT^. Correlations between mCherry+ bouton numbers and Z^SwT^ scores were positive but non-significant for other regions, except for lamina IX **(d)** in which the trend was negative (Pearson correlation, see Table S16). **e:** Experimental paradigm for sensory testing. **f:** Responses of *slomo* VGaT-ires-cre mice with Z^SwT^>2 and WT controls to graded von Frey filaments at baseline (saline) and CNO. *Slomo* mice (Z^SwT^>2) had a significantly decreased response following CNO compared to saline for larger filaments 1 and 2 (bars indicate SD; n=9 with Z^SwT^>2 of 14 mice that entered von Frey testing; 2-way RM-ANOVA; Table S17). **g-h:** Response time to Pinprick and Hotplate tests did not reveal significant differences (n=6 Z^SwT^>2 of 10 mice that entered this testing battery; two-tailed Wilcoxon Rank test; median, 25th and 75th percentile; bars: min and max; Table S17).

### *Slomo* style involves gait phase specific mechanisms

Activation of the *slomo* hotspot shifted walking signatures of swing but not stance time. This dissociation suggests that this site functions in a phase specific way, a phenomenon that is critical for the integration of afferent streams of information into a gait cycle appropriate response. We used an optogenetic approach to test this. We transfected VGaT neurons at and near the *slomo* hotspot with AAV-FLEX-ChR2-mCherry and optically stimulated these sites through a fiber placed above the target site. Stimulation of the transfected *slomo* hotspot for several gait cycles of steady walking (seconds) reproduced *slomo* walking (Fig. 9a-b; Supplementary Table S18; Video S4), with similar shifts in walking signatures between laser-ON and OFF conditions as seen with chemogenetics. Using an EMG-locked trigger, we then delivered one or two 5 ms pulses prior to, during or after the onset of the TA burst during walking (Fig. 9c) and compared TA EMG envelopes averaged over multiple walking cycles with the laser-OFF baseline. Stimulation of the *slomo* hotspot during or < 0.1s prior to TA burst activity prolonged TA bursts compared to laser-OFF baseline, whereas TA burst duration remained unchanged when pulses were delivered more than > 0.1s prior to onset or after termination of the burst (Fig. 9d i). Stimulation of the ventral mRF caudal to the hotspot (paradoxically) increased TA EMG amplitude compared to sham, irrespective of the timing of the pulse (Fig. 9d ii). Temporal or amplitude responses were absent when ChR2 transfection avoided the ventral mRF (Fig. 9d iii). These results confirm phase specificity of the VGaT *slomo* site and support chemogenetic findings that modulation of muscle burst duration and amplitude are independently controlled by nearby mRF substrates.

**Figure 9:**
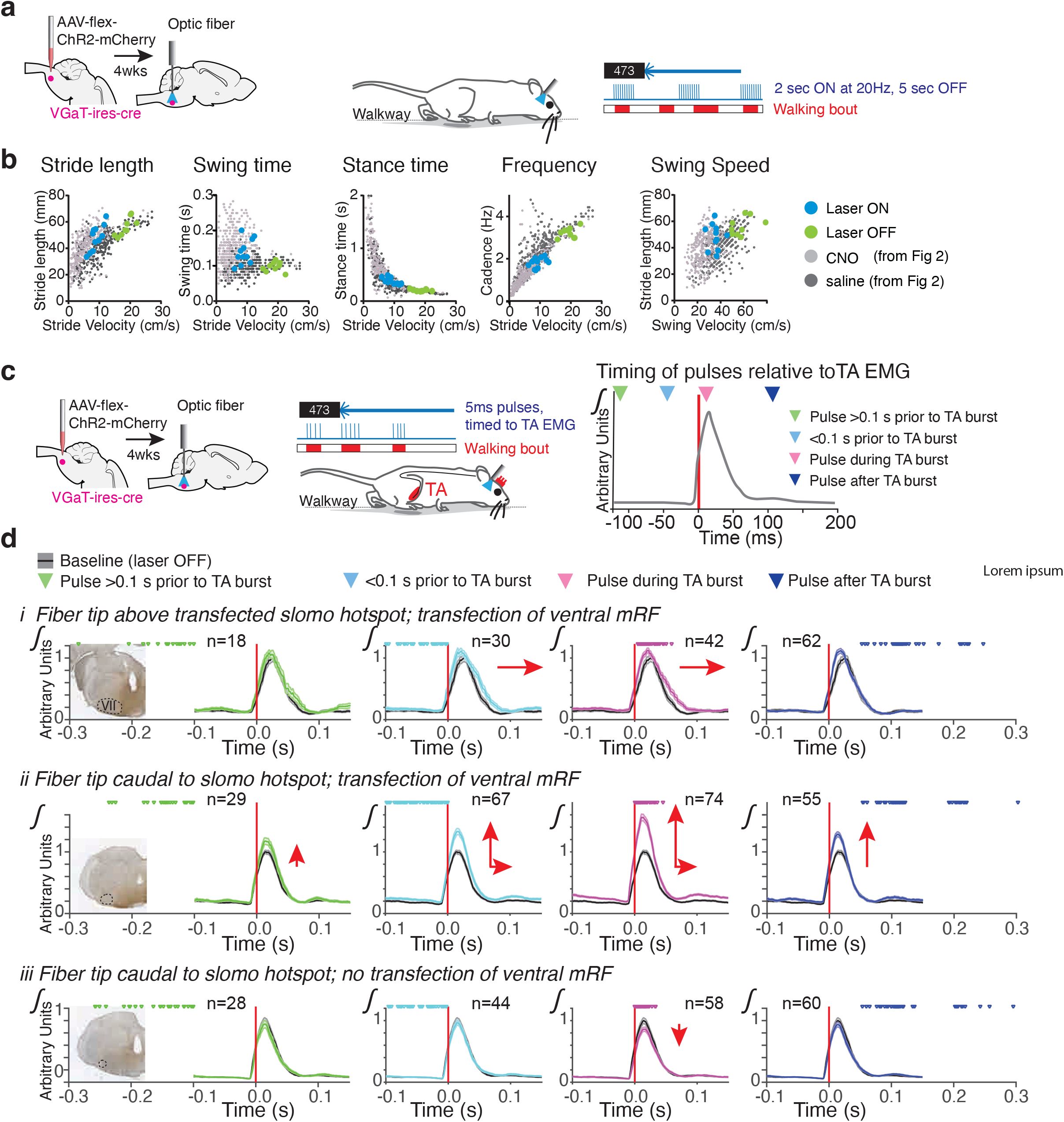
Two seconds of optogenetic stimulation of the *slomo* site reproduced chemogenetic changes in gait style, whereas temporally precise stimulation revealed phase specificity. **a:** In VGaT-ires-cre mice, AAV5-FLEX-ChR2-mCherry was delivered bilaterally to the slomo hotspot and an optic fiber was implanted in the midline above this site. Walking gait was captured on a walkway in laser OFF and ON conditions (wavelength 473nm; frequency 20Hz; pulse width 5ms, train duration 2s). **b:** In the laser OFF state (green), gait metrics overlap with the saline condition in VGaT^hM3Dq^ mice (dark gray). In the laser ON state (blue), metrics overlap with the CNO condition in VGaT^hM3Dq^ mice (light gray). **c:** To study phase specificity during steady walking, single 5ms pulses were delivered at different time points relative to TA burst onset during walking based upon real time TA EMG. **d:** TA-EMG (average +/- SEM; n = number of cycles averaged) without pulses (gray) or following 1-2 5ms pulses >0.1s before (green), <0.1 sec before (light blue), during (pink) and after (dark blue) TA activation. Red arrows mark size and direction of changes in EMG envelope between sham and stimulation parameters. Phase specific responses of TA EMG envelope were found following ventral mRF transfection including the slow motion hotspot. Stimulation just prior or just after TA burst onset delayed the ramp down of the EMG envelope when the hotspot was involved (i), and had marked effect on burst amplitude when more caudal sites were involved (ii). Stimulation of more dorsally transfected sites (iii) did not result in robust changes. Inserts illustrate the fiber tip placement (hole) in relation to the ChR2-mCherry transfected area. Table S18 for statistics.

## Discussion

We showed that slow motion-like (*slomo*) and shuffle-like (*shuffle*) styles represent walking styles that are distinct from each other and from normal slowed walking. Walking signatures served to classify and quantify key components of these contrasting styles and to distinguish these slowed styles from regular slowed walking. These styles mapped to discrete inhibitory and excitatory hotspots in the ventral brainstem (Graphical Abstract). In turn, these hotspots preferentially innervated contrasting substrates in the spinal cord. Activation of brainstem sites outside the style hotspots, irrespective of cell type, modulated walking speed alone, without style changes. These results advance concepts related to the rich but poorly understood repertoire of adaptive variations that color walking gait and of maladaptive variations that define gait disorders (26).

### The unique design served to elicit, quantify and map walking styles

The conceptual and technical design of this study stood out from prior work. We enhanced translational potential by focusing on walking gait, which is shared among bi- and quadripeds, rather than speed capacity (20) or transitions between gait patterns that are unique to quadrupeds (22). Secondly, while treadmill can be used to control speed (23) this would have impeded self-selection of speed, crucial for this study. Furthermore, we did not optimize our assay to halt mice (18), as that would have defied the purpose of our study. Additionally, canonical activation of the hM3Dq receptor magnifies physiological responses and differs from direct depolarization by optogenetic activation (44, 45) that was used in many prior mouse studies (18, 20, 23). Next, we combined the high specificity of discrete hM3Dq modulation sites in eliciting walking styles with a sensitive walking signature toolkit into an unbiased assay to map out a functional sub-organization, a common feature of supraspinal systems (46, 47), but beyond prior locomotor studies that employed other methodologies (18, 20, 23). Finally, hindlimbs provide the propulsive component of walking in both bi- and quadruped models and therefore we focused on mRF sites with connections to the lumbar cord, whether with or without connections to other CNS sites, while avoiding sites exclusively serving forelimb goal directed movements (48) and lateral sites serving respiration and autonomic functions (25, 49).

### Discrete mRF sites serve distinct walking styles

The degree of functional-anatomical organization within the mRF remains controversial. We revealed a more precise functional-anatomical organization within the mRF than reported in studies using larger transfection sites, other functional tools, and anatomy-based site selection (18, 20, 48, 49). Both *slomo* and *shuffle* hotspots reside within the ventral mRF, but their centers were rostrocaudally distinct per stereotaxic coordinates and mapping. Hotspots were not anatomically demarcated and they extended beyond their respective centers. Each occupied only a fraction of the gigantocellular reticular formation, the *shuffle* site rostrally in the ventral sub-nucleus and the *slomo* site at the rostral tip of the alpha sub-nucleus. These findings do not implicate that all mRF populations adhere to such an organization or that these hotspots only serve walking style. For example, a rostrocaudally extensive ventral mRF excitatory territory innervated lateral spinal laminae V-VII as documented before (18, 48, 49) and intermixed with and expanded well beyond the hotspots towards the cervico-medullary junction.

Slowing of walking speed *without* style or signature modification was not cell-type and location specific (18, 20, 23, 24). In contrast to efferents from hotspots, projections from “off” sites reached lumbar spinal sites with variable patterns and density while ascending projections from dorsal “off” sites were robust, particularly for the inhibitory system. Our study did not probe whether the heterogeneous “off” sites primarily serve other motor and non-motor functions (50, 51) or modulate local interneurons that promote non-specific reticulospinal activation (52). Much remains unknown about the interplay between descending, local and ascending mRF circuits in integrating centrally-driven motor commands and conveying a unified response to spinal circuits with supraspinal feedforward and feedback mechanisms serving well-integrated motor function.

### Walking style modules represent components of more complex mobility repertoires

Focally elicited *slomo* and *shuffle* styles only represent single components of complex adaptations. For example, slow-motion-like movement is a feature of stalking gait. However, crouching posture can but does not necessarily accompany stalking gait. The finding that lowered posture elicited by mRF modulation did not map specifically to the *slomo* site supports this notion. Similarly, mice that walked with EMG features of halting steps following VGluT2 activation completed walking and other tests, implicating that substrates that mediate halting steps are dissociated from substrates that mediate the intent to stop. Such compartmentalization provides flexibility in movement repertoires by allowing different combinations of circuits (18–23, 32, 53) to work in concert as necessary to meet exploratory, appetitive, and defensive needs, integrate postural adjustments and goal directed movements and fits with classical concepts of supraspinal motor control (14, 54, 55).

Activation of walking style hotspots had only minor effects on swimming, a different locomotor mode. The shorter cycle duration during swimming, despite higher forward resistance, may not leave sufficient time for (multisynaptic) modulatory actions (14).

Alternatively, robust swimming-associated changes in circuit function may bypass or override modulatory actions (12, 56) or the hotspots may preferentially serve slower spinal locomotor speed modules (12).

A feature of *slomo* gait but not of postural changes was phase-specificity, a mechanism that allows circuits to mediate phase specific commands, as during walking, even when afferents provide temporally less precise or tonic activation, or, as in this study, during chemogenetic activation. Shifts in swing time but not stance time signatures suggested phase-specificity in *Slomo* gait. Phase-specificity is an intrinsic characteristic of sub-sets of reticulospinal neurons shown with electrophysiology (30, 57, 58) and optogenetics of excitatory (23) but not inhibitory mRF systems. EMG-locked optogenetic pulses to activate the VGaT *slomo* hotspot in temporally precise manner confirmed this phenomenon for the *slomo* hotspot: prolongation of swing time and anterior tibial EMG burst duration (key *slomo* features) occurred mainly when activation was administered during the swing phase. Caudal ventral VGaT sites that modulated EMG interburst amplitude (i.e. tone) did not display the same degree of phase specificity.

### Ideas and speculation

### Are different walking styles driven by unique downstream neural substrates?

In line with their distinct effects on walking style, connections from *slomo* and *shuffle* hotspots to the lumbar spinal cord differed from each other and from nearby sites. Unexpectedly, VGaT *slomo* and VGluT2 *shuffle* hotspots only weakly innervated motoneurons, in contrast to VGaT neurons just caudal to the *slomo* hotspot and VGluT2 neurons just rostral to the *shuffle* hotspot. In hindsight this is less surprising, as these styles differentially affected swing time, stance time and/or stride length signatures. As baseline signatures for these 3 metrics behave differently (Fig.1), it follows that stance time, swing time and stride length are mediated by different neural substrates. In line with this, hotspots that differentially modulated these 3 metrics preferably projected to distinct spinal substrates. These data provide concrete leads for further hypothesis development and testing.

#### Slomo walking

Activation of the VGaT *slomo* hotspot changed walking style while dampening the response to low threshold sensory stimuli. The *slomo* site projected among large fiber primary afferents and neurons in the deep dorsal horn, rather than ventral horn, suggesting that sensory modulation may form the basis for slow-motion movements. While surprising, there is a rich literature on the sensory control of spinal locomotor circuits as reviewed by Rossignol et al., (14). Modulation of proprioceptor and joint receptor input during walking is known to reduce proprioceptive reflexes, while modulation of cutaneous transmission limit sudden changes to the step cycle (59). Such changes in gain of sensory motor circuits may lead to fewer errors in locomotion (60). Indeed, we found that locomotor patterns, hindlimb kinematics and motor tests did not degrade in *slomo* walking, in contradistinction to the absence of proprioceptive feedback (61) or following dorsal horn interneuron ablation (62). Further studies will be necessary to confirm such a modulatory role of pre- and/or postsynaptic inhibition of neurochemically and molecularly diverse (63) lamina III interneuron subtypes and classes of sensory afferents.

#### Shuffle walking

Descriptive and quantitative approaches demonstrated that the VGluT2 *shuffle* hotspot heavily targeted area X, including cholinergic interneurons known to give rise to so-called C-boutons that terminate onto motoneurons. We found that this hotspot also innervated C-boutons themselves. This intrinsic cholinergic spinal system mediates an increase in motoneuron firing during locomotor-like activity by modulating afterhyperpolarization (64). Thus, through both pre- and post-synaptic mechanisms, the VGluT2 *shuffle* hotspot may enhance the actions of this modulatory cholinergic system. This may underlie the disproportionally shortened stride length of *shuffle* walking through the co-activation of agonist-antagonist hindlimb muscles detected via EMG. A VGaT site caudally overlapping with the *slomo* hotspot targeted this same cholinergic substrate (Fig. S8), raising the question whether these VGlut2 and VGaT sub-systems may exert interdependent control of this motor feature analogous to the respiratory system (65).

In addition we found that the VGluT2 *shuffle* hotspot projected ipsilaterally to excitatory commissural interneurons (neurons that project to the opposite side of the spinal cord) in area X. These connections may explain that *shuffle* swing time signature shifts were more robust in limbs *contra*lateral to unilateral transfection sites. Dedicated studies will need to confirm the role of these spinal targets in *shuffle* gait and reveal additional targets.

#### Faster walking

Activation of sets of medullary SerT neurons differentially modulated speed, but without shifts in walking signatures. Activation of rostral sites, i.e. raphe magnus, decreased walking speed similar to activation of “off site” VGaT and VGluT2 mRF sites. The more caudal raphe obscurus, on the other hand, represented the only site in this study from which activation increased walking speed. Serotonergic raphe obscurus neurons innervate spinal motoneurons (66), where serotonin mediates facilitation of voltage-sensitive changes. This results in increased input–output excitability, i.e. gain (59), with serotonergic activity increasing with increased motor output (67). The effect size of serotonergic activation was modest, which may reflect molecular, anatomical and functional heterogeneity of medullary serotonergic neurons (68). Alternatively, more robust increases in gait velocity may require interdependent control of mRF-SerT and other reticulospinal systems. We did not focus on lateral paragigantocellular SerT neurons, which also project to motoneurons, as that site co-expresses GABA and/or VGluT3 (69) which would have confounded interpretation.

### How may these insights translate to human walking?

Decreased walking speed is a cardinal marker of system failure in neurological conditions, aging, and systemic diseases (4, 70–72), but speed alone is too non-specific to disentangle underlying pathophysiology. Objective quantification while disentangling CNS circuit nodes mediating the various changes has remained challenging. As human walking adheres to signatures as it does in mice, as walking signatures shift in persons with CNS disease (36, 38), and as caudal brainstem structures are well preserved among species, brainstem sites homologous to those defined in this study are likely important for human gait control. As such, *slomo* and *shuffle* features may serve a repertoire of adaptations in a context dependent way. This includes stalking/cautious gait (*slomo*) and vital real-time corrections of single strides through phase-specific temporal control (both styles). Is the stimulatory modulation applied in this study relevant for pathologic mechanisms? While it obviously does not represent loss of function at the sites of interest, it may well be representative for imbalances due to dysfunction of connected circuits. This is a known principle in neurodegenerative diseases, analogous to disinhibition of excitatory subthalamic neurons due to nigral pathology in parkinsonism. Signature shifts seen in *shuffle* mice manifested in a subset of persons with Parkinson disease (36, 38), whereas *slomo* signature shifts characterized the final stage of aging in wild type mice (39), months after a decrease in walking speed alone. Such concepts of maladaptive modulation of walking style can now be tested through collection of gait signatures in human subjects in combination with structural and functional imaging tools, using sites homologous to those unveiled in this study as an anchor.

## Materials and Methods

### Ethics statement

Handling and housing of animals, surgical procedures, post-operative monitoring, behavioral tests and euthanasia were performed in strict accordance with the Guide for the Care and Use of Laboratory Animals of the National Institutes of Health at the animal research facility of the Center for Life Sciences. The Institutional Animal Care and Use Committee at Beth Israel Deaconess Medical Center reviewed and approved the experimental protocols (#069-2022). Survival surgeries were performed under isoflurane anesthesia and under aseptic conditions. The analgesic agent Meloxicam was given prior to surgery and on the first postoperative day.

### Viral vectors and tracers

All viral vectors have been described and used before:pAAV8-hSyn-DIO-hM3D(Gq)-mCherry (Bryan Roth (45); UNC Vector core; RRID: Addgene_44361), AAV2/5-hSyn-hChR2(H134R)-EYFP or AAV2/5-hSyn-hChR2(H134R)-mCherry(1.3×10^12^; Stanford Gene Vector and Virus Core; K. Deisseroth), AAV8-EF1a-DIO-synaptophysin-mCherry (MGH Vector core; gift by Dr. B. Lowell, BIDMC). Cholera Toxin subunit B (CTb; List Biological) was used as conventional tracer. For AAV injections into the brainstem volumes ranged from 5-100nl (median 35nl). For spinal cord and muscle CTb injections, volumes ranged from 25-150nl (median 100nl; 1% in distilled water), and 0.5-1µl (median 0.8µl; 0.1% in distilled water), respectively.

### DREADD activation experiments

#### Animals

To study effects of chemogenetic activation of excitatory, inhibitory and serotonergic neurons in the medial pontomedullary reticular formation (mRF) on walking we used 35 male VGaT-ires-cre knock-in mice and 10 WT litter mate controls, C57Bl6/J background; RRID: IMSR_JAX#: 016962 (34), 46 VGluT2-ires-cre and 7 WT litter mate controls, C57Bl6/J background; RRID: IMSR_JAX#: 016963 (34), and 18 transgenic SERT-cre mice and 15 litter mate controls: (Tg(Slc6a4-cre)ET33Gsat; RRID: MMRRC_031028-UCD (35), both sexes, C57Bl6/J background. Mice were 8-16 weeks at the time of AAV injections and 12-20 weeks at the time of first behavioral testing. Mice in each experimental cohort served as their own control and were randomly assigned to a set of mRF target sites. We worked with 5-15 mice at the time, consisting of a mix of the various knock-in/transgenic mice and WT litter mates and with different target sites, to facilitate blinding of testers and raters for genotype and site. This was repeated until group size for each of the sub-cohorts was established, while also providing replication throughout the study. Following completion of behavioral testing, sub-cohorts were then randomly assigned to further EMG implantation, additional circuit mapping experiments, or in case of VGaT mice, sensory testing. One mouse was excluded from analyses as the transfection site was located far outside the area of interest.

#### Group size

Based upon an average walking speed of 11.2 +/- 4.5 cm/s (saline) and 6.6 +/- 2.9 cm/s (CNO) for VGaT, and 11.5 +/- 3.4 cm/s (saline) and 7.9 +/- 2.7 cm/s (CNO) for VGluT2 mice in pilot experiments, we calculated a group size of 5 (VGaT) to 7 (VGluT2) mice to detect a difference in gait metrics with a power of 90% and an alpha level of 0.05. We used a similar group size for the SerT cohort to keep approaches comparable. To map out hotspots for walking styles, we expanded these numbers to account for inclusion of positive, mixed and negative sites and to account for laterality of transfection sites.

#### Placement and size of AAV injections

Injections were placed as described before (25, 49, 73). In short, we injected the medulla using an approach through the foramen magnum with the head at a 90 degree angle to avoid a needle tract through the cerebellum and to allow for direct guidance by nearby landmarks (Kopf stereotact; relative to the Obex: AP +0.25 to −0.45mm; L: 0.1-0.4mm; DV 1.9-3.2mm). Coordinates of hotspots, as deducted following completion of the study, were: slow motion-site: AP 0, L +/-0.3, DV −2.7; shuffle site: AP −0.2, L +/-0.25, DV −2.8; raphe obscurus: AP −0.45, L0.0, DV 2.9. The corresponding coordinates for a standard approach using Bregma are: slow motion site: AP −5.8, D −5.4, L 0.3; shuffle site: AP −6.3, D −5.1, L 0.25; obscurus: AP −6.9, DV 5.5, L 0.0. AAV-hsyn-DIO-hM3Dq-mCherry injections were aimed to cover the large population of reticulospinal neurons that reside in the dorsal and ventral regions of the reticular formation at the levels of the rostral medulla oblongata and pontomedullary junction (25). Individual injections were kept discrete (median 35nl), as large injection sizes (100nl) in pilot experiments did not produce more robust phenotypes and lacked the spatial resolution necessary to identify hotspots. Pilot experiments suggested that bilateral activation was necessary to induce VGaT mediated gait phenotypes, whereas unilateral activation appeared sufficient for the VGluT2-ires-cre group. Therefore we placed the majority of injections bilaterally in the VGaT-ires-cre cohort, and added a unilateral group that was aimed at the slomo hotspot to verify that unilateral injections were not sufficient to elicit a phenotype. For the VGluT2-ires-cre group, we placed the majority of the injections unilaterally, and added a smaller bilateral cohort for verification.

#### CNO dosing and timing

To assess the appropriate dose range and time window for the experiments we obtained CNO dose response data in 14 VGaT-ires-cre with Z^SwT^>2 (see below) and 20 VGluT2-ires-cre mice with Z^SwT^<-2 (doses of 0.15mg/kg, 0.3mg/kg i.p and/or 0.6mg/kg i.p) and time response data in 5 VGaT-ires-cre mice with Z^SwT^>2 (at 0.5 – 6 hours following a single 0.3mg/kg CNO dose; Supplementary Figure 1; Supplementary Table 1). A dose of 0.3mg/kg CNO produced robust effects without side effects and without further increase in effect size at higher doses. The optimal testing window ranged from 45 minutes to 2.5 hours after i.p. injection and this window was applied throughout the study. Changes in swing time and stride velocity did not augment further at 0.6mg/kg dose.

#### Behavioral assessments

Following AAV-hsyn-hM3Dq-mCherry injections, mice were habituated to all behavioral testing conditions for at least 4 sessions. We used cohorts of 5-15 mice, consisting of a mix of genotypes and target sites to limit bias by testers and raters. Testing started at least 4 weeks following AAV injections to assure stable transfection levels. Tests were performed during the first half of the light-ON period on non-consecutive days to prevent sleep deprivation, and included a minimum of 2 saline and 2 CNO session to allow for replication within each mouse. Gait analysis was the primary test, whereas balance beam, horizontal ladder, and rotarod testing were performed to assess whether modulatory effects were gait specific. All tests were performed in the same order (rotarod, gait, beam, ladder). A sensory screen was added to the testing battery of the VGaT cohort, based upon projections from mRF^VGaT^ to the spinal dorsal horn. This screen was performed on a separate day.

a. *Video gait assessment:* Using high speed video, we obtained spatial and temporal gait parameters from hindlimb and forelimbs as described and validated before (36, 38). Briefly, video frames were captured at 120fps from the middle 40cm of a 120cmx8cm plexiglass walkway (74). Foot print data was visualized via a mirror, placed underneath at an angle of 45 degrees, aided by side view (Fig 2a). At least 4 trials were recorded per mouse per session or more as needed to obtain at least 4 valid trials. A valid trial contained at least 4 consecutive strides per limb of continuous walking. Spatial (location of the footfalls) and temporal data (marking of stance and swing phases) was extracted from video frames using a custom MATLAB code and was used to obtain the following parameters: stride velocity, swing and stance time, stride length, coupling of limbs, cadence, and swing speed (36). The average number of hindlimb strides per animal per session is 36, 35 and 38 in the VGaT, VGluT2, and SerT groups at baseline, respectively, and 36, 42, and 40 in the CNO condition, respectively. Total number of strides in each group is indicated in the corresponding Figures.
b. *Balance beam:* We assessed balance with the horizontal beam test. We recorded mice with a conventional video camera while they walked across a 6mm wide, 120cm long beam (75). We quantified numbers of slips and misses of the hind paws from these recordings and calculated average performance of 2 trials per session in each mouse.
c. *Rotarod*: We used a paradigm designed to test performance of learnt motor behavior. Mice were trained on an accelerating rotarod (Ugo Basile) prior to baseline testing for at least 4 sessions on non-consecutive days. A session consisted of 3-5 trials, starting at 2 rpm and increasing to 75rpm over 4 minutes. Time to fail, either by falling or clamping for 2 consecutive rotations, was recorded. We calculated average performance of 3 trials per session in each mouse.
d. *Horizontal ladder:* This test assesses motor planning. Mice were recorded with a conventional video camera when traversing a 8cm wide, 75cm long horizontal ladder with metal rungs at irregular interspaces (76). We counted numbers of slips and misses of the hind paws and calculated average performance of 2 trials per session in each mouse.
e. *Kinematic data:* To obtain kinematic data from representative gait cycles at walking speed were marked base of the tail, iliac crest, knee, ankle, 5^th^ metatarsal, and tip of 4^th^ digit using the same frame by frame footage as for video gait analysis (Adobe Photoshop and Matlab).
f. *Sensory screen:* We used i) Von Frey, ii) pin, and iii) hotplate tests as described before (77) in sub sets of VGaT-ires-cre mice and WT littermates.

i. To test the response to static punctate mechanical stimuli we used von Frey testing. Mice were placed on a table with 0.5 x0.5cm stainless steel mesh, confined by a 9cm x 12.4cm glass cylinder, and their response to mechanical stimuli was determined using a series of von Frey filaments that produced a bending forces of 0.4 to 2g. Stimuli were applied to the sciatic nerve region of the plantar surface of the foot for 1-3 s with intervals of at least 5 s after mice returned to their resting state. Each filament was applied 10 times starting with the smallest force.
ii. The pinprick test assesses response to high threshold mechanical stimuli. Mice were placed on the mesh table as above and the plantar surface of the hindpaw was touched with an insect pin (Fine Science Tools, Black Anodized Steel, size 000). Application lasted less than 1 s and was performed 3 times per trial. We recorded the average time spent flinching or licking the paw with a stopwatch for 2 trials.
iii. The hotplate test determines the response to contact heat pain. Mice were placed on a plate heated at 55 C, confounded by an acrylic container (Bioseb, France). Latency to flinching, licking or jumping was measured. Habituation took place at 30 C and baseline measurements served to reduce the risk of habituation of sensitization. Only one reading was performed per day and this test was always performed last.
g. *EMG:* Following completion of behavioral testing, in a subset of mice we placed a maximum of 3 pairs of bipolar EMG electrodes in 3 sets of muscles as guided by the phenotype. This included anterior tibial (TA), gastrocnemius (G), gluteal, quadriceps, or triceps brachii muscles (78). EMG electrodes were custom made from multistranded Teflon coated stainless steel wires (A-M systems, Washington; Cat #793200) that have little effect on kinematics (74). Electrodes were tunneled underneath the skin and attached to a microconnector (Hirose Electric) which was affixed to the skull with dental cement.

We recorded EMG during quiet Rest, Walking, and Swimming using home cage, walkway and swimming pool, respectively, to study muscle activation patterns. Recordings took place between 5-10 days after implantation. A lightweight cable was attached to the connector on the mouse’s head. EMG was collected at a sampling rate of 10,000Hz, signals were amplified (Grass Technologies, Model 15LT Physiodata Amplifier System), digitized (Grass Polyview software or Cambridge Electronic Design Spike2 software with Micro 1401-3 data acquisition unit) and stored for analysis.

### Mapping experiments

We performed 3 *circuit mapping studies* to study efferent projections and probe targets from different sites and cell types in the mRF, guided by functional mapping of DREADD transfection sites using 12 VGaT-ires-cre, 12 VGluT2-ires-cre, 6 SerT-cre mice (see DREADD experiments for source details; C57Bl6/J background; both sexes; age 8-16 weeks at the start of the study) and 8 WT mice (C57Bl6/J or mixed C57Bl6/J:129P3/J background, both sexes; RRID: IMSR_JAX#: 000690; RRID: IMSR_JAX#: 000664; age 8-16 weeks at the start of the study). A group size of 3-4 mice was adequate to ensure reproducibility. Injections into the brainstem, spinal cord or muscles were placed as described before(25, 49). Calibrated target volumes were delivered through a glass pipette using air pressure (tip diameter 25-35µm).

a. CTb was injected unilaterally into the cervical (n=4) or lumbar (n=4) spinal cord in 8 WT mice to retrogradely label reticulospinal neurons and determine their neurochemical subtype in the gait hotspots using *in situ* hybridization (see below).
b. Following completion of DREADD activation behavioral experiments, in 6 VGaT-ires-cre, 6 VGluT2-ires-cre, and 6 SERT-cre mice we injected CTb tracer into sub-regions of the spinal cord to retrogradely label hM3Dq-mCherry transfected brainstem neurons. In separate animals we injected limb muscles (see above) to identify spinal motoneurons innervated by hM3Dq-mCherry neurons.
c. We injected 5-10nl AAV8-EF1a-DIO-synaptophysin-mCherry into discrete mRF sites of 6 VGaT-ires-cre and 6 VGluT2-ires-cre mice to anterogradely label boutons in the spinal cord derived from and neurons in the mRF^VGaT^ or mRF^VGluT2^ neurons. We combined this with CTb tracer injections into gluteal, gastrocnemius or anterior tibial muscles to identify spinal motoneurons and large diameter muscle afferents, or with CTb injections into the contralateral lumbar cord to identify commissural interneurons.

### Optogenetic activation experiments

To assess whether mRF-mediated changes in walking and associated EMG characteristics were gait phase specific in the VGaT cohort, as suggested by the chemogenetic gait study, we performed optogenetic experiments. In 10 VGaT-ires-cre mice (for details see DREADD section; both sexes, C57Bl6/J background; age 10-16 weeks at the time of injection), we injected AAV2-DIO-ChR2-mCherry into the mRF using a posterior approach through the foramen magnum, and 2-4 weeks later implanted a single optic fiber (diameter 200µm; Zirconia ferrule, Kientec) above the target area using standard stereotactic approach (Bregma: AP −5.8, D −4.9, L 0) and near the midline to allow for bilateral stimulation. In one paradigm, transfected VGaT neurons were stimulated at 20Hz, with 5ms pulse width, and a train duration of 2-5 seconds at 30-180sec intervals, while video gait analysis was performed (N=4). In a second paradigm (N=6), optogenetic instrumentation was combined with EMG implantation (see above) into left and right TA and left G muscles. Single optogenetic pulses (pulse width 5ms) or double pulses (two 5ms pulses 20ms apart) were then delivered at different time points in the gait cycle based upon real-time TA-EMG feedback (Cambridge Electronic Design, Spike2, custom stimulus sequencer script). Video gait data and/or EMG data was collected for analysis as in DREADD experiments.

### Tissue processing and reagents

After a survival time of 3-14 days (conventional tracing only), 28-45 days (conditional tracing only) or 42-180 days (behavioral cohort), animals were anaesthetized with an overdose of ketamine/xylazine or chloralhydrate (i.p.) and transcardially perfused. After removal of the CNS, tissue was cryo-protected in 20% sucrose in PBS-azide (4°C) and cut into 4 series of consecutive, 40µm transverse sections using a sliding microtome. Free floating sections were stored in PBS-azide or RNAlater (Thermo Fisher Scientific; Cat # AM 7021). To visualize GFP or mCherry labeled neurons at brainstem transfection sites and axons derived from these neurons in the spinal cord, c-fos activation patterns, the tracer CTb, or ChAT, VGluT2, VGaT, SerT or TPH, tissue series were incubated in rabbit anti-GFP (Invitrogen, A11122; 1:20,000; RRID: AB_221569), rabbit anti-dsred (Clontech, 632496;1:10,000; RRID:AB_10013483), choline acetyltransferase (goat anti-ChAT, Millipore, AB144P; lot NG1780580;1:200 in PBT; RRID:AB_11214092), goat anti-CTb (1:3-15,000; List Biological, 703; RRID:10013220), rabbit anti-c-fos (ABE457; lot 2672548; 1:20,000; RRID:AB_2631318), rabbit anti-VGaT (Millipore AB5062P, lot 2482123, 1:1,000; RRID:AB_2301998), guinea pig anti-VGluT2 (Chemicon AB2251-I, lot 2894024, 1:5,000; RRID:AB_2665454), rabbit anti-SerT (Immunostar 24330; 1:1,000; RRID: AB_572209), or goat anti-TPH (Immunostar 20079; 1:4,000; RRID: AB_572262) followed by biotinylated secondary antibodies raised in donkey (1:500; Jackson Immuno Research Laboratories; 1 h; RRID:AB_2340593, AB_2340375, AB_2340451), and avidin-biotin-complex-peroxidase solution (ABC; 1:500 in PBT; Vector Elite Kit, Vector Laboratories, Burlingame, CA; 1-2 h; RRID:AB_2336819). A brown reaction product was obtained following incubation in 0.04% Diaminobenzidine (DAB; Sigma) and 0.012 % H_2_O_2_ in PBS, or a black reaction product by adding 0.2% nickel ammonium sulfate to the DAB solution. For fluorescent staining, we used Alexa 488 (Molecular Probes RRID:AB_141708 or Invitrogen RRID:AB_2535792), −568 (Invitrogen RRID:AB_2534017) or -Cy5 (Jackson ImmunoResearch Labs, RRID:AB_2340415) tagged secondary antibodies (1:500, raised in donkey), or appropriate biotinylated secondary antibodies followed by 1:100 streptavidin-488, -568, -Cy5, or -Pacific Blue (Molecular Probes, RRID:AB_2315383). Selected brainstem sections were processed for both in-situ hybridization (ISH) to visualize VGaT-mRNA or VGluT2-mRNA in combination with IHC to visualize CTb or mCherry (see above), as described previously (34, 79–81) for free-floating conditions.

### Behavioral and electrophysiological data analyses

Behavioral tests were scored blinded to the genotype, experimental condition and injection site, with the limitation that raters were able to recognize some of the phenotypes. We used Graphpad Prism 7 software (San Diego, CA, USA) or custom written MATLAB scripts for statistical analysis. Family wise significance was set at 0.05, except for velocity dependent gait analysis, where it was set at 0.001 (36).

*a) Group definition based upon gait phenomenology:* To define subgroups that represented mice with similar direction and degree of gait changes, we calculated gait metric Z scores based upon each animal’s difference in performance between baseline and CNO, with each animal serving as its own control. This approach is conservative and limits random effects of mouse line, size and age. Scores were based on swing duration in the VGaT and VGluT2 cohorts:

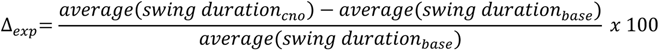

Differences between conditions in the corresponding WT groups defined the population averages, and Z-scores of the change in gait metrics were then calculated as follows:

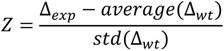

In the SerT cohort, we used speed, rather than swing time as the primary metric to define subgroups as there were no obvious swing time changes in this group. In addition, we calculated Z-scores for each of the other motor tests to determine whether changes in gait and other motor tests correlated and came from the same manipulation site.

*b) Video gait analysis:* We used a validated approach (36, 38) to answer the question whether there is evidence that the CNO treatment shifted gait curves or signatures from saline baseline in the same mice. In other words: do the best-fit values of unshared baseline and CNO gait parameters differ between data sets? For each stride we measured swing and stance durations based on the number of video frames and stride length based upon spatial coordinates using MATLAB. We then visualized these metrics as a function of the velocity of the stride. We derived cadence (1/cycle duration; cycle duration being the sum of stance and swing durations) from these measures, which we then also plotted against stride velocity. We also plotted stride length against swing speed as relevant for the swing phase. This data was then fitted to a curve using the simplest best fit model that was established for the respective gait metric within the speed bin that covers walking and diagonal trotting (3-16cm/s (36)), i.e. establishes the walking signature. We then used a sum of squares F test to assess whether regression curves summarizing the baseline or endpoint datasets were shared. This model takes into account the entire curve, not only intercept or slope. As the comparisons involve the same mice under saline and CNO conditions, other corrections are not necessary. We contrasted these results with an analysis of averaged data sets using 2 tailed t tests, to provide a framework of comparison, as this latter method is widely used even though it is flawed when speeds differ between data sets, and to provide additional insight in animal to animal variability. Stride to stride data on interlimb temporal and spatial coupling was represented in polar plots and analyzed using circular statistics including Rayleigh to test for circular unimodality and Watson-Williams t-test to compare baseline and endpoint datasets (38, 82) for the relevant speed range of 3-16cm/sec.

*c) Kinematic data:* In representative mice, we constructed stick figures based on joint markers from video frames (120fps).

*e) Rotarod*: Data was normally distributed (D’Agostino omnibus K2 normality test). We used a Wilcoxon matched pairs signed rank test compare saline versus CNO performance.

*f) Horizontal ladder, balance beam and grip strength:* Data sets did not all pass the D’Agostino omnibus K2 normality test. Baseline and CNO performance were compared with a Wilcoxon matched pairs signed rank test.

*g) Sensory screen*: pin and hot plate datasets in mice with Z^SwT^>2 and WT littermates were analyzed using two tailed Wilcoxon Rank test, whereas von Frey data was analyzed with 2 way RM-ANOVA followed by Sidak’s multiple comparison test.

*d) EMG analysis in hM3Dq transfected mice:* Of the 26 VGaT-ires-cre, 14 VGluT2-ires-cre, 9 SerT-cre and 6 VGaT WT control, and 7 SerT WT control mice with TA and G EMG implants, we excluded 2 VGaT, 1 VGluT2, 1 SerT, and 2 VGaT WT control with insufficient signal quality. 8 VGaT mice were used for qualitative purposes only, as EMG was placed in other muscles, (i.e. forelimb or other hindlimb muscles). Remaining animals were grouped based upon gait Z-scores as follows: n=6 VGaT Z>2, n=10 VGaT Z<2, n=7 VGluT2 Z<-2, n=6 VGluT2 Z>-2, n=5 SerT Z>0, n=3 SerT Z<0, n=4 VGaT WT controls, and n=7 SerT WT controls. We analyzed data from 20 cycles/mouse during walking and swimming conditions at baseline and during CNO activation (i.e. total of n=80 cycles/mouse). Signal was filtered (IIR 7th order Butterworth filter, Band pass 67-1500 Hz), zeroed (DC remove, 0.3smoving window), rectified, and smoothed (RMS amplitude, 10ms moving window) using Spike2 software (version 8.03b). For normalization, data points in each of the conditions were divided by the average peak value of the 20 baseline swimming bursts. For each muscle, burst and interburst duration were measured from the rectified signal, and amplitude and integral were calculated from the smoothed signal. EMG datasets were normally distributed but group size varied. For the VGaT and VGluT2 cohorts, we used a paired 2 tailed t-test to assess for differences between saline and CNO conditions, but due to small SerT group sizes, we used a Wilcoxon matched pairs signed rank test for SerT group data.

To obtain composite EMG traces, in representative cases, for each of the 20 scored TA bursts during walking at baseline and CNO conditions, and in 3-8 TA bursts representing start and stop bursts at baseline, a 300ms segment starting 50ms before burst onset time was extracted. These 20 segments were aligned, averaged point-by-point into composite traces with +/- standard error of the mean, and baseline and CNO traces were overlaid on a shared axis (MATLAB).

*h) Optogenetic experiments:* Gait was analyzed as in the DREADD study for the paradigm with long trains of optogenetic stimulation. For the phase specificity paradigm, EMG traces were analyzed. TA bursts were categorized based upon the timing of the pulse to the burst onset (i.e. >0.1s before burst onset (n=18-29/case), <0.1s before burst onset (n=30-67/case), during burst (n=42-74/case), or after burst termination (n=55-62/case). For each of these 4 categories, we analyzed TA burst duration and maximum amplitude, as in the DREADD study, except that data in each mouse was normalized to quiet rest at baseline. Two way ANOVA was then used to assess the interaction of timing of pulse delivery and burst duration or peak amplitude. Of note, steps with two pulses falling within the burst were grouped with steps with one pulse falling within the burst. Steps where the nearest pulse fell before burst onset, but which also contained a pulse within the burst, were excluded from analysis. For visualization, in 3 cases representing distinct mRF sites, composite EMG traces were derived from smoothed EMG at baseline and from >0.1s before burst onset, <0.1 s before burst onset, during burst, or after burst offset, which had been normalized to average peak baseline amplitude, represented as average +/- standard error of the mean.

### Histological analysis

We digitized whole slides (Hamamatsu Nanozoomer XR RRID:SCR_017105) or details of sections (Zeiss Axioskop for brightfield or epifluorescent illumination; Zeiss LSM5 for confocal imaging). Brightness and contrast were adjusted using Adobe Photoshop CS6 or CC2018, NIH-ImageJ64 (RRID:SCR_003073) or FIJI software (RRID:SCR_002285)

#### Reconstructions of injection sites

We reconstructed each injection site from sections at intervals of 320µm as outlined in Supplementary Figure 2. We assigned an opacity of 8% to the core of each injection and 2% to the periphery. Individual cases were then merged as separate layers into a composite series as described previously (25). Cases were then grouped based upon gait Z-scores, and composite grayscale group images were pseudo-colored using complementary colors for visual subtraction or in JET to visualize hotspots using NIH-Image software.

#### Mapping of RSNs

To compose a map of VGaT and VGluT2 RSN neurons, we identified CTb^+^ RSNs immunohistochemically in combination with *in situ* hybridization to visualize VGaT or VGluT2 mRNA. In two representative cases, one with cervical cord projecting CTb^+^ RSNs and one with lumbar cord projecting CTb^+^ RSNs we reconstructed brainstem levels at intervals of 320µm as described before (25).

#### Spinal projections

Global distribution: We selected spinal sections of representative cases from VGaT, VGluT2 and SerT cohorts with small AAV transfection sites and low versus high Z-scores to determine whether projection patterns to the upper and mid lumbar cord varied in distribution. In each case, 3 upper- and 3 mid-lumbar sections were digitized at a resolution of 7680×4320 pixels. Using ImageJ software, color thresholding was applied to select labeled axons and terminals in each image and convert them into binary images. The 3 images/level were then superimposed into a composite image and a 5-pixel radius mean filter was applied, followed by a color lookup table to code a color gradient onto the pixel values for visualization.

#### Spinal projections

Density measurements: Given the marked variation in projection patterns among VGaT cases with smaller injection sites and distinct Z-scores, we then quantified hM3Dq-mCherry labeled boutons in the lumbar cord of 10 VGaT-ires-cre mice from 2 subsequent cohorts of 5 animals with AAV-DIO-hM3Dq-mCherry injections into the mRF to correlate regional bouton density with gait and EMG data. Image stacks were acquired in 6 regions of interest: lamina I-II, lamina III-IV, lateral lamina V, lamina VIII, lamina IX and area X, from 3 lumbar cord sections/mouse with a Zeiss LSM 5 Pascal confocal microscope using Zeiss ZEN 2009, 6.0 SP2 software under Plan-Achromat 63x/1.4 oil DIC objective, and excitation source 543nm rhodamine laser with emission filter set at 560nm and intensity of 70nM. Ten slices of 0.8µm were captured at intervals of 0.8µm, scan speed of 6, and pinhole setting of 96 (diameter: 1 airy-unit). Image size was 142.7µm X 142.7µm with an 8 bit depth and pixel size of 0.14µm. Z-stacks, each representing 162,906 µm^3^, were compressed using ImageJ (standard deviation compression), from which we quantified bouton-like profiles in each of the ROIs. The relation between high versus low Z^SwT^ and ROI was determined using 2 way ANOVA followed by Sidak’s multiple comparison test. We calculated Pearson correlation coefficient between counts in each of the ROIs and Z^SwT^ scores.

#### Spinal projections

Identification of neuronal sub-specialization: To distinguish target substrates from neighboring mRF^VGaT^ and mRF^VGluT2^ sites, following small injections of AAV-DIO-synaptophysin-mCherry into the mRF of VGaT or VGluT2-ires-cre mice (n=8), we measured size and number of synaptophysin-mCherry fusion protein labeled bouton-like profiles in the spinal cord using confocal imaging (as above), using 0.8µm thick slices at intervals of >4 slices to avoid overlap of bouton profiles. We compared appositions upon CTb labeled motoneuron somata and proximal dendrites in lamina IX, of ChAT-IR motoneurons, dendrites, and C-boutons in lamina IX, and of ChAT-IR premotor interneurons in area X using Ordinary One way ANOVA followed by Sidak’s multiple comparison test or Kruskal-Wallis test followed by Dunn’s multiple comparison test.

## Data availability statement

The data that support the findings of this study are presented in the Figures, Supplementary Figures, and an extensive set of Tables. Additional source data can be made available through the corresponding author upon reasonable request.

## Code availability statement

Custom MATLAB code was used to extract gait data from video files and Spike2 code was used to deliver optogenetic pulses based upon EMG feedback. Code has been made available: https://sourceforge.net/projects/gait-analysis-gui/ https://sourceforge.net/projects/gait-emg-step-trigger/

## Acknowledgements

The authors would like to thank Dr Brad Lowell (BIDMC) for AAV-DIO-synaptophysin-mCherry and Dr Susan Dymecki (Harvard Medical School) for SerT-cre breeders. This work was supported by R01 NS079623 (VV), R25 NS070682 (AK), the JG Foundation (VV) and donations from grateful patients (VV).

## Adherence to scientific standards

Reporting of study details is in accordance with ARRIVE guidelines for animal research.

## Competing interests

The authors declare no competing interests.

## Author Contributions

AW: electrophysiological recordings, analyses, code, preparation of figures and tables, data organization, editing

AK: optogenetic study, compilation and organization of code, editing

SL: EMG and kinematic analyses, preparation of figures and tables; editing

TS: behavioral testing, histology, gait and behavioral scoring; editing

BE: gait analysis code, gait scoring and analyses, editing

LB: confocal quantification, preparation of figures and tables, editing

AL: study design, sensory experiments, editing

VGV: conception, study design, surgeries and instrumentation, behavioral testing, recordings, analyses, confocal imaging, drafting of manuscript, preparation of figures and tables

---

**Video 1: Slowed, *slow motion*-like walking induced by mRF^VGaT-hM3Dq^ activation**

Representative example of *Slomo* walking style on a runway and movement along a beam following by mRF^VGaT-hM3Dq^ activation. The same mouse is shown following saline i.p. (top walkway, left beam) and 1 hour following 0.3mg/kg CNO i.p. (bottom walkway, right beam). Videos from the walkway and beam are played at 25% and 100% of real-time speed, respectively. The transfection site was bilateral and centered in the ventral mRF in the caudal pons.

**Video 2: Slowed, *shuffling-like* walking induced by mRF^VGluT2-hM3Dq^ activation**

Representative example of *Shuffle* walking style on a runway and movement along a beam following mRF^VGluT2-hM3Dq^ activation. The same mouse is shown following saline i.p. (top walkway, left beam) and 1 hour following 0.3mg/kg CNO i.p. (bottom walkway, right beam). Videos from the walkway and beam are played at 25% and 100% of real-time speed, respectively. The transfection site was centered in the ventral mRF immediately dorsal to the rostral pole of the inferior olive.

**Video 3: Accelerated walking induced by mRF^SerT-hM3Dq^ activation**

Representative example of a *Faster* gait following mRF^SerT-hM3Dq^ activation. The same mouse is shown following saline i.p. (top walkway, left beam) and 1 hour following 0.3mg/kg CNO i.p. (bottom walkway, right beam). Videos from the walkway and beam are played at 25% and 100% of real-time speed, respectively. The transfection site was centered in the raphe obscurus.

**Video 4: Slowed, *slow motion-like* activity induced by optogenetic mRF^VGaT-ChR2^ activation**

Representative example of the effect of mRF^VGaT-ChR2^ activation from the *Slomo* hotspot on gait. The same mouse is shown at baseline (top walkway) and during optogenetic stimulation (bottom walkway). Videos from the walkway and beam are played at 25% and 100% of real-time speed, respectively. The transfection site was bilateral and centered in the ventral mRF at and rostral to the rostral pole of the facial nucleus, with a midline fiber positioned immediately dorsal to the transfection site.

**Figure 2-Figure supplement 1: Dose and time response data.** Supplementary to results (paragraph 1), and methods. **a:** Effects of doses of 0.15, 0.3 and 0.6mg/kg CNO (i.p.) on average stride velocity and average swing time (n=14 VGaT-ires-cre mice with Z^SwT^>2; n=20 VGluT2-ires-cre mice with Z^SwT^<-2; one way ANOVA, followed by Dunnett’s multiple comparisons test with single pooled variance). **b:** Effect of a single dose of 0.3mg/kg CNO on swing time and stride velocity at different time points (n=5 VGaT-ires-cre mice with Z^SwT^>2; Friedman test, followed by Dunn’s multiple comparisons test). Table S1 for statistical details. Bars indicate SD.

**Figure 3-Figure supplement 1: Gait signatures and postural characteristics of slowed gait in the non-slomo (Z^Swt^<2) VGaT-hM3Dq group.** Supplementary to Fig. 3; see Table S3 for statistical details. Effects of CNO activation on gait signatures of the VGaT-hM3Dq group with Z^SwT^<2. Scatter plots depict swing time, stance time, stride length and frequency as a function of stride velocity or stride length as a function of swing velocity in saline (black) or CNO (color) conditions. A sum of squares F-test was used to assess whether saline and CNO datasets share regression lines (α 0.001) in the walking speed range (3-16cm/s). Polar plots summarize temporal coupling of the hindlimbs (Watson and Williams test; α 0.05). Stick figures represent left 5^th^ metatarsal, ankle, knee, trochanter, iliac crest and tail base during the swing phase in the CNO condition in representative mice. Bar graphs show averaged metrics in saline (gray) and CNO (color) conditions (two tailed paired t-test; α 0.05; bars represent SD) in each mouse.

**Figure 3-Figure supplement 2: Quantification of changes in anterior tibial EMG in experimental hM3Dq transfected VGaT mice.** Supplementary to Fig. 3. **a:** Graphs showing anterior tibial (TA) burst and interburst duration, amplitude and area under the curve (AUC) during steady walking in saline and CNO conditions for each of the Z score based subgroups of VGaT hM3Dq mice. Bars represent average, error bars SD. Two tailed paired t-test between conditions. EMG amplitude and AUC were normalized to the amplitude and AUC during swimming, with recruitment being near maximum, in the saline condition. Data is not controlled for speed. Table S4 for statistical details including interburst, swimming, gastrocnemius and WT control data. **b:** Rectified averaged EMG during ground walking and swimming of a representative VGaT^hM3Dq^ slomo mouse (Z^SwT^>3) showing activity of biceps femoris (BF) and triceps muscles (Tr; a forelimb muscle), in addition to TA activity (as in Fig. 3) during saline and CNO conditions. **c:** Correlation between Z^SwT^ and TA Z^BurstDuration^ in VGaT^hM3Dq^ mice instrumented with EMG (Pearson correlation coefficient; Table S5). Z scores indicate the change between baseline and CNO activation conditions. Changes in TA burst duration are strongly correlated with changes in swing time in VGaT mice. **d:** Correlation between Z^SwT^ and Z^BurstAmplitude^ in VGaT^hM3Dq^ mice (Pearson correlation coefficient; Table S5). Changes in TA burst amplitude are not correlated with changes in swing time.

**Figure 3-Figure supplement 3: Absence of gait asymmetries following activation of unilaterally transfected VGaT^hM3Dq^ neurons in the slow motion hotspot.** Supplementary to Fig. 3; text results. Assessment of spatial (stride length) and temporal (swing time) asymmetries following unilateral transfection of VGaT-hM3Dq neurons involving the slow-motion hotspot (N=6). i: Stride length or swing time ipsi- or contralateral to the transfected site did not vary between saline and CNO conditions. ii: In line with this, in each of the saline or CNO conditions, stride length and swing time of the ipsi- and contralateral hindlimbs were similar. See Table S7 for statistical details. See Supplementary Fig. 5 for injection sites.

**Figure 4-Figure supplement 1: Gait signatures and postural characteristics of slowed gait in the non-shuffle (Z^Swt^>-2) VGluT2-hM3Dq group.** Supplementary to Fig. 4; see Table S8 for statistical details. Effects of CNO activation on gait signatures of the VGluT2-hM3Dq group with Z^SwT^>-2. Scatter plots depict swing time, stance time, stride length and frequency as a function of stride velocity or stride length as a function of swing velocity in saline (black) or CNO (color) conditions. A sum of squares F-test was used to assess whether saline and CNO datasets share regression lines (α 0.001) in the walking speed range (3-16cm/s). Polar plots summarize temporal coupling of the hindlimbs (Watson and Williams test; α 0.05). Stick figures represent left 5^th^ metatarsal, ankle, knee, trochanter, iliac crest and tail base during the swing phase in the CNO condition in representative mice. Bar graphs show averaged metrics in saline (gray) and CNO (color) conditions (two tailed paired t-test; α 0.05; bars represent SD) in each mouse.

**Figure 4-Figure supplement 2: Quantification of changes in anterior tibial EMG in experimental hM3Dq transfected VGluT2 mice.** Supplementary to Fig. 4. a: Graphs showing anterior tibial (TA) burst and interburst duration, amplitude and area under the curve (AUC) during steady walking in saline and CNO conditions for each of the Z score based subgroups of VGluT2^hM3Dq^ mice. Bars represent average, error bars SD. Two tailed paired t-test between conditions. EMG amplitude and AUC were normalized to the amplitude and AUC during swimming, with recruitment being near maximum, in the saline condition. Data is not controlled for speed. Table S10 for statistical details including interburst, swimming, gastrocnemius and WT control data. b: Correlation between Z^SwT^ and TA Z^BurstDuration^ in VGluT2^hM3Dq^ mice instrumented with EMG (Pearson correlation coefficient; Table S5). Z scores indicate the change between baseline and CNO activation conditions. Changes in TA burst duration are not strongly correlated with changes in swing time in VGluT2 mice, suggesting that factors other than timing of TA activity drive the shortened swing time, such as co-activation of antagonist muscles (see text). c: Correlation between Z^SwT^ and Z^BurstAmplitude^ in VGluT2^hM3Dq^ mice (Pearson correlation coefficient; Table S5). Changes in TA burst amplitude are not correlated with changes in swing time.

**Figure 4-Figure supplement 3: Does unilateral transfection of the sites for shuffle gait cause asymmetries in temporal and spatial gait metrics?** Supplementary to Fig. 4; text results. Assessment of spatial (stride length) and temporal (swing time) asymmetries following unilateral transfection of VGluT2-hM3Dq neurons involving the shuffle-like hotspot (N=15). i: Stride length and swing time decreased in the CNO condition compared to saline both ipsi- and contralateral to the transfected side. ii: The decrease in stride length in CNO condition did not differ between ipsi- and contralateral hindlimbs, but the decrease in swing time was larger in the hindlimb contralateral to the transfection site, indicating a gait asymmetry that may depend on a crossed circuit mechanism. See Table S11 for statistical details. See Supplementary Fig. 5 for injection sites.

**Figure 5-Figure supplement 1: Gait signatures and postural characteristics of slowed gait in the slow (Z^Vel^<2) SerT-hM3Dq group.** Supplementary to Fig. 5; see Table S12 for statistical details. Effects of CNO activation on gait signatures of the VGaT-hM3Dq group with Z^SwT^<2. Scatter plots depict swing time, stance time, stride length and frequency as a function of stride velocity or stride length as a function of swing velocity in saline (black) or CNO (color) conditions. A sum of squares F-test was used to assess whether saline and CNO datasets share regression lines (α 0.001) in the walking speed range (3-16cm/s). Polar plots summarize temporal coupling of the hindlimbs (Watson and Williams test; α 0.05). Stick figures represent left 5^th^ metatarsal, ankle, knee, trochanter, iliac crest and tail base during the swing phase in the CNO condition in representative mice. Bar graphs show averaged metrics in saline (gray) and CNO (color) conditions (two tailed paired t-test; α 0.05; bars represent SD) in each mouse.

**Figure 5-Figure supplement 2: Quantification of changes in anterior tibial EMG in experimental SerT-hM3Dq transfected mice.** Supplementary to Fig. 5. Graphs showing anterior tibial (TA) burst and interburst duration, amplitude and area under the curve (AUC) during steady walking in saline and CNO conditions for each of the Z score based subgroups of SerT^hM3Dq^ mice. Bars represent average, error bars SD. Two tailed paired t-test between conditions. EMG amplitude and AUC were normalized to the amplitude and AUC during swimming, with recruitment being near maximum, in the saline condition. Data is not controlled for speed. Table S13 for statistical details including interburst, swimming, gastrocnemius and WT control data.

**Figure 6-Figure supplement 1: Reconstruction of transfection sites.** Supplementary to Fig. 6. **a:** Reconstruction of individual transfection sites. i: Scanned slides with sections from the spinomedullary junction to the caudal pons showing AAV8-hsyn-DIO-hM3Dq-mcherry transfection sites of representative VGaT-ires-cre (Z^SwT^ 3.7, in Figures 2a, d ii, e), VGlut2-ires-cre (Z^SwT^ −4.7, Figure 3a, d ii, e) and SerT-cre (Z^StrV^ 2.1, Figure 4a, d ii, e). Tissue was immunostained with dsRed to visualize mCherry transfected neurons (brown) and c-fos (black nuclei). ii: Enlargement of dashed boxes in i, showing transfected neurons and extensive dendritic branches (brown) in relation to major landmarks. iii: Magnifications of dashed boxes in ii, showing c-fos immunoreactivity (black) in hM3Dq-mCherry-transfected neurons (brown) following CNO activation. iv: Based upon the presence of transfected neuronal somata, but not dendrites, core and periphery of transfection sites were demarcated. v: Core and periphery of transfection sites were assigned 8% and 3% levels of opacity, respectively. This process was repeated at rostrocaudal intervals of 320µm. i: bar= 1 mm; ii: bar = 200 µm; iii: bar= 50 µm; iv-v: bar = 100 µm. **b:** Overview of the total transfected region in each of the cre-lines, derived from superimposed transfection sites from each mouse. These data sets were used for mapping of Z score based hot spots (Fig. 6 and Supplementary Fig. S9). Key landmarks to identify appropriate level of the ventral mRF are indicated to the right.

**Figure 6-Figure supplement 2: Compilation of transfection sites in VGaT-ires-cre, VGluT2-ires-cre and SerT-cre cohorts based upon functional metrics.** Supplementary to Fig. 6. a-c: Transfection sites are represented based upon gait characteristics, i.e. Z scores for swing time (VGaT and VGluT2 cohorts) and stride velocity (SerT group) or of EMG characteristics (VGaT mice: Z^InterburstAmplitude^<-1.3; VGluT2 mice: Z^InterburstAmplitude^>2 as proxies for low and high tone respectively). Groups are further subdivided based upon laterality when relevant. Dashed boxes represent the hotspots of the VGaT *slomo*, VGluT2 *shuffle* and SerT *fast* subgroups, which all localize to the mRF but with preference for rostrocaudally different levels as summarized in Fig. 6. Decreased EMG amplitude in the VGaT group also localized to the ventral mRF but with a site extending far caudal from the *slomo* hotspot. The hotspot of increased EMG amplitude in the VGluT2 group localized to a region just dorsal and rostral to the *shuffle* hotspot. These data suggest that *slow motion*-like, *shuffle-like* gait and muscle tone are modulated via brainstem substrates that are distinct from eachother, albeit partially intermixed.

**Figure 6-Figure supplement 3: In contrast to nearby mRF sites, slomo and shuffle hotspots do not heavily innervate mid- and forebrain.** Photomicrographs of hM3-mCherry transfection sites in the mRF and their projections to forebrain, midbrain, cervical and lumbar spinal cord in VGaT-ires-cre (a and b) and VGluT2-ires-cre (c and d) mice. Injection sites restricted to the hotspots of Z>3 *slomo* (a) and Z<-3 *shuffle* (c) mice innervate spinal regions as presented in Fig. 6, i.e. deep dorsal horn (a; arrows) and area X (c; arrows). These sites do not heavily innervate other parts of the CNS. In contrast, transfection sites that involve inhibitory neurons caudal to and dorsal to the slomo hotspot (b) heavily innervate fore- and midbrain (asterisks) as well as the spinal ventral horn (b, arrows), whereas transfected excitatory neurons dorsal and rostral to the shuffle hotspot innervate discrete sites in the midbrain tegmentum (arrow; extending to the red nucleus-not shown), thalamus (asterisk) and spinal medial ventral horn (arrows). Series were stained for mCherry with DAB (brown) and c-fos (black, nuclear stain to verify activation of mCherry transfected neurons). Bar represents 1000µm for supraspinal regions and 500µm for spinal levels.

**Figure 7-Figure supplement 1: Validation of spinal connectivity, cell type specificity, vesicular transporter of hotspots for slomo, shuffle and faster walking gaits. a:** In selected mice with high [Z^SwT^ or ^StrV^] scores, following completion of behavioral experiments, a CTb injection was placed a spinal cord subregion that was most densely innervated by the respective mRF hotspot (Fig. 6). In addition, transfection specificity was determined using ISH or IHC at the site of transfection, and vesicular transporter specificity was confirmed using IHC at the spinal level. **b:** i: Injection sites of CTb into the dorsal horn in VGaT-ires-cre, intermediate zone of VGluT2-ires-cre and ventral half of the lumbar cord in SerT-cre mice. ii: Confocal images of hM3Dq-mCherry transfection sites centered to hotspots (see a) that drive Z scores in VGaT (Z^SwT^ 3.7), VGluT2 (Z^SwT^ −4.7) or SerT mice (Z^StrV^ 2.1). Note labeling of RSNs (green; CTb), many colocalizing (white) to mCherry transfected (magenta) neurons. iii, iv: ISH for VGaT or VGluT2 or IHC for SerT confirmed cell type specificity of hM3Dq-mCherry transfected neurons. v,vi: hM3Dq-mCherry labeled (dsRed-IR) terminals derived from transfected neurons in iii, in combination with immunostaining for VGaT, VGluT2, or SerT in the spinal cord demonstrated neurotransmitter specificity (i.e. no co-expression) in terminal boutons. Bar=150µm in i, 50µm in ii and iii, 10µm in iv-vi.

**Figure 7-Figure supplement 2: Spinal projections from ventral VGaT and VGluT2 mRF sites centered just adjacent to hotspots differ in density from those that originate from the hotspots.** Supplementary to Fig. 7 which illustrates spinal connectivity from the hotspots. a: Conditional anterograde tracing from sites near the hotspots was combined with retrograde tracing to identify interneurons or motoneurons. b-e: Epifluorescence or confocal images of transfection sites (i) and spinal projections (ii-v) in 2 VGaT-ires-cre mice with transfection sites extending (a) mediolaterally (*) and (b) caudal to the hotspot (Fig. 7) and of VGluT2-ires-cre and VGluT2-L10-GFP reporter mice with transfection sites (d) rostral and (e) caudal to the hotspot (Fig. 7). i: Transfection sites (magenta). ii: Mid lumbar cord projection patterns from sites in i (magenta) and CTb labeled motoneurons (green). iii: SYP+ terminal profiles (magenta) near proximal dendrites (d) or somata (mn) of CTb labeled motoneurons (green) (b-d) or VGluT2-L10-GFP neurons in area X, some of which represent commissural interneurons retrogradely labeled (blue) from the contralateral spinal cord (e). iv: SYP+ boutons (magenta) near ChAT-IR (green) motoneurons (mn) in lamina IX (b-d) or (e) VGluT2-L10-GFP neurons in area X. v: SYP+ boutons (magenta) near ChAT-IR interneurons in area X (b-d) or lamina VII (e). *** in b-ii: superficial dorsal horn projections; * in e: CTb labeled commissural VGluT2 interneuron. arrowheads in ii: dense lamina IX projections; X: area X; V: lamina V; IX lamina IX; NTS: nucleus of the solitary tract; rIO: rostral inferior olive; cVII: caudal facial nucleus; XII: hypoglossal nucleus; LRN: lateral reticular nucleus; L5: 5th lumbar segment. Bar= 500µm in i, 200µm in ii, 10µm in iii-v. See Figure 7-Figure supplement 3 and Table S15 for quantification of profiles in different regions of the lumbar gray matter at the group level.

**Figure 7-Figure supplement 3: Quantification of synaptophysin bouton-like profiles derived from rostrocaudally distinct mRF VGaT and VGluT2 regions to lumbar laminae IX and X show gradients in innervation density.**

**a:** Large hM3Dq transfection sites gave rise to widespread projections. This contrasts with discrete transfection sites that resulted in regional differentiation in projection density (Fig. 7, Figure 7-Figure supplement 2). **b:** Experimental paradigm to validate and quantify regional differentiation from projections from different ventral pontomedullary levels to lamina IX and X. **c:** Numbers of Synaptophysin (SYP) labeled bouton-like profiles in apposition to ChAT immunostained motoneuron somata, dendrites or C-boutons in lamina IX of the lumbar cord, derived from VGluT2+ (cyan) or VGaT+ (magenta) neurons in the ventral mRF. Dots represent the number of profiles in a 0.8µm x 142.7µm x 142.7µm confocal slice, with 2 slices at >4µm intervals per mouse from 3 VGaT-ires-cre and 3 VGluT2-ires-cre mice. **d and e:** Number of SYP+ bouton-like profiles in lamina IX (d) or in apposition to CTb labeled gluteal, gastrocnemius or anterior tibial motoneuron somata or proximal dendrites in lamina IX (e) of the mid lumbar spinal cord, ipsilateral to the injection site. **f and g:** Number of SYP+ bouton-like profiles in area X or in close apposition to ChAT immunolabeled interneuron somata or proximal dendrites in area X (g). Data are from representative mice per genotype per level, with 4-9 (lamina IX) or 3 (area X) non-adjacent sections examined per mouse, including a compressed 8µm x 142.7µm x 142.7µm confocal Z-stack from each region per section, ipsilateral to the injection site. Error bars represent SD. Ordinary One Way ANOVA or Kruskall Wallis followed by multiple comparison test (see Supplementary Table 15S).

## Notes

### Competing Interest Statement

The authors have declared no competing interest.

