## Supplemental Figures for "Contrasting walking styles map to discrete neural substrates in the mouse brainstem"

### Figure 2-Figure Supplement 1

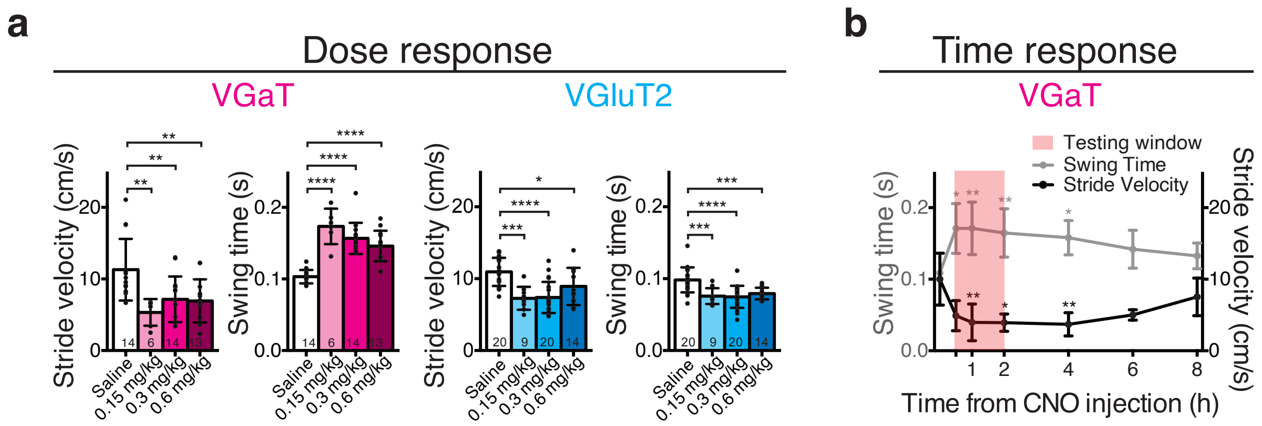


**Figure 2-Figure Supplement 1: Dose and time response data.** Supplementary to results (paragraph 1), and methods. **a:** Effects of doses of 0.15, 0.3 and 0.6mg/kg CNO (i.p.) on average stride velocity and average swing time (n=14 VGaT-ires-cre mice with $Z^{SwT}>2$; n=20 VGluT2-ires-cre mice with $Z^{SwT}<-2$; one way ANOVA, followed by Dunnett's multiple comparisons test with single pooled variance). **b:** Effect of a single dose of 0.3mg/kg CNO on swing time and stride velocity at different time points (n=5 VGaT-ires-cre mice with $Z^{SwT}>2$; Friedman test, followed by Dunn's multiple comparisons test). See Table S1 for statistical details. Bars indicate SD.

### Figure 3-Figure Supplement 1

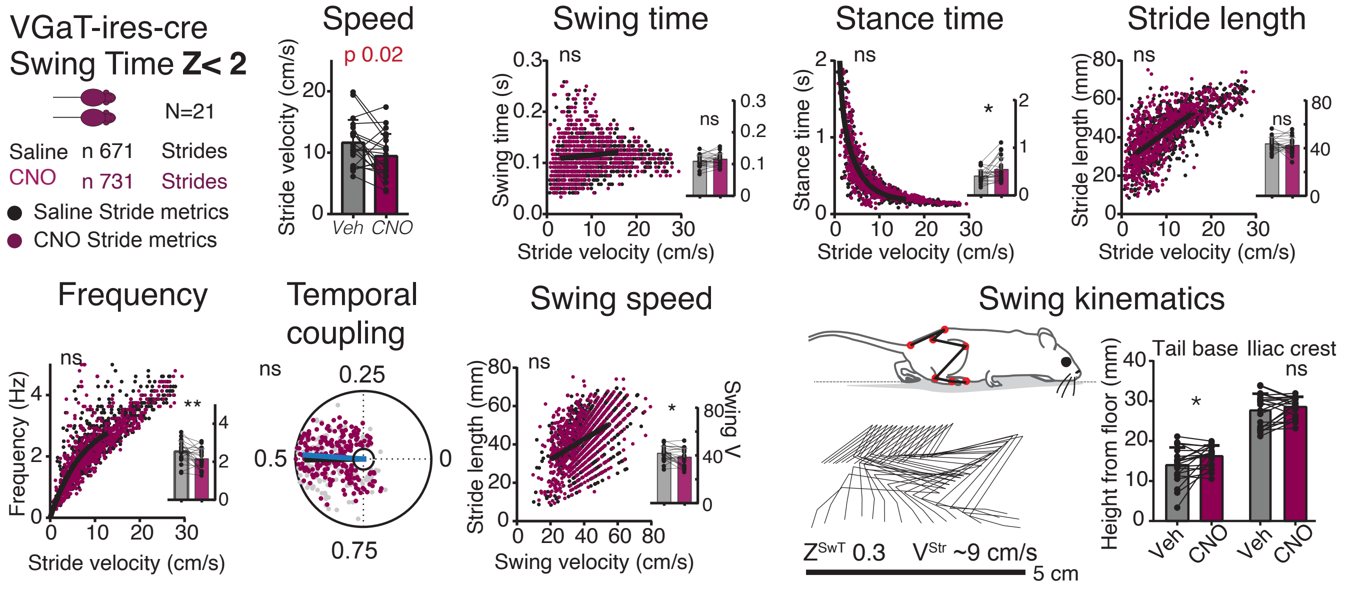


**Figure 3-Figure Supplement 1: Gait signatures and postural characteristics of slowed gait in the non-slomo (ZSwt<2) VGaT-hM3Dq group.** Accompanying Fig. 3; see Table S3 for statistical details. Effects of CNO activation on gait signatures of the VGaT-hM3Dq group with ZSwT<2. Scatter plots depict swing time, stance time, stride length and frequency as a function of stride velocity or stride length as a function of swing velocity in saline (black) or CNO (color) conditions. A sum of squares F-test was used to assess whether saline and CNO datasets share regression lines (  0.001) in the walking speed range (3-16cm/s). Polar plots summarize temporal coupling of the hindlimbs (Watson and Williams test;  0.05). Stick figures represent left 5th metatarsal, ankle, knee, trochanter, iliac crest and tail base during the swing phase in the CNO condition in representative mice. Bar graphs show averaged metrics in saline (gray) and CNO (color) conditions (two tailed paired t-test;  0.05; bars represent SD) in each mouse.

### Figure 3-Figure Supplement 2

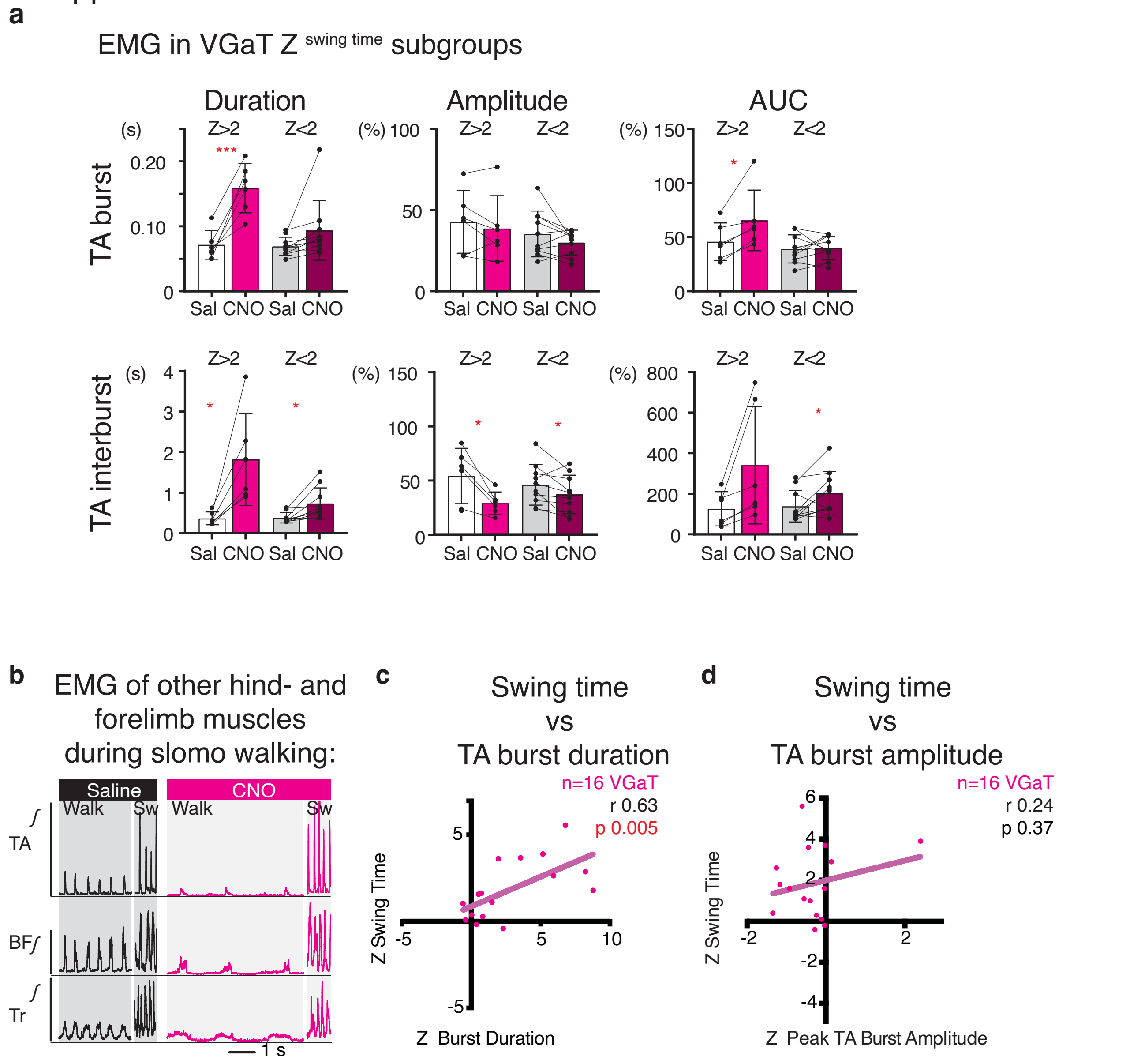


**Figure 3-Figure Supplement 2: Quantification of changes in anterior tibial EMG in experimental hM3Dq transfected VGaT mice.** **a:** Graphs showing anterior tibial (TA) burst and interburst duration, amplitude and area under the curve (AUC) during steady walking in saline and CNO conditions for each of the Z score based subgroups of $VGaT^{hM3Dq}$ mice. Bars represent average, error bars SD. Two tailed paired t-test between conditions. EMG amplitude and AUC were normalized to the amplitude and AUC during swimming, with recruitment being near maximum, in the saline condition. Data is not controlled for speed. Table S4 for statistical details including interburst, swimming, gastrocnemius and WT control data. **b:** Rectified averaged EMG during ground walking and swimming of a representative $VGaT^{hM3Dq}$ slomo mouse ($Z^{SwT}$>3) showing activity of biceps femoris (BF) and triceps muscles (Tr; a forelimb muscle), in addition to TA activity (as in Fig. 3) during saline and CNO conditions. **c:** Correlation between $Z^{SwT}$ and TA $Z^{BurstDuration}$ in $VGaT^{hM3Dq}$ mice instrumented with EMG (Pearson correlation coefficient; Table S5). Z scores indicate the change between baseline and CNO activation conditions. Changes in TA burst duration are strongly correlated with changes in swing time in VGaT mice. **d:** Correlation between $Z^{SwT}$ and $Z^{BurstAmplitude}$ in $VGaT^{hM3Dq}$ mice (Pearson correlation coefficient; Table S5). Changes in TA burst amplitude are not correlated with changes in swing time.

### Figure 3-Figure Supplement 3

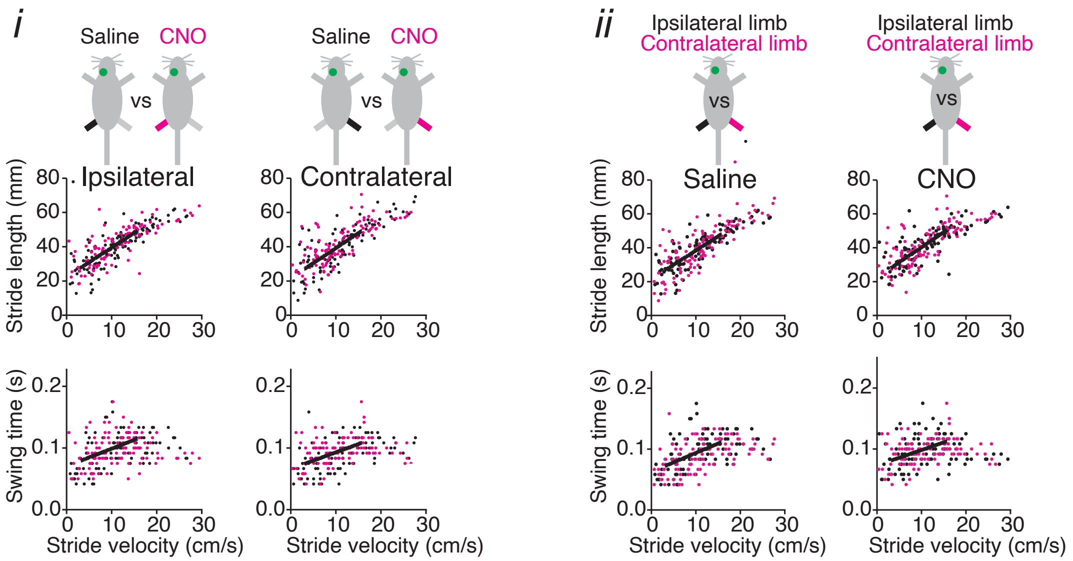


**Figure 3 -Figure Supplement 3: Absence of gait asymmetries following activation of unilaterally transfected VGaT$^{hM3Dq}$ neurons in the slow motion hotspot.** Supplementary to text results.

Assessment of spatial (stride length) and temporal (swingtime) asymmetries following unilateral transfection of VGaT$^{hM3Dq}$ neurons involving the slow-motion hotspot (N=6). **i:** Stride length or swing time ipsi- or contralateral to the transfected site did not vary between saline and CNO conditions. **ii:** In line with this, in each of the saline or CNO conditions, stride length and swing time of the ipsi- and contralateral hindlimbs were similar. Analyses as in Fig. 3. See Table S7 for statistical details. Fig. 6- Figure Supplement 2 for injection sites.

### Figure 4-Figure Supplement 1

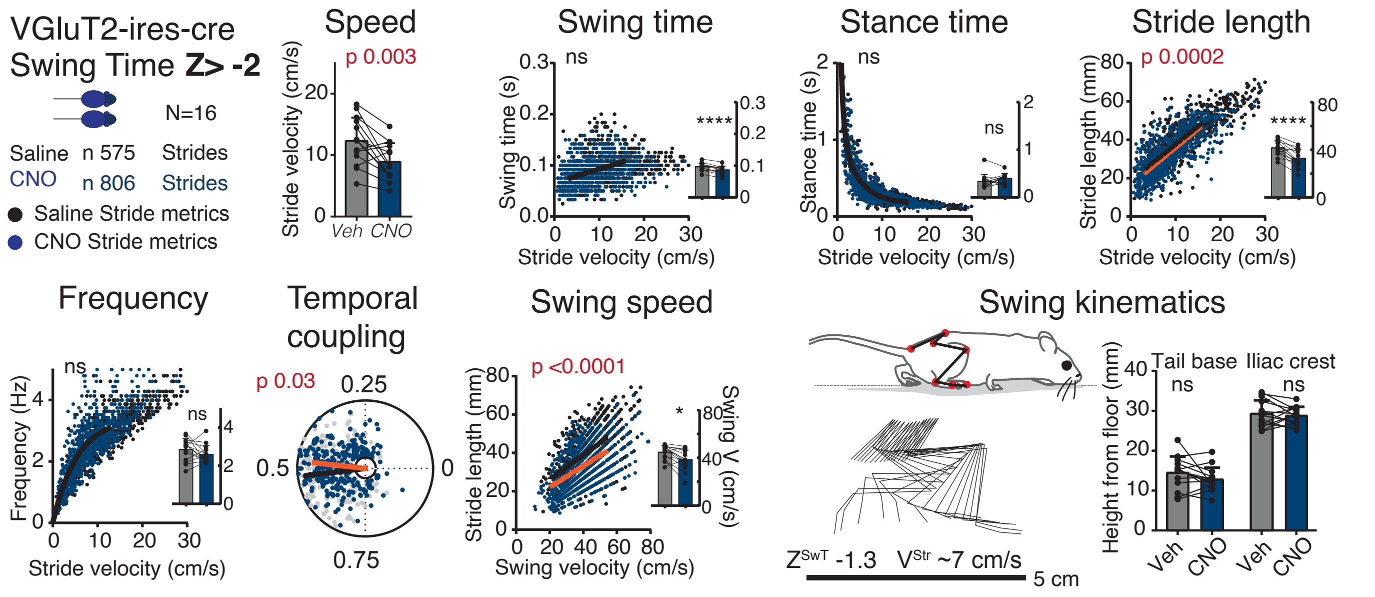


**Figure 4-Figure Supplement 1: Gait signatures and postural characteristics of slowed gait in the non-shuffle (ZSwt>-2) VGluT2-hM3Dq group.** See Table S8 for statistical details. Effects of CNO activation on gait signatures of the VGluT2-hM3Dq group with ZSwT>-2. Scatter plots depict swing time, stance time, stride length and frequency as a function of stride velocity or stride length as a function of swing velocity in saline (black) or CNO (color) conditions. A sum of squares F-test was used to assess whether saline and CNO datasets share regression lines (  0.001) in the walking speed range (3-16cm/s). Polar plots summarize temporal coupling of the hindlimbs (Watson and Williams test;  0.05). Stick figures represent left 5th metatarsal, ankle, knee, trochanter, iliac crest and tail base during the swing phase in the CNO condition in representative mice. Bar graphs show averaged metrics in saline (gray) and CNO (color) conditions (two tailed paired t-test;  0.05; bars represent SD) in each mouse.

### Figure 4-Figure Supplement 2

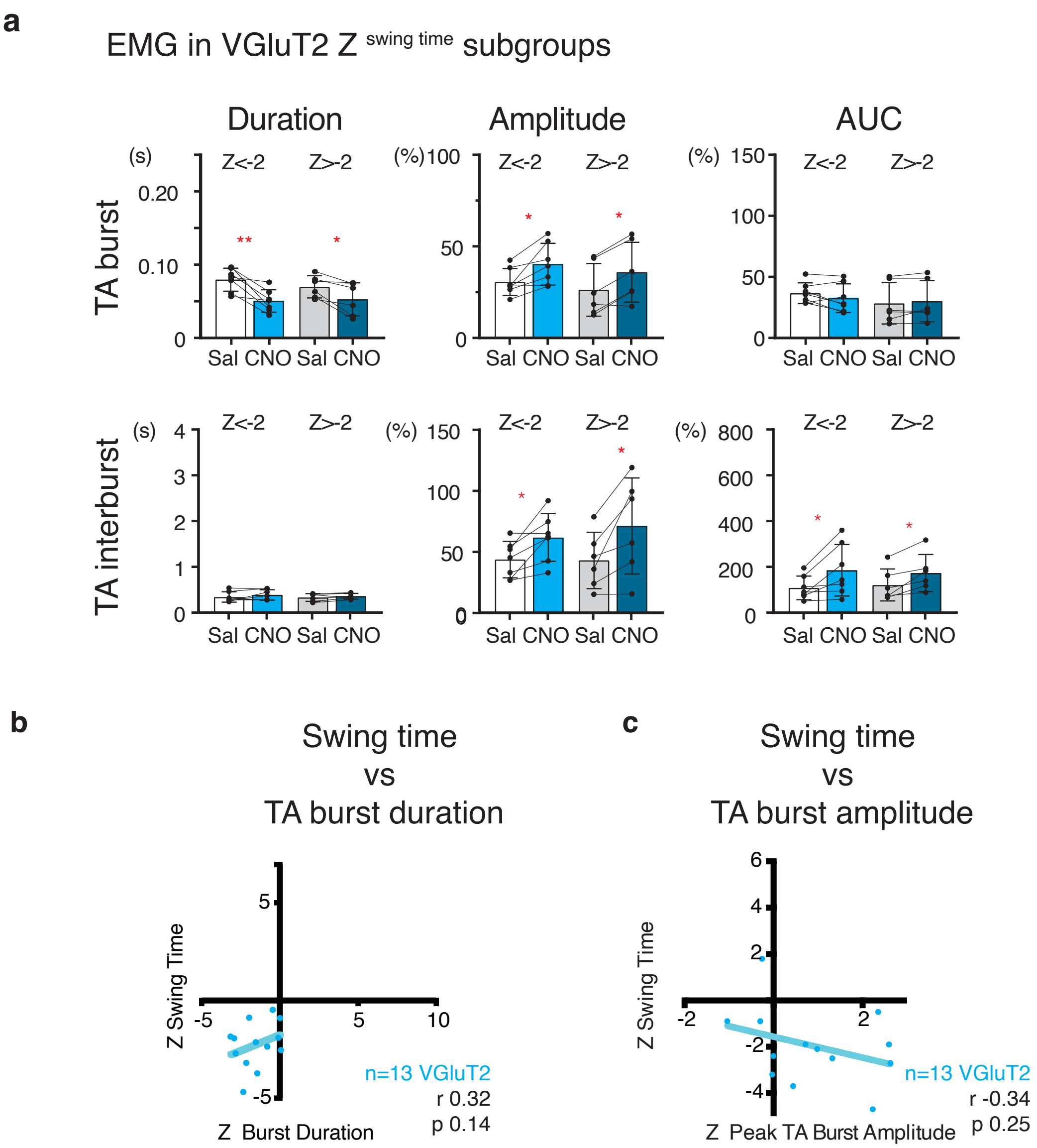


**Figure 4-Figure Supplement 2: Quantification of changes in anterior tibial EMG in experimental hM3Dq transfected VGluT2 mice.** **a:** Graphs showing anterior tibial (TA) burst and interburst duration, amplitude and area under the curve (AUC) during steady walking in saline and CNO conditions for each of the Z score based subgroups of VGluT2$^{hM3Dq}$ mice. Bars represent average, error bars SD. Two tailed paired t-test between conditions. EMG amplitude and AUC were normalized to the amplitude and AUC during swimming, with recruitment being near maximum, in the saline condition. Data was not corrected for speed. Tables S4 (VGaT), S10 (VGluT2) and S13 (SerT) for statistical details including interburst, swimming, gastrocnemius and WT control data. **b:** Correlation between $Z^{SwT}$ and TA $Z^{BurstDuration}$ in VGluT2$^{hM3Dq}$ mice instrumented with EMG (Pearson correlation coefficient; Table S5). Z scores indicate the change between baseline and CNO activation conditions. Changes in TA burst duration are not strongly correlated with changes in swing time in VGluT2 mice, suggesting that factors other than timing of TA activity drive the shortened swing time, such as co-activation of antagonist muscles (see text). **c:** Correlation between $Z^{SwT}$ and $Z^{BurstAmplitude}$ in VGluT2$^{hM3Dq}$ mice (Pearson correlation coefficient; Table S5). Changes in TA burst

### Figure 4- figure Supplement 3

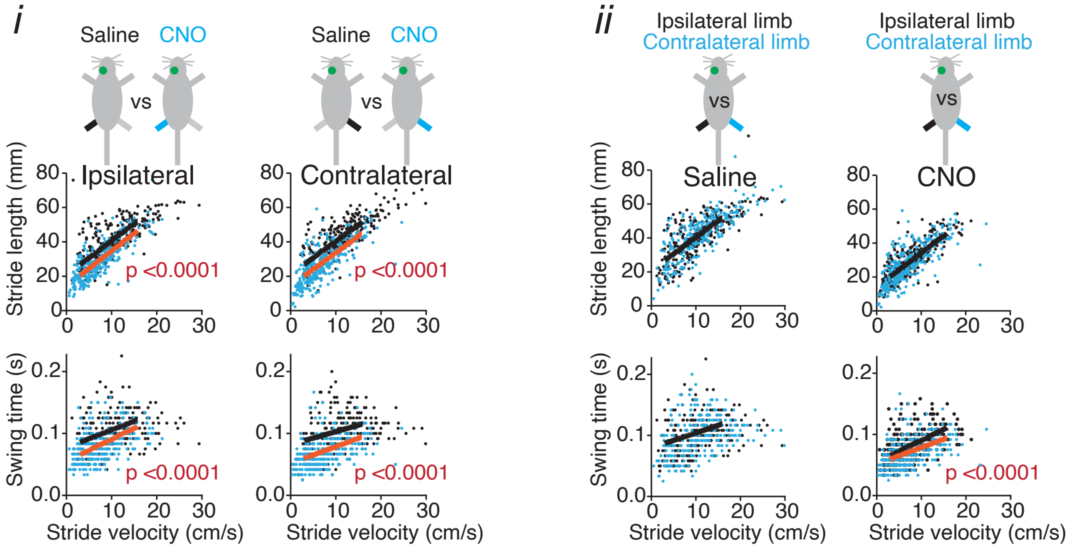


**Figure 4- Figure Supplement 3: Activation of unilaterally transfected VGluT2 neurons in the shuffle hotspot elicited asymmetries in temporal but not spatial gait metrics.** Supplementary to text results.
Assessment of spatial (stride length) and temporal (swingtime) asymmetries following unilateral transfection of $VGluT^{hM3Dq}$ neurons involving the shuffle-like hotspot (N=15). **i:** Stride length and swing time decreased in the CNO condition compared to saline both ipsi- and contralateral to the transfected side. **ii:** The decrease in stride length in CNO condition did not differ between ipsi- and contralateral hindlimbs, but the decrease in swing time was larger in the hindlimb *contra*lateral to the transfection site, indicating a gait assymmetry that may depend on a crossed circuit mechanism. Signature analyses as in Fig. 4. See Table S11 for statistical details. Fig. 6- Figure Supplement 2 for injection sites.

Figure 5-Figure Supplement 1

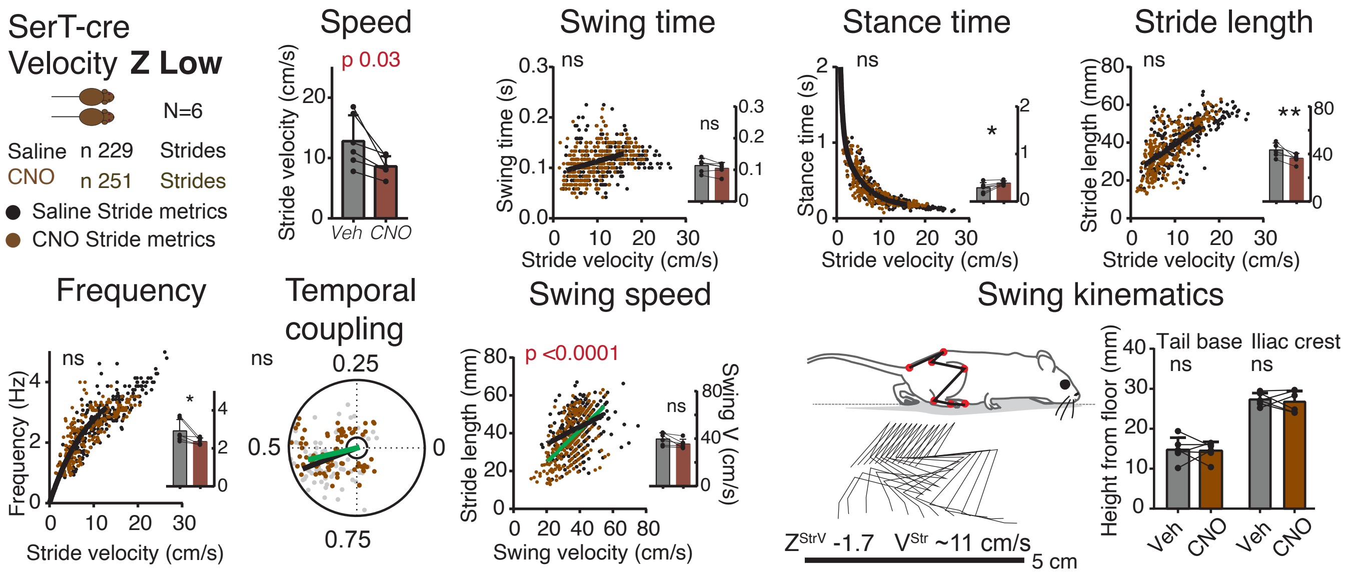


**Figure 5-Figure Supplement 1: Gait signatures and postural characteristics of slowed gait in the slow (ZVel low) SerT-hM3Dq group.** Accompanying Fig. 5; see Table S12 for statistical details. Effects of CNO activation on gait signatures of the SerT-hM3Dq group with low ZStrV. Scatter plots depict swing time, stance time, stride length and frequency as a function of stride velocity or stride length as a function of swing velocity in saline (black) or CNO (color) conditions. A sum of squares F-test was used to assess whether saline and CNO datasets share regression lines (  0.001) in the walking speed range (3-16cm/s). Polar plots summarize temporal coupling of the hindlimbs (Watson and Williams test;  0.05). Stick figures represent left 5th metatarsal, ankle, knee, trochanter, iliac crest and tail base during the swing phase in the CNO condition in representative mice. Bar graphs show averaged metrics in saline (gray) and CNO (color) conditions (two tailed paired t-test;  0.05; bars represent SD) in each mouse.

### Figure 5-Figure Supplement 2

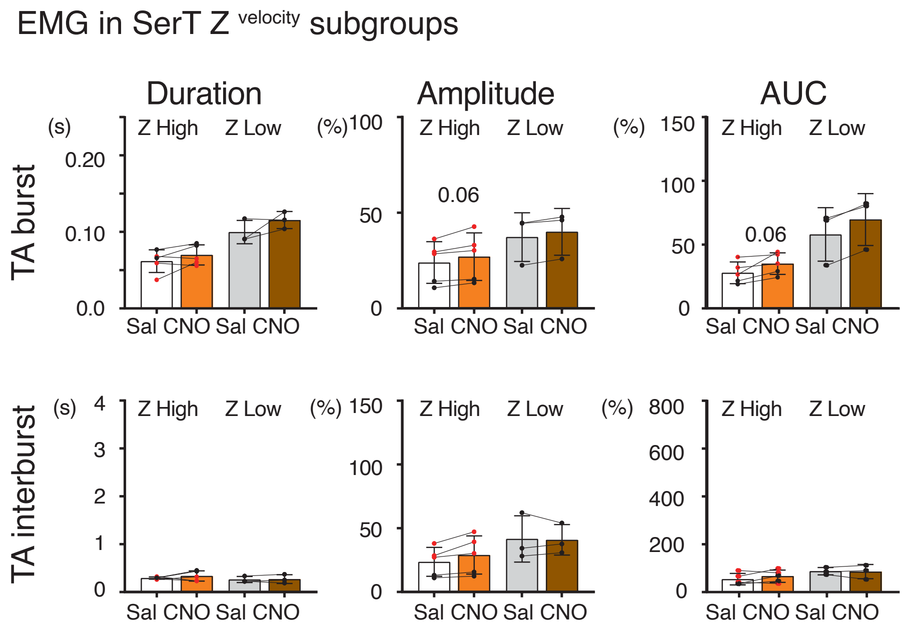


**Figure 5-Figure Supplement 2: Quantification of changes in anterior tibial EMG in experimental hM3Dq transfected SerT mice.** Graphs showing anterior tibial (TA) burst and interburst duration, amplitude and area under the curve (AUC) during steady walking in saline and CNO conditions for each of the Z score based subgroups of $SerT^{hM3Dq}$ mice. Bars represent average, error bars SD. Two tailed paired t-test between conditions. EMG amplitude and AUC were normalized to the amplitude and AUC during swimming, with recruitment being near maximum, in the saline condition. Data was not corrected for speed. Table S13 for statistical details including interburst, swimming, gastrocnemius and WT control data.

### Figure 6-Figure Supplement 1

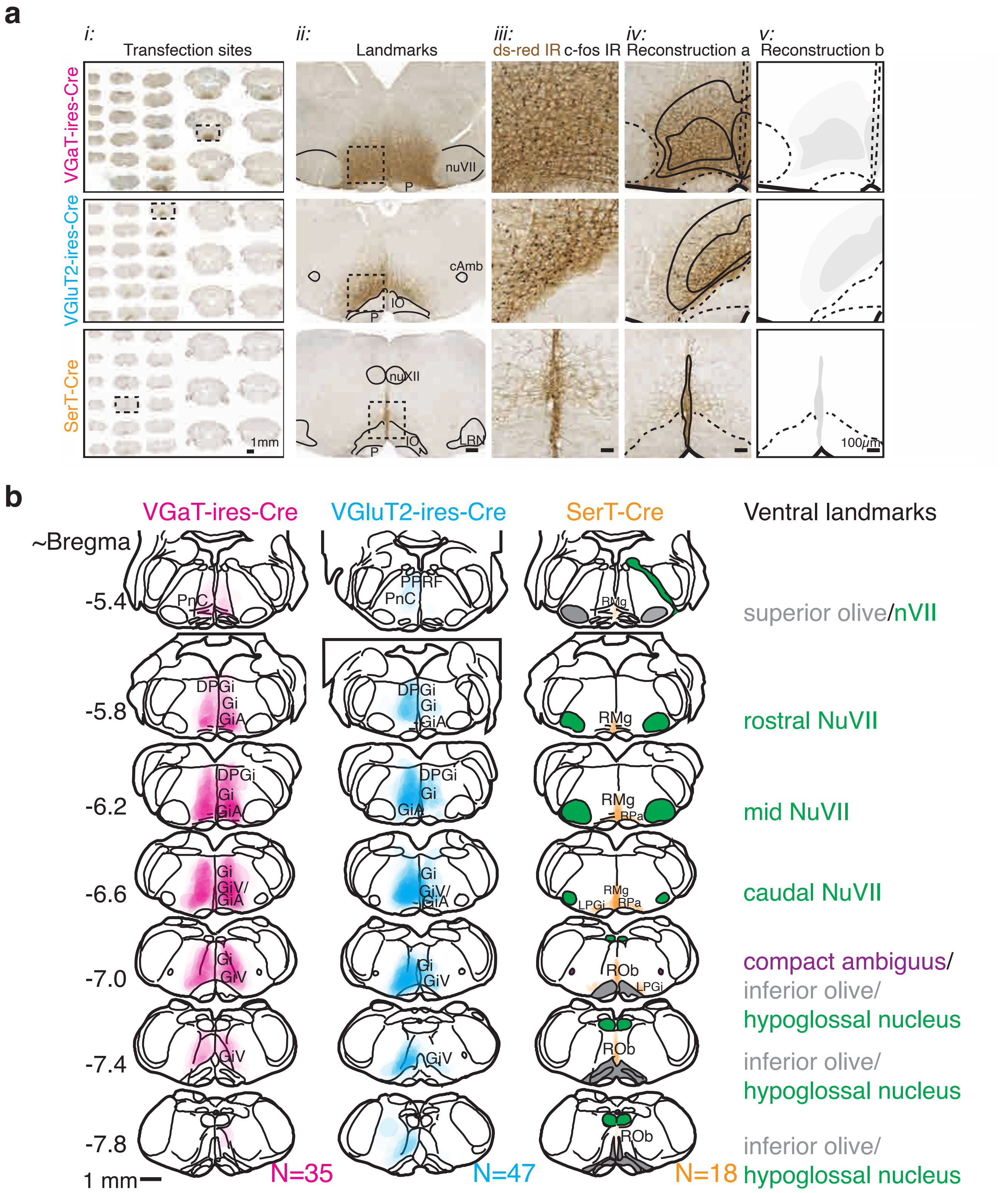


**Figure 6- Figure Supplement 1: Reconstruction of transfection sites.**

**a:** Reconstruction of individual transfection sites. i: Scanned slides with sections from the spinomedullary junction to the caudal pons showing AAV8-hsyn-DIO-hM3Dq-mcherry transfection sites of representative VGaT-ires-cre ($Z^{SwT}$ 3.7, in Fig. 3a, d ii, e), VGlut2-ires-cre ($Z^{SwT}$ -4.7, Fig. 4a, d ii, e) and SerT-cre ($Z^{StrV}$ 2.1, Fig. 5a, d ii, e). Tissue was immunostained with dsRed to visualize mCherry transfected neurons (brown) and c-fos (black nuclei). ii: Enlargement of dashed boxes in i, showing transfected neurons and extensive dendritic branches (brown) in relation to major landmarks. iii: Magnifications of dashed boxes in ii, showing c-fos immunoreactivity (black) in hM3Dq-mCherry-transfected neurons (brown) following CNO activation. iv: Based upon the presence of transfected neuronal somata, but not dendrites, core and periphery of transfection sites were demarcated. v: Core and periphery of transfection sites were assigned 8% and 3% levels of opacity, respectively. This process was repeated at rostrocaudal intervals of 320μm. i: bar= 1 mm; ii: bar = 200 μm; iii: bar= 50 μm; iv-v: bar = 100 μm. **b:** Overview of the total transfected region in each of the cre-lines, derived from superimposed transfection sites from each mouse. These data sets were used for mapping of Z score based hot spots (Fig. 6 and Figure 6-Figure Supplement 2). Key landmarks to identify appropriate level of the ventral mRF are indicated to the right.

Figure 6-Figure Supplement 2

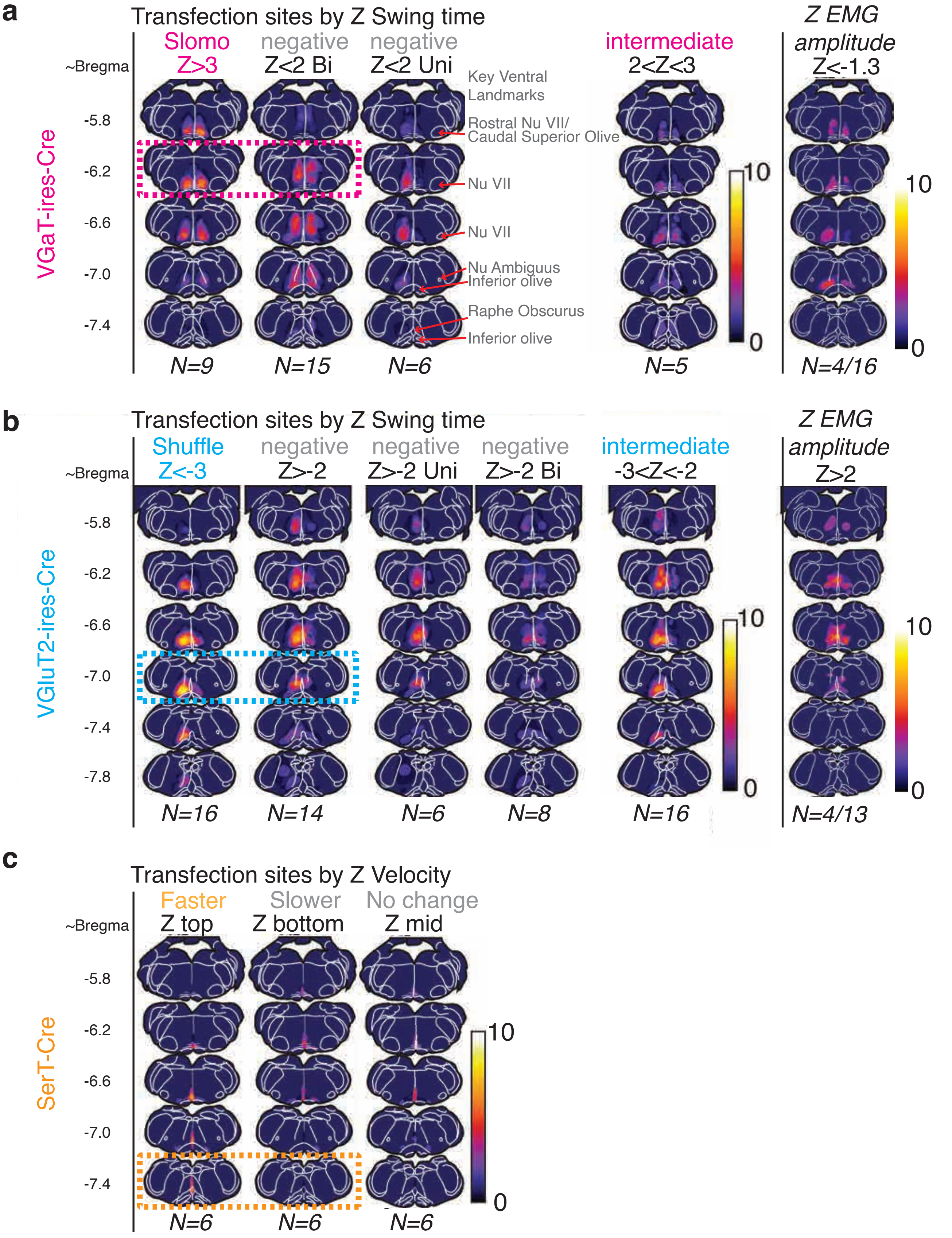


**Figure 6- Figure Supplement 1: Compilation of transfection sites in VGaT-ires-cre, VGluT2-ires-cre and SerT-cre cohorts based upon functional metrics. a-c:** Transfection sites are represented based upon gait characteristics, i.e. Z scores for swing time (VGaT and VGluT2 cohorts) and stride velocity (SerT group) or of EMG characteristics (VGaT mice: $Z^{InterburstAmplitude}$<-1.3; VGluT2 mice: $Z^{InterburstAmplitude}$>2 as proxies for low and high tone respectively). Groups are further subdivided based upon laterality when relevant. Dashed boxes represent the hotspots of the VGaT *slomo*, VGluT2 *shuffle* and SerT *fast* subgroups, which all localize the mRF but with preference for rostrocaudally different levels as summarized in Fig.6. Decreased in EMG amplitude in the VGaT group also localized to the ventral mRF but with a site extending far caudal from the *slomo* hotspot. The hotspot of increased EMG amplitude in the VGluT2 group localized to a region just dorsal and rostral to the *shuffle* hotspot. These data suggest that *slow motion*-like, *shuffle*-like gait and muscle tone are modulated via brainstem substrates that are distinct from eachother, albeit partially intermixed.

Figure 6-Figure Supplement 3

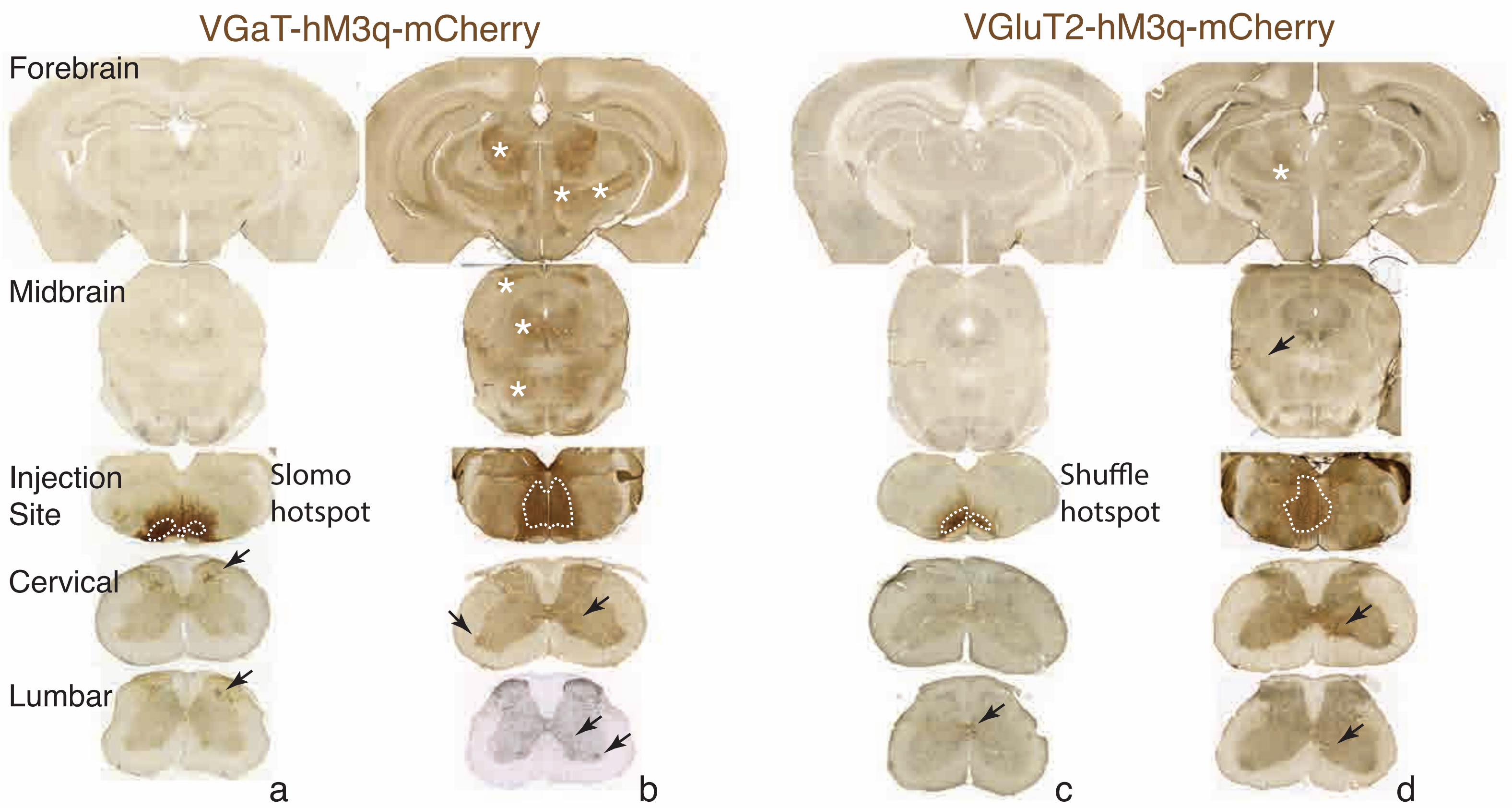


**Figure 6- Figure Supplement 3: Examples of the extent of projections from slomo- and shuffle hotspots and nearby regions in the mRF.** Photomicrographs of hM3-mCherry transfection sites in the mRF and their projections to forebrain, midbrain, cervical and lumbar spinal cord in VGaT-ires-cre (a and b) and VGluT2-ires-cre (c and d) mice. Injection sites restricted to the hotspots of Z>3 slomo (a) and Z<-3 shuffle (c) mice innervate spinal regions as presented in Fig. 6, i.e. deep dorsal horn (a; arrows) and area X (c; arrows). These sites do not heavily innervate other parts of the CNS. In contrast, transfection sites that involve inhibitory neurons caudal to and dorsal to the slomo hotspot (b) heavily innervate fore- and midbrain (asterisks) as well as the spinal ventral horn (b, arrows), whereas transfected excitatory neurons dorsal and rostral to the shuffle hotspot innervate discrete sites in the midbrain tegmentum (arrow; extending to the red nucleus-not shown), thalamus (asterisk) and spinal medial ventral horn (arrows). Series were stained for mCherry with DAB (brown) and c-fos (black, nuclear stain to verify activation of mCherry transfected neurons).

### Figure 7-Figure Supplement 1

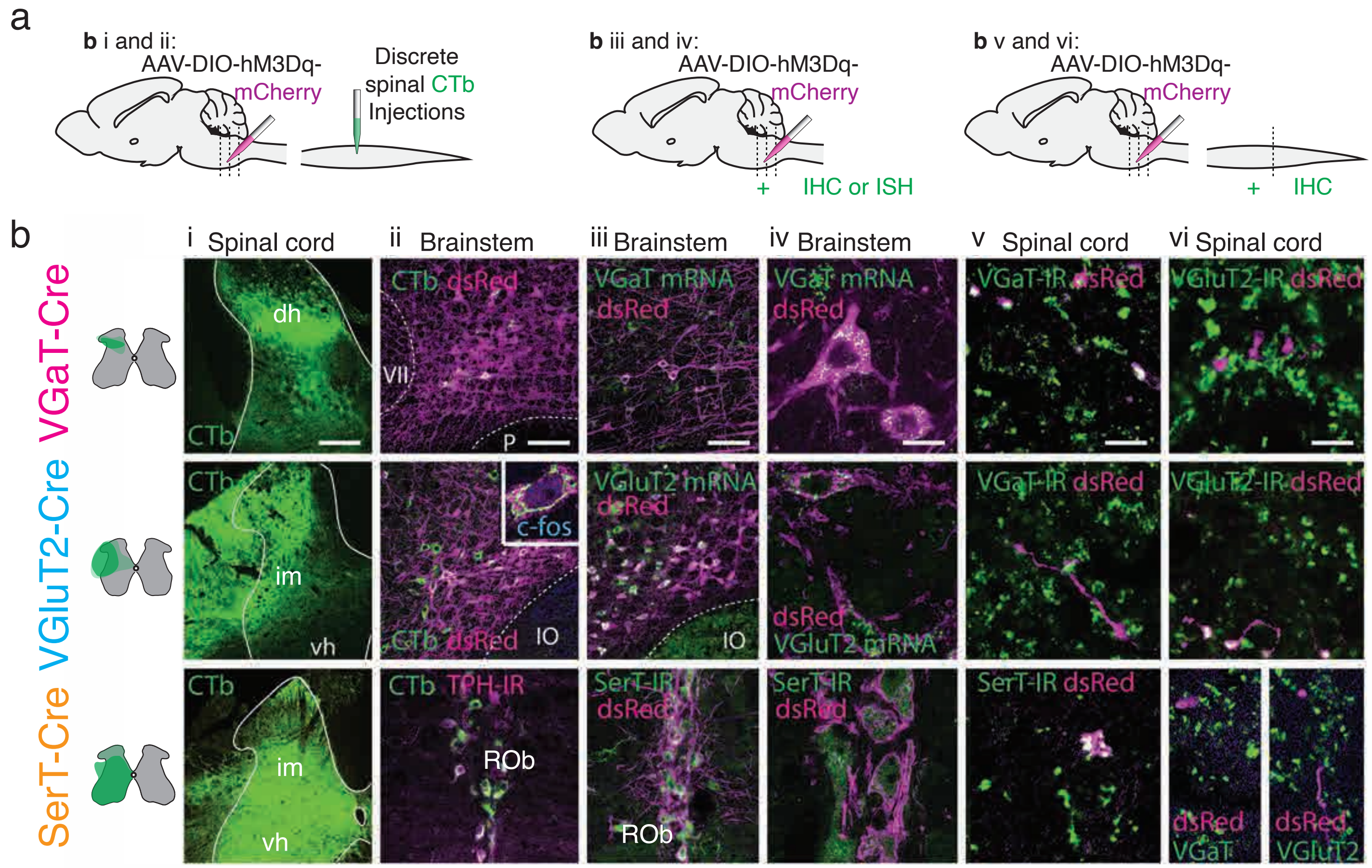


**Figure 7-Figure Supplement 1: Validation of spinal connectivity, cell type specificity, vesicular transporter of hotspots for slomo, shuffle and faster walking gaits. a:** In selected mice with high [$Z^{SwT}$ or $^{StrV}$] scores, following completion of behavioral experiments, a CTb injection was placed a spinal cord subregion that was most densely innervated by the respective mRF hotspot (Fig. 6). In addition, transfection specificity was determined using ISH or IHC at the site of transfection, and vesicular transporter specificity was confirmed using IHC at the spinal level. b: i: Injection sites of CTb into the dorsal horn in VGaT-ires-cre, intermediate zone of VGluT2-ires-cre and ventral half of the lumbar cord in SerT-cre mice. ii: Confocal images of hM3Dq-mCherry transfection sites centered to hotspots (see a) that drive Z scores in VGaT ($Z^{SwT}$ 3.7), VGluT2 ($Z^{SwT}$ -4.7) or SerT mice ($Z^{StrV}$ 2.1). Note labeling of RSNs (green; CTb), many colocalizing (white) to mCherry transfected (magenta) neurons. iii, iv: ISH for VGaT or VGluT2 or IHC for SerT confirmed cell type specificity of hM3Dq-mCherry transfected neurons. v,vi: hM3Dq-mCherry labeled (dsRed-IR) terminals derived from transfected neurons in iii, in combination with immunostaining for VGaT, VGluT2, or SerT in the spinal cord demonstrated neurotransmitter specificity (i.e. no co-expression) in terminal boutons. Bar=150$\mu$m in i, 50$\mu$m in ii and iii, 10$\mu$m in iv- vi.

Figure 7-Figure Supplement 2

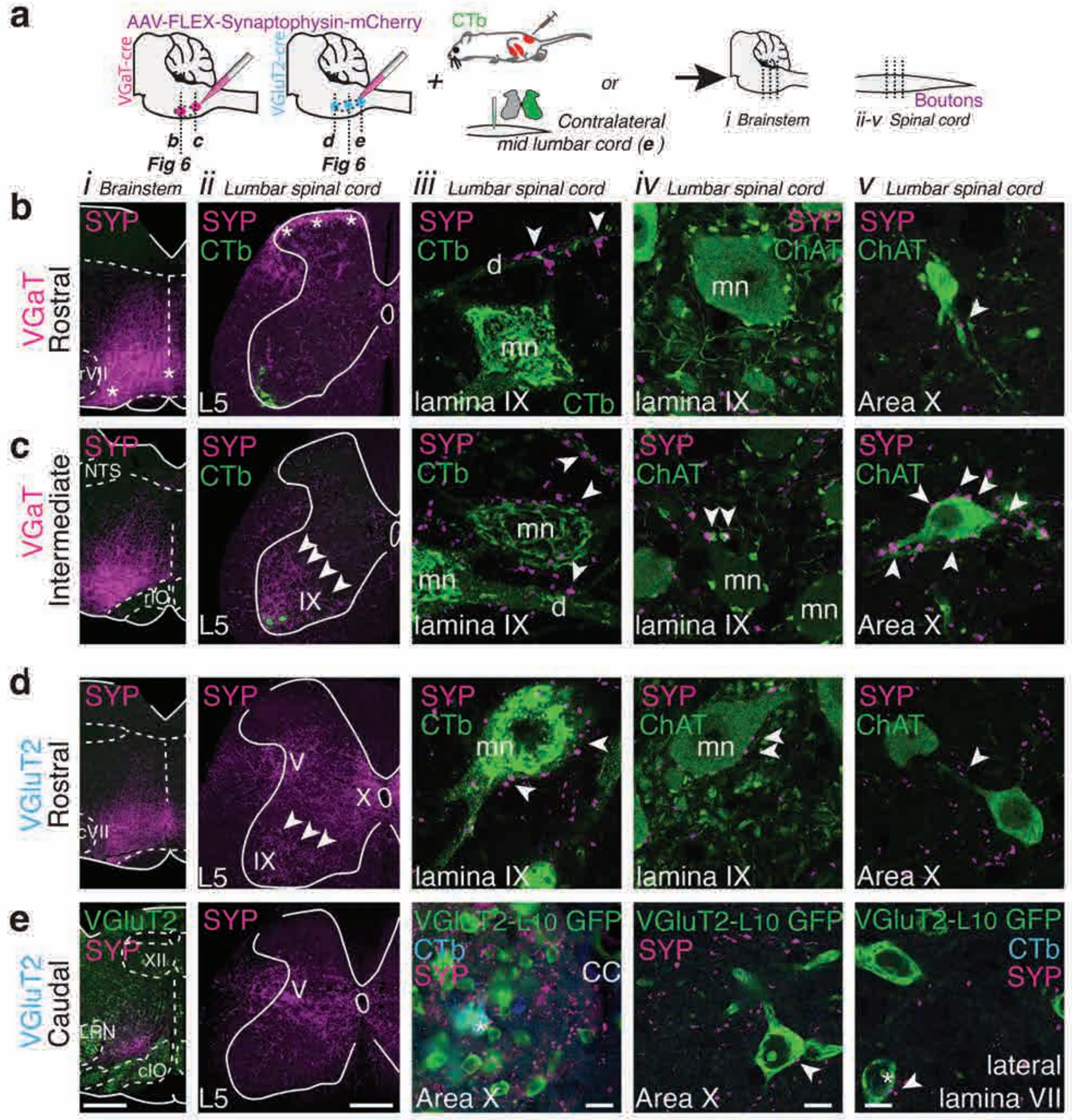


**Figure 7-Figure Supplement 2: Spinal projections from ventral VGaT and VGluT2 mRF sites centered just adjacent to hotspots differ in density from those that originate from the hotspots.** Supplementary to Fig. 7 which illustrates spinal connectivity from the hotspots.

**a:** Conditional anterograde tracing from sites near the hotspots was combined with retrograde tracing to identify interneurons or motoneurons. **b-e:** Epifluorescence or confocal images of transfection sites (i) and spinal projections (ii-v) in 2 VGaT-ires-cre mice with transfection sites extending (a) mediolaterally (*) and (b) caudal to the hotspot (Fig. 7) and of VGluT2-ires-cre and VGluT2-L10-GFP reporter mice with transfection sites (d) rostral and (e) caudal to the hotspot (Fig. 7). **i:** Transfection sites (magenta). **ii:** Mid lumbar cord projection patterns from sites in i (magenta) and CTb labeled motoneurons (green). **iii:** SYP+ terminal profiles (magenta) near proximal dendrites (d) or somata (mn) of CTb labeled motoneurons (green) (b-d) or VGluT2-L10-GFP neurons in area X, some of which represent commissural interneurons retrogradely labeled (blue) from the contralateral spinal cord (e). **iv:** SYP+ boutons (magenta) near ChAT-IR (green) motoneurons (mn) in lamina IX (b-d) or (e) VGluT2-L10-GFP neurons in area X. **v:** SYP+ boutons (magenta) near ChAT-IR interneurons in area X (b-d) or lamina VII (e). *** in b-ii: superficial dorsal horn projections; * in e: CTb labeled commissural VGluT2 interneuron. arrowheads in ii: dense lamina IX projections; X: area X; V: lamina V; IX lamina IX; NTS: nucleus of the solitary tract; rIO: rostral inferior olive; cVII: caudal facial nucleus; XII: hypoglossal nucleus; LRN: lateral reticular nucleus; L5: 5th lumbar segment.  Bar= 500μm in i, 200μm in ii, 10μm in iii-v. See Fig. 7-Figure Supplement 3 and Table S15 for quantification.

Figure 7 -Figure Supplement 3

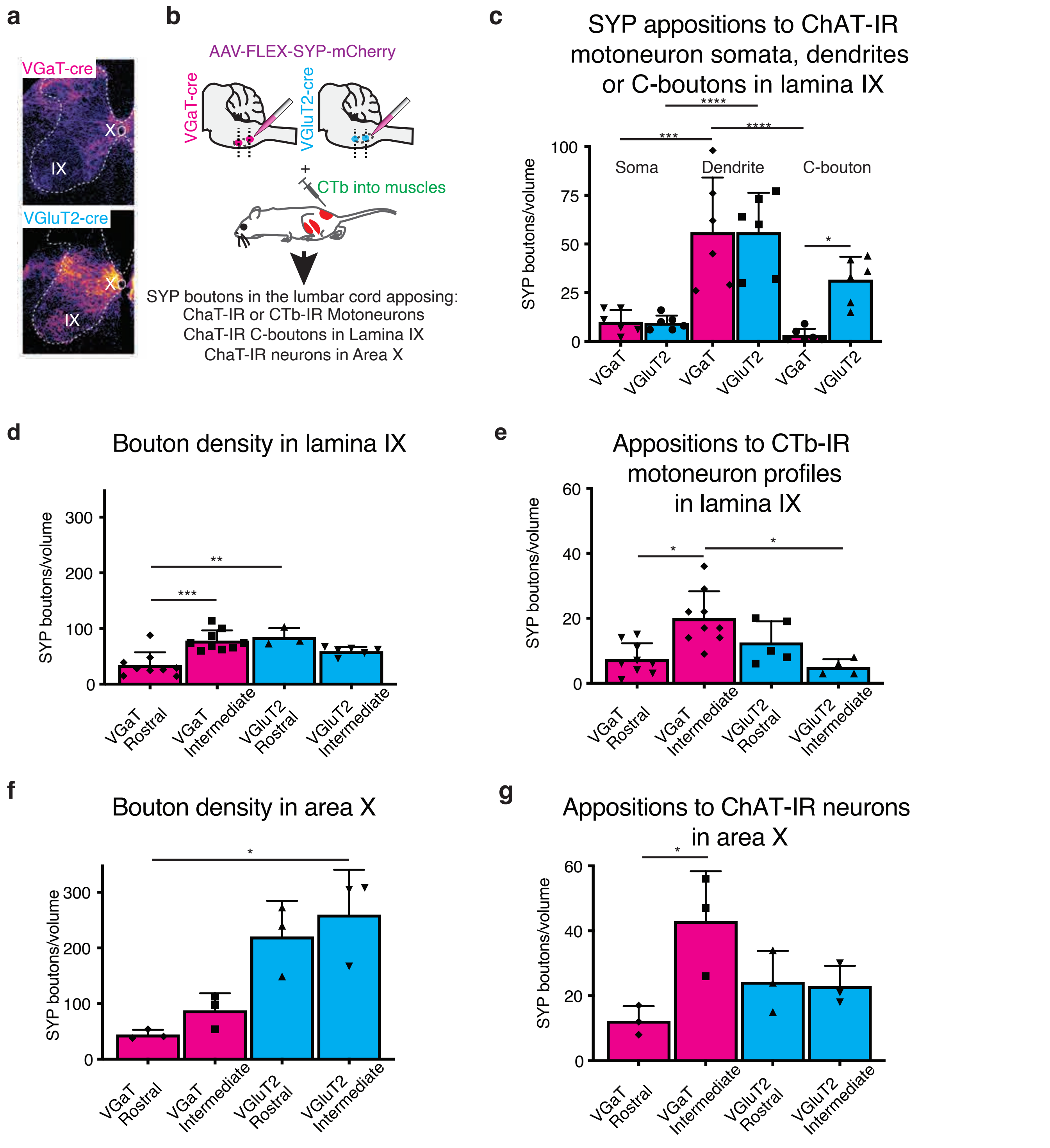


**Figure 7- Figure Supplement 3: Quantification of synaptophysin bouton-like profiles derived from rostrocaudally distinct mRF VGaT and VGluT2 regions to lumbar laminae IX and X show gradients in innervation density.**
**a:** Large hM3Dq transfection sites gave rise to widespread projections. This contrasts with discrete transfection sites that resulted in regional differentiation in projection density (Fig. 7, Fig. 7-Figure Supplement 2). **b:** Experimental paradigm to validate and quantify regional differentiation from projections from different ventral pontomedullary levels to lamina IX and X. **c:** Numbers of Synaptophysin (SYP) labeled bouton-like profiles in apposition to ChAT immunostained motoneuron somata, dendrites or C-boutons in lamina IX of the lumbar cord, derived from VGluT2+ (cyan) or VGaT+ (magenta) neurons in the ventral mRF. Dots represent the number of profiles in a 0.8$\mu$m x 142.7$\mu$m x 142.7$\mu$m confocal slice; 2 slices at >4$\mu$m intervals per mouse from 3 VGaT-ires-cre and 3 VGluT2-ires-cre mice. **d and e:** Number of SYP+ bouton-like profiles in lamina IX (d) or in apposition to CTb labeled gluteal, gastrocnemius or anterior tibial motoneuron somata or proximal dendrites in lamina IX (e) of the mid lumbar spinal cord, ipsilateral to the injection site. **f and g:** Number of SYP+ bouton-like profiles in area X or in close apposition to ChAT immunolabeled interneuron somata or proximal dendrites in area X (g). Data are from representative mice per genotype per level, with 4-9 (lamina IX) or 3 (area X) non-adjacent sections examined per mouse, including a compressed 8$\mu$m x 142.7$\mu$m x 142.7$\mu$m confocal Z-stack from each region per section, ipsilateral to the injection site. Error bars represent SD. Ordinary One Way ANOVA or Kruskall Wallis followed by multiple comparison test (see Supplementary Table 15S).
