## Supplemental Tables for "Contrasting walking styles map to discrete neural substrates in the mouse brainstem"

**Table 1** | **CNO Dose and Time response titration**

**with Supplementary Figure 1**

| **VGaT Dose Response** | Time tested 1-2 hrs post CNO injection | **Swing Time** | | **One-way ANOVA** | **α 0.05** |
|---|---|---|---|---|---|
| **Group Z > 2 (slomo)** | **N** | **Mean (s)** | **SD (s)** | **F (DFn, DFd)** | **P value** |
| saline | 14 | 0.103 | 0.009 | F (3, 43) = 27.14 | P<0.0001 |
| 0.15 mg/kg CNO | 6 | 0.173 | 0.025 | | |
| 0.30 mg/kg CNO | 14 | 0.157 | 0.022 | | |
| 0.60 mg/kg CNO | 13 | 0.146 | 0.021 | | |

| **Dunnett's multiple comparisons test** | **Mean Diff.** | **95.00% CI of diff.** | **P Value, α 0.05** |
|---|---|---|---|
| Saline vs. 0.15 mg/kg | -0.07015 | -0.09294 to -0.04736 | 0.0001 |
| Saline vs. 0.3 mg/kg | -0.05342 | -0.07107 to -0.03577 | 0.0001 |
| Saline vs. 0.6 mg/kg | -0.04278 | -0.06077 to -0.02479 | 0.0001 |

| **VGaT Dose Response** | Time tested 1-2 hrs post CNO | **Stride velocity** | | **One-way ANOVA** | **α 0.05** |
|---|---|---|---|---|---|
| **Group Z > 2 (slomo)** | **N** | **Mean (cm/s)** | **SD (cm/s)** | **F (DFn, DFd)** | **P value** |
| saline | 14 | 11.29 | 4.31 | F (3, 43) = 6.283 | 0.001 |
| 0.15 mg/kg CNO | 6 | 5.33 | 1.87 | | |
| 0.30 mg/kg CNO | 14 | 7.16 | 3.19 | | |
| 0.60 mg/kg CNO | 13 | 6.94 | 3.00 | | |

| **Dunnett's multiple comparisons test** | **Mean Diff.** | **95.00% CI of diff.** | **P Value, α 0.05** |
|---|---|---|---|
| Saline vs. 0.15 mg/kg | 5.963 | 1.915 to 10.01 | 0.002 |
| Saline vs. 0.3 mg/kg | 4.133 | 0.9976 to 7.269 | 0.007 |
| Saline vs. 0.6 mg/kg | 4.358 | 1.163 to 7.554 | 0.005 |

| **VGluT2 Dose Response** | Time tested 1-2 hrs post CNO | **Swing Time** | | **One-way ANOVA** | **α 0.05** |
|---|---|---|---|---|---|
| **Group Z< -2 (shuffle)** | **N** | **Mean (s)** | **SD (s)** | **F (DFn, DFd)** | **P value** |
| saline | 20 | 0.098 | 0.017 | F (3, 59) = 10.96 | P<0.0001 |
| 0.15 mg/kg CNO | 9 | 0.076 | 0.011 | | |
| 0.30 mg/kg CNO | 20 | 0.075 | 0.015 | | |
| 0.60 mg/kg CNO | 14 | 0.079 | 0.008 | | |

| **Dunnett's multiple comparisons test** | **Mean Diff.** | **95.00% CI of diff.** | **P Value, α 0.05** |
|---|---|---|---|
| Saline vs. 0.15 mg/kg | 0.02248 | 0.00867 to 0.0363 | 0.0007 |
| Saline vs. 0.3 mg/kg | 0.02355 | 0.01267 to 0.03444 | 0.0001 |
| Saline vs. 0.6 mg/kg | 0.01908 | 0.007086 to 0.03107 | 0.0009 |

| **VGluT2 Dose Response** | Time tested 1-2 hrs post CNO | **Stride velocity** | | **One-way ANOVA** | **α 0.05** |
|---|---|---|---|---|---|
| **Group Z < -2 (shuffle)** | **N** | **Mean (cm/s)** | **SD (cm/s)** | **F (DFn, DFd)** | **P value** |
| saline | 20 | 10.95 | 1.97 | F (3, 59) = 11.19 | P<0.0001 |
| 0.15 mg/kg CNO | 9 | 7.27 | 1.59 | | |
| 0.30 mg/kg CNO | 20 | 7.38 | 2.17 | | |
| 0.60 mg/kg CNO | 14 | 8.92 | 2.59 | | |

| **Dunnett's multiple comparisons test** | **Mean Diff.** | **95.00% CI of diff.** | **P Value, α 0.05** |
|---|---|---|---|
| Saline vs. 0.15 mg/kg | 3.681 | 1.61 to 5.753 | 0.0002 |
| Saline vs. 0.3 mg/kg | 3.566 | 1.934 to 5.198 | 0.0001 |
| Saline vs. 0.6 mg/kg | 2.031 | 0.2325 to 3.829 | 0.02 |

| **VGaT Time Response** | Dose tested: 0.3mg/kg | **Swing Time (s)** | | | **Friedman test** | **α 0.05** |
|---|---|---|---|---|---|---|
| **Group Z > 2 (slomo)** | **N = 5** | **Median** | **Lower 95% CI** | **Upper 95 % CI** | **Friedman statistic** | **P value** |
| Baseline (0 h) | | 0.110 | 0.104 | 0.113 | 19.2 | 0.004 |
| 0.5 h | | 0.167 | 0.128 | 0.214 | | |
| 1 h | | 0.162 | 0.125 | 0.217 | | |
| 2 h | | 0.181 | 0.123 | 0.207 | | |
| 4 h | | 0.170 | 0.128 | 0.188 | | |
| 6 h | | 0.134 | 0.109 | 0.175 | | |
| 8 h | | 0.129 | 0.110 | 0.155 | | |

| **Dunn's multiple comparisons test** | **Rank sum diff.** | **P Value, α 0.05** |
|---|---|---|
| 0 vs. 30 min | -21 | 0.01 |
| 0 vs. 60 min | -23 | 0.005 |
| 0 vs. 120 min | -22 | 0.01 |
| 0 vs. 240 min | -19 | 0.03 |
| 0 vs. 360 min | -12 | 0.47 |
| 0 vs. 480 min | -8 | >0.99 |

| **VGaT Time Response** | Dose tested: 0.3mg/kg | **Stride velocity (cm/s)** | | | **Friedman test** | **α 0.05** |
|---|---|---|---|---|---|---|
| **Group Z > 2 (slomo)** | **N = 5** | **Median** | **Lower 95% CI** | **Upper 95 % CI** | **Friedman statistic** | **P value** |
| Baseline (0 h) | | 9.40 | 5.50 | 14.54 | 21.09 | 0.002 |
| 0.5 h | | 4.75 | 2.30 | 7.53 | | |
| 1 h | | 4.59 | 0.76 | 7.16 | | |
| 2 h | | 3.43 | 2.42 | 5.42 | | |
| 4 h | | 3.37 | 1.67 | 5.72 | | |
| 6 h | | 4.86 | 4.09 | 5.91 | | |
| 8 h | | 6.81 | 4.26 | 10.79 | | |

| **Dunn's multiple comparisons test** | **Rank sum diff.** | **P Value, α 0.05** |
|---|---|---|
| 0 vs. 30 min | 15 | 0.17 |
| 0 vs. 60 min | 22 | 0.008 |
| 0 vs. 120 min | 21 | 0.01 |
| 0 vs. 240 min | 23 | 0.005 |
| 0 vs. 360 min | 13 | 0.34 |
| 0 vs. 480 min | 4 | >0.99 |

**Table 2** Average stride velocity with saline versus CNO

with Figure 2b

| | | Saline | | CNO | | paired, 2 tailed t-test | α 0.05 |
|---|---|---|---|---|---|---|---|
| **VGaT Cohort** | **N** | **Mean (cm/s)** | **SD (cm/s)** | **Mean (cm/s)** | **SD (cm/s)** | **t(df)** | **p** |
| VGaT-ires- cre | 35 | 11.5 | 3.8 | 8.5 | 3.7 | t=4.965 df=34 | < 0.0001 |
| WT litter mates | 10 | 10.3 | 3.2 | 9.6 | 2.2 | t=0.6937 df=9 | 0.51 |
| **VGluT2 Cohort** | **N** | **Mean (cm/s)** | **SD (cm/s)** | **Mean (cm/s)** | **SD (cm/s)** | **t(df)** | **p** |
| VGluT2-ires-cre | 46 | 11.1 | 3.0 | 7.7 | 2.5 | t=6.657 df=45 | < 0.0001 |
| WT litter mates | 7 | 10.7 | 2.3 | 9.9 | 2.9 | t=1.518 df=6 | 0.18 |
| **SerT Cohort** | **N** | **Mean (cm/s)** | **SD (cm/s)** | **Mean (cm/s)** | **SD (cm/s)** | **t(df)** | **p** |
| SerT - cre | 18 | 12.0 | 2.9 | 12.3 | 3.5 | t=0.2782 df=17 | 0.78 |
| WT litter mates | 15 | 12.9 | 2.2 | 12.9 | 2.6 | t=0.005928 df=14 | 1.00 |

**Table 3** — **Gait metrics with saline versus CNO in VGaT sub-cohorts**

with Figure 3 and Supplementary Figure 2

### Footfall metrics

| | Group Z Swing Time >3 (slomo) | | Sum of squares F test (alpha 0.001) F (DFn, DFd) | p | | Saline Average | Saline SD | CNO Average | CNO SD | paired, 2 tailed t-test (alpha 0.05) t(df) | p |
|---|---|---|---|---|---|---|---|---|---|---|---|
| Figure 3 | N= 9 | Stride velocity (cm/s) | - | - | Stride velocity (cm/s) | 9.91 | 5.70 | 5.67 | 3.71 | t=3.33 df=8 | 0.01 |
| | | Stride length (mm) | 44.7 (2, 479) | <0.0001 | Stride length (mm) | 40.93 | 11.88 | 40.25 | 12.49 | t=0.09 df=8 | 0.93 |
| | n Strides: | Swing time (s) | 160.1 (2, 479) | <0.0001 | Swing time (s) | 0.10 | 0.02 | 0.16 | 0.05 | t=11.88 df=8 | <0.0001 |
| | saline: n = 320 | Stance time (s) | 2.83 (4, 519) | 0.02 | Stance time (s) | 0.43 | 0.33 | 0.83 | 0.69 | t=5.12 df=8 | 0.0009 |
| | CNO: n = 313 | Cadence (Hz) | 24.27 (2, 548) | <0.0001 | Cadence (Hz) | 2.35 | 0.97 | 1.38 | 0.78 | t=6.89 df=8 | 0.0001 |
| | | Swing speed (cm/s) | 42.4 (2, 522) | <0.0001 | Swing speed (cm/s) | 41.69 | 10.82 | 26.63 | 8.35 | t=8.07 df=8 | <0.0001 |

| | Group 3>Z Swing Time>2 (intermediate slomo) | | Sum of squares F test (alpha 0.001) F (DFn, DFd) | p | | Saline Average | Saline SD | CNO Average | CNO SD | paired, 2 tailed t-test (alpha 0.05) t(df) | p |
|---|---|---|---|---|---|---|---|---|---|---|---|
| | N=5 | Stride velocity (cm/s) | - | - | Stride velocity (cm/s) | 13.59 | 3.66 | 9.98 | 2.88 | t=5.99 df=4 | 0.004 |
| | | Stride length (mm) | 38.84 (2, 260) | <0.0001 | Stride length (mm) | 46.84 | 6.05 | 49.02 | 5.17 | t=2.489 df=4 | 0.07 |
| | n Strides: | Swing time (s) | 41.56 (2, 260) | <0.0001 | Swing time (s) | 0.11 | 0.01 | 0.15 | 0.02 | t=9.679 df=4 | 0.0006 |
| | saline: n = 186 | Stance time (s) | 5.171 (4, 252) | 0.0005 | Stance time (s) | 0.32 | 0.09 | 0.43 | 0.12 | t=2.659 df=4 | 0.06 |
| | CNO: n = 170 | Cadence (Hz) | 21.21 (2, 225) | <0.0001 | Cadence (Hz) | 2.84 | 0.41 | 1.99 | 0.36 | t=7.336 df=4 | 0.002 |
| | | Swing speed (cm/s) | 31.18 (2, 322) | <0.0001 | Swing speed (cm/s) | 42.68 | 5.72 | 34.30 | 5.52 | t=6.651 df=4 | 0.003 |

| | Group Z Swing Time <2 (non-slomo) | | Sum of squares F test (alpha 0.001) F (DFn, DFd) | p | | Saline Average | Saline SD | CNO Average | CNO SD | paired, 2 tailed t-test (alpha 0.05) t(df) | p |
|---|---|---|---|---|---|---|---|---|---|---|---|
| Supplementary Figure 2 | N= 21 | Stride velocity (cm/s) | - | - | Stride velocity (cm/s) | 11.63 | 3.84 | 9.45 | 3.73 | t=2.49 df=20 | 0.02 |
| | | Stride length (mm) | 2.62 (2, 1012) | 0.07 | Stride length (mm) | 43.76 | 5.75 | 42.43 | 6.84 | t=0.85 df=20 | 0.41 |
| | n Strides: | Swing time (s) | 3.91 (2, 1012) | 0.02 | Swing time (s) | 0.11 | 0.02 | 0.12 | 0.02 | t=1.94 df=20 | 0.07 |
| | saline: n = 671 | Stance time (s) | 1.47 (4, 1133) | 0.21 | Stance time (s) | 0.41 | 0.14 | 0.55 | 0.25 | t=2.54 df=20 | 0.02 |
| | CNO: n = 731 | Cadence (Hz) | 1.00 (2, 974) | 0.37 | Cadence (Hz) | 2.54 | 0.56 | 2.16 | 0.54 | t=2.85 df=20 | 0.01 |
| | | Swing speed (cm/s) | 0.34 (2, 1229) | 0.71 | Swing speed (cm/s) | 41.89 | 6.12 | 38.62 | 7.61 | t=2.32 df=20 | 0.03 |

| | WT control | | Sum of squares F test (alpha 0.001) F (DFn, DFd) | p | | Saline Average | Saline SD | CNO Average | CNO SD | paired, 2 tailed t-test (alpha 0.05) t(df) | p |
|---|---|---|---|---|---|---|---|---|---|---|---|
| | N= 10 | Stride velocity (cm/s) | - | - | Stride velocity (cm/s) | 10.33 | 3.24 | 9.62 | 2.15 | t=0.6937 df=9 | 0.51 |
| | | Stride length (mm) | 1.84 (2, 547) | 0.16 | Stride length (mm) | 44.27 | 7.31 | 42.40 | 4.99 | t=0.6455 df=9 | 0.53 |
| | n Strides: | Swing time (s) | 3.00 (2, 547) | 0.05 | Swing time (s) | 0.10 | 0.01 | 0.10 | 0.01 | t=1.019 df=9 | 0.34 |
| | saline: n = 363 | Stance time (s) | 1.98 (4, 597) | 0.10 | Stance time (s) | 0.47 | 0.12 | 0.46 | 0.13 | t=0.2033 df=9 | 0.84 |
| | CNO: n = 346 | Cadence (Hz) | 3.13 (2, 490) | 0.04 | Cadence (Hz) | 2.27 | 0.41 | 2.23 | 0.40 | t=0.2983 df=9 | 0.77 |
| | | Swing speed (cm/s) | 3.06 (2, 614) | 0.05 | Swing speed (cm/s) | 43.82 | 8.28 | 43.84 | 4.21 | t=0.01127 df=9 | 0.99 |

### Temporal coupling

| | | Comparison | Saline r | CNO r | Watson (alpha 0.05) |
|---|---|---|---|---|---|
| Figure 3 | Group Z>3 (slomo) | Back left-Back right | 0.59 | 0.79 | 0.62 |
| | Group 3>Z>2 (intermediate) | Back left-Back right | 0.52 | 0.72 | 0.98 |
| Supplementary Figure 2 | Group Z<2 (non-slomo) | Back left-Back right | 0.67 | 0.68 | 0.49 |
| | WT control | Back left-Back right | 0.62 | 0.58 | 0.71 |

### Postural metrics

| | Z swingtime >3 (slomo) | N | Mean | SD | Mean | SD | paired, 2 tailed t-test (alpha 0.05) t, df | p |
|---|---|---|---|---|---|---|---|---|
| Figure 3 | **Height start swing** | | | | | | | |
| | Iliac crest to floor (mm) | 9 | 26.4 | 3.0 | 28.2 | 2.8 | t=3.113 df=8 | 0.01 |
| | Tail base to floor (mm) | 9 | 12.6 | 3.5 | 16.5 | 2.7 | t=8.571 df=8 | <0.0001 |
| | **Height end swing** | | | | | | | |
| | Iliac crest to floor (mm) | 9 | 27.4 | 28.3 | 12.3 | 15.4 | t=1.306 df=8 | 0.23 |
| | Tail base to floor (mm) | 9 | 3.5 | 2.0 | 4.0 | 2.4 | t=3.87 df=8 | 0.005 |

| | Z swing time<2 (non-slomo) | N | Mean | SD | Mean | SD | t, df | p |
|---|---|---|---|---|---|---|---|---|
| Supplementary Figure 2 | **Height start swing** | | | | | | | |
| | Iliac crest to floor (mm) | 21 | 27.6 | 4.2 | 28.5 | 2.5 | t=1.284 df=20 | 0.21 |
| | Tail base to floor (mm) | 21 | 14.0 | 4.4 | 16.2 | 2.6 | t=2.63 df=20 | 0.02 |
| | **Height end swing** | | | | | | | |
| | Iliac crest to floor (mm) | 21 | 28.2 | 3.9 | 28.8 | 2.3 | t=0.8365 df=20 | 0.41 |
| | Tail base to floor (mm) | 21 | 13.3 | 4.3 | 15.6 | 3.0 | t=2.478 df=20 | 0.02 |

**Supplementary Table 4**
with Supplementary Figure 3
with Supplementary Figure 3

### A: Anterior Tibial EMG in VGaT sub-cohorts

| Walk | | Sub-cohort (Z Swing Time) | N | Saline Average | SD | CNO Average | SD | Paired, two-tailed t test t, df | (alpha=0.05) p |
|---|---|---|---|---|---|---|---|---|---|
| Duration | burst (sec) | Z>2 (slomo) | 6 | 0.07 | 0.022 | 0.16 | 0.04 | t=6.475 df=5 | 0.001 |
| | | Z<2 (non-slomo) | 10 | 0.07 | 0.014 | 0.09 | 0.05 | t=2.076 df=9 | 0.07 |
| | | WT | 4 | 0.09 | 0.037 | 0.10 | 0.03 | t=1.376 df=3 | 0.26 |
| | interburst (sec) | Z>2 (slomo) | 6 | 0.37 | 0.157 | 1.82 | 1.14 | t=3.445 df=5 | 0.02 |
| | | Z<2 (non-slomo) | 10 | 0.38 | 0.126 | 0.74 | 0.38 | t=3.316 df=9 | 0.01 |
| | | WT | 4 | 0.45 | 0.231 | 0.49 | 0.13 | t=0.4227 df=3 | 0.70 |
| Peak amplitude (Normalized) | burst | Z>2 (slomo) | 6 | 0.43 | 0.19 | 0.39 | 0.20 | t=1.678 df=5 | 0.15 |
| | | Z<2 (non-slomo) | 10 | 0.35 | 0.14 | 0.30 | 0.08 | t=0.7027 df=9 | 0.50 |
| | | WT | 4 | 0.34 | 0.03 | 0.34 | 0.02 | t=0.04207 df=3 | 0.97 |
| | interburst | Z>2 (slomo) | 6 | 0.54 | 0.26 | 0.29 | 0.11 | t=3.04 df=5 | 0.03 |
| | | Z<2 (non-slomo) | 10 | 0.46 | 0.19 | 0.37 | 0.18 | t=2.721 df=9 | 0.02 |
| | | WT | 4 | 0.51 | 0.19 | 0.39 | 0.12 | t=3.05 df=3 | 0.06 |
| Integral (Normalized) | burst | Z>2 (slomo) | 6 | 0.46 | 0.17 | 0.66 | 0.28 | t=3.303 df=5 | 0.02 |
| | | Z<2 (non-slomo) | 10 | 0.39 | 0.13 | 0.40 | 0.11 | t=0.1362 df=9 | 0.89 |
| | | WT | 4 | 0.47 | 0.18 | 0.50 | 0.14 | t=1.003 df=3 | 0.39 |
| | interburst | Z>2 (slomo) | 6 | 1.27 | 0.85 | 3.41 | 2.89 | t=2.38 df=5 | 0.06 |
| | | Z<2 (non-slomo) | 10 | 1.39 | 0.77 | 2.04 | 1.07 | t=2.549 df=9 | 0.03 |
| | | WT | 4 | 1.50 | 1.09 | 1.60 | 0.93 | t=0.3774 df=3 | 0.73 |

| Swim | | Sub-cohort (Z Swing Time) | N | Saline Average | SD | CNO Average | SD | Paired, two-tailed t test t, df | (alpha=0.05) p |
|---|---|---|---|---|---|---|---|---|---|
| Duration | burst (sec) | Z>2 (slomo) | 6 | 0.06 | 0.012 | 0.07 | 0.0 | t=1.95 df=5 | 0.11 |
| | | Z<2 (non-slomo) | 10 | 0.06 | 0.014 | 0.06 | 0.0 | t=0.4518 df=9 | 0.66 |
| | | WT | 4 | 0.07 | 0.021 | 0.05 | 0.0 | t=1.034 df=3 | 0.38 |
| | interburst (sec) | Z>2 (slomo) | 6 | 0.11 | 0.017 | 0.14 | 0.0 | t=2.313 df=5 | 0.07 |
| | | Z<2 (non-slomo) | 10 | 0.11 | 0.029 | 0.12 | 0.0 | t=1.591 df=9 | 0.15 |
| | | WT | 4 | 0.11 | 0.024 | 0.11 | 0.0 | t=0.06607 df=3 | 0.95 |

| | | Sub-cohort (Z Swing Time) | N | Saline Median | Lower 95% CI | Upper 95% CI | CNO Median | Lower 95% CI | Wilcoxon two-tailed paired ranks test Upper 95% CI | W | p |
|---|---|---|---|---|---|---|---|---|---|---|---|
| Frequency | (Hz) | Z>2 (slomo) | 6 | 6.0 | 5.4 | 6.9 | 5.2 | 4.0 | 6.2 | -21 | 0.03 |
| | | Z<2 (non-slomo) | 10 | 5.9 | 5.7 | 6.5 | 5.7 | 5.3 | 6.2 | -33 | 0.11 |
| | | WT | 4 | 6.5 | 4.2 | 7.9 | 6.0 | 5.0 | 7.6 | 4 | 0.63 |

### B: Gastrocnemius EMG in VGaT sub-cohorts

| Walk | | Sub-cohort (Z Swing Time) | N | Saline Median | Lower 95% CI | Upper 95% CI | CNO Median | Lower 95% CI | Wilcoxon two-tailed paired ranks test Upper 95% CI | W | p |
|---|---|---|---|---|---|---|---|---|---|---|---|
| Duration | burst (sec) | Z>2 (slomo) | 4 | 0.24 | 0.15 | 0.35 | 0.90 | 0.28 | 1.88 | 10 | 0.13 |
| | | Z<2 (non-slomo) | 4 | 0.33 | 0.10 | 0.66 | 0.49 | 0.25 | 0.77 | 10 | 0.13 |
| | interburst (sec) | Z>2 (slomo) | 4 | 0.09 | 0.04 | 0.15 | 0.13 | -0.06 | 0.48 | 10 | 0.13 |
| | | Z<2 (non-slomo) | 4 | 0.08 | 0.05 | 0.12 | 0.11 | 0.01 | 0.24 | 6 | 0.38 |
| Peak amplitude (Normalized) | burst | Z>2 (slomo) | 4 | 0.30 | 0.15 | 0.41 | 0.22 | 0.10 | 0.38 | -8 | 0.25 |
| | | Z<2 (non-slomo) | 4 | 0.22 | 0.10 | 0.32 | 0.19 | 0.00 | 0.42 | -2 | 0.88 |
| | interburst | Z>2 (slomo) | 4 | 0.23 | 0.17 | 0.31 | 0.21 | 0.12 | 0.28 | -8 | 0.25 |
| | | Z<2 (non-slomo) | 4 | 0.15 | 0.00 | 0.36 | 0.15 | 0.02 | 0.29 | -4 | 0.63 |
| Integral (Normalized) | burst | Z>2 (slomo) | 4 | 1.06 | 0.68 | 1.27 | 2.87 | 0.52 | 5.86 | 10 | 0.13 |
| | | Z<2 (non-slomo) | 4 | 1.09 | 0.23 | 2.20 | 1.52 | 0.27 | 2.86 | 8 | 0.25 |
| | interburst | Z>2 (slomo) | 4 | 0.27 | 0.17 | 0.38 | 0.41 | -0.02 | 1.23 | 10 | 0.13 |
| | | Z<2 (non-slomo) | 4 | 0.25 | 0.12 | 0.37 | 0.34 | 0.00 | 0.68 | 4 | 0.63 |

| Swim | | Sub-cohort (Z Swing Time) | N | Saline Median | Lower 95% CI | Upper 95% CI | CNO Median | Lower 95% CI | Upper 95% CI | W | p |
|---|---|---|---|---|---|---|---|---|---|---|---|
| Duration | burst | Z > 2 | 4 | 0.05 | 0.02 | 0.10 | 0.08 | 0.04 | 0.14 | 10 | 0.13 |
| | | Z < -2 | 4 | 0.05 | 0.03 | 0.07 | 0.06 | 0.04 | 0.08 | 6 | 0.38 |
| | interburst | Z > 2 | 4 | 0.10 | 0.02 | 0.22 | 0.09 | 0.03 | 0.16 | -10 | 0.13 |
| | | Z < 2 | 4 | 0.11 | 0.05 | 0.16 | 0.11 | 0.10 | 0.13 | 4 | 0.63 |

**Supplementary Table 5** **Correlation between Swing Time and TA EMG metrics**

**with Supplementary Figs. 3 and 5**

| | Group | N | Pearson correlation | r | 95% confidence interval | R squared | P (two-tailed)* |
|---|---|---|---|---|---|---|---|
| Supplementary Fig. 3 | VGaT | 16 | Z swing time versus Z EMG burst duration | 0.63 | 0.1911 to 0.8567 | 0.39 | 0.005 |
| | VGaT | 16 | Z swing time versus Z EMG burst amplitude | 0.24 | -0.2914 to 0.6568 | 0.057 | 0.37 |
| Supplementary Fig. 5 | VGlut2 | 13 | Z swing time versus Z EMG burst amplitude | -0.34 | -0.7523 to 0.2555 | 0.12 | 0.25 |
| | VGlut2 | 13 | Z swing time versus Z EMG burst duration | 0.32 | -0.2798 to 0.7407 | 0.10 | 0.14 |

**Supplementary Table 6** | **Motor tests in VGaT sub-cohorts**

**Supplementary to Results**

| Rotarod | Sub-cohort (Z Swing Time) | N | Saline Median | Lower 95% CI | Upper 95% CI | CNO Median | Lower 95% CI | Upper 95% CI | Wilcoxon matched-pairs signed rank test, tw W | (alpha=0.05) p |
|---|---|---|---|---|---|---|---|---|---|---|
| Latency to fall (sec) | Group Z>3 (slomo) | 9 | 110 | 91 | 134 | 77 | 61 | 89 | -45 | 0.004 |
| | Group 3>Z>2 (intermediate) | 5 | 96 | 45 | 156 | 86 | 39 | 121 | -10 | 0.13 |
| | Group Z<2 (non-slomo) | 21 | 108 | 103 | 125 | 88 | 84 | 111 | -151 | 0.007 |
| | WT control | 10 | 112 | 102 | 121 | 116 | 101 | 130 | 19 | 0.38 |

| Beam | Sub-cohort (Z Swing Time) | N | Median | Lower 95% CI | Upper 95% CI | Median | Lower 95% CI | Upper 95% CI | W | p |
|---|---|---|---|---|---|---|---|---|---|---|
| # Slips and misses | Group Z>3 (slomo) | 9 | 8 | 2.5 | 13.7 | 4 | 0.2 | 12.7 | -8 | 0.58 |
| | Group 3>Z>2 (intermediate) | 5 | 5 | 2.4 | 9.6 | 6 | -10.8 | 36.0 | -1 | >0.9999 |
| | Group Z<2 (non-slomo) | 21 | 5 | 3.5 | 12.3 | 4 | 3.3 | 9.6 | -32 | 0.53 |
| | WT control | 10 | 6 | 3.0 | 19.4 | 10 | 4.8 | 15.8 | 6 | 0.75 |

| Ladder | Sub-cohort (Z Swing Time) | N | Median | Lower 95% CI | Upper 95% CI | Median | Lower 95% CI | Upper 95% CI | W | p |
|---|---|---|---|---|---|---|---|---|---|---|
| # Slips and misses | Group Z>3 (slomo) | 9 | 0 | -0.11 | 1.00 | 0 | -0.19 | 1.53 | 4 | 0.81 |
| | Group 3>Z>2 (intermediate) | 5 | 0 | -1.02 | 3.42 | 1 | 0.16 | 2.24 | -1 | >0.9999 |
| | Group Z<2 (non-slomo) | 21 | 1 | 0.44 | 1.75 | 1 | 0.33 | 1.10 | -30 | 0.32 |
| | WT control | 10 | 0 | -0.20 | 1.60 | 0.5 | -0.02 | 1.82 | 7 | 0.80 |

| Supplementary Table 7 | Subanalysis: Asymmetry | Does unilateral or bilateral activation drive results in VGaT sub-cohorts? |
|---|---|---|

with results text and Suppplementary Fig. 10

**Note: Z>3 slomo group is exclusively bilateral; all unilateral cases resulted in Z<2.**

| Saline vs CNO of bilateral Z Swing time<2 cases (non-slomo) | | Sum of squares F test (alpha 0.001)<br>F (DFn, DFd) | p |
|---|---|---|---|
| N= 15 | Stride velocity (cm/s) | - | - |
| | Stride length (mm) | 2.87 (2, 725) | 0.06 |
| n Strides: | Swing time (s) | 2.69 (2, 725) | 0.07 |
| saline: n = 458 | Stance time (s) | 0.91 (4, 810) | 0.46 |
| CNO: n = 523 | Cadence (Hz) | 1.09 (2, 704) | 0.34 |
| | Swing speed (mm) | 1.07 (2, 878) | 0.34 |

| Saline vs CNO of unilateral Z Swing time<2 cases (non-slomo; target in slomo hotspot) | | Sum of squares F test (alpha 0.001)<br>F (DFn, DFd) | p |
|---|---|---|---|
| N= 6 | Stride velocity (cm/s) | - | - |
| | Stride length (mm) | 2.43 (2, 283) | 0.09 |
| n Strides: | Swing time (s) | 1.84 (2, 283) | 0.16 |
| saline: n = 213 | Stance time (s) | 2.73 (4, 315) | 0.03 |
| CNO: n = 208 | Cadence (Hz) | 2.77 (2, 266) | 0.06 |
| | Swing speed (mm) | 0.32 (2, 347) | 0.72 |

| Temporal coupling<br>Sub analysis: unilateral and bilateral | | saline<br>r | CNO<br>r | Watson (alpha 0.05) |
|---|---|---|---|---|
| Group Z<2 swingtime, bilateral Sal vs CNO | Back left-Back right | 0.67 | 0.69 | 0.12 |
| Group Z<2 swingtime, unilateral Sal vs CN | Back left-Back right | 0.69 | 0.66 | 0.37 |

| Supplementary Fig. 10 | Subanalysis: Asymmetry | Gait metrics in ipsi- or contralateral hindlimbs: comparison between saline versus CNO |
|---|---|---|

| Ipsilateral hindlimb Saline versus CNO<br>unilateral Z Swing time<2 cases, target in slomo hotspot | | Sum of squares F test (alpha 0.001)<br>F (DFn, DFd) | p |
|---|---|---|---|
| N = 6 | Stride velocity (cm/s) | - | - |
| | Stride length (mm) | 1.095 (2, 140) | 0.34 |
| Strides | Swing time (s) | 0.2479 (2, 140) | 0.78 |
| saline n = 107 | Stance time (s) | - | - |
| CNO n =105 | Cadence (Hz) | 1.584 (2, 129) | 0.21 |
| | Swing speed (mm) | 0.6532 (2, 176) | 0.52 |

| Contralateral hindlimb Saline versus CNO<br>unilateral Z Swing time<2 cases, target in slomo hotspot | | Sum of squares F test (alpha 0.001)<br>F (DFn, DFd) | p |
|---|---|---|---|
| N = 6 | Stride velocity (cm/s) | - | - |
| | Stride length (mm) | 1.493 (2, 139) | 0.23 |
| Strides | Swing time (s) | 2.541 (2, 139) | 0.08 |
| saline n = 106 | Stance time (s) | - | - |
| CNO n = 105 | Cadence (Hz) | 1.272 (2, 135) | 0.28 |
| | Swing speed (mm) | 1.352 (2, 169) | 0.26 |

| Supplementary Fig. 10 | Subanalysis: Asymmetry | Gait metrics in saline or CNO condition: comparison between ipsi- versus contralateral hindlimbs |
|---|---|---|

| Saline<br>ipsi- vs contralateral hindlimb of unilateral Z Swing time<2 cases, target in slomo hotspot | | Sum of squares F test (alpha 0.001)<br>F (DFn, DFd) | p |
|---|---|---|---|
| N = 6 | Stride velocity (cm/s) | - | - |
| | Stride length (mm) | 0.1399 (2, 138) | 0.87 |
| Strides | Swing time (s) | 2.283 (2, 138) | 0.11 |
| ipsilateral n = 71 | Stance time (s) | - | - |
| contralateral n = 71 | Cadence (Hz) | 0.5874 (2, 136) | 0.56 |
| | Swing speed (mm) | 1.477 (2, 172) | 0.23 |

| CNO<br>ipsi- vs contralateral hindlimb of unilateral Z Swing time<2 cases, target in slomo hotspot | | Sum of squares F test (alpha 0.001)<br>F (DFn, DFd) | p |
|---|---|---|---|
| N = 6 | Stride velocity (cm/s) | - | - |
| | Stride length (mm) | 0.1938 (2, 141) | 0.82 |
| Strides | Swing time (s) | 0.3329 (2, 141) | 0.72 |
| ipsilateral n = 73 | Stance time (s) | - | - |
| contralateral n = 72 | Cadence (Hz) | 0.6518 (2, 128) | 0.52 |
| | Swing speed (mm) | 1.306 (2, 173) | 0.27 |

**Subanalysis: Motor tests in bilateral and unilateral cases**

| Rotarod | Sub-cohort (laterality and Z Swing Time) | N | Saline<br>Median | Lower 95% CI | Upper 95% CI | CNO<br>Median | Lower 95% CI | Upper 95% CI | Wilcoxon matched-pairs signed rank test, two-ta<br>W | (alpha=0.05)<br>p |
|---|---|---|---|---|---|---|---|---|---|---|
| Latency to fall (sec) | Group Z swingtime <2 bilateral | 15 | 107.30 | 97.30 | 120.40 | 81.00 | 75.92 | 100.60 | -86 | 0.01 |
| | Group Z swingtime<2 unilateral | 6 | 127.70 | 95.34 | 155.80 | 124.70 | 86.41 | 154.30 | -9 | 0.44 |
| **Beam** | **Sub-cohort (laterality and Z Swing Time)** | **N** | **Median** | **Lower 95% CI** | **Upper 95% CI** | **Median** | **Lower 95% CI** | **Upper 95% CI** | **W** | **p** |
| # Slips and misses | Group Z swingtime <2 bilateral | 15 | 2.00 | 0.26 | 11.08 | 4.00 | 3.30 | 11.90 | 37 | 0.26 |
| | Group Z swingtime <2 unilateral | 6 | 15.00 | 5.94 | 21.06 | 3.00 | 0.07 | 6.93 | -15 | 0.06 |
| **Ladder** | **Sub-cohort (laterality and Z Swing Time)** | **N** | **Median** | **Lower 95% CI** | **Upper 95% CI** | **Median** | **Lower 95% CI** | **Upper 95% CI** | **W** | **p** |
| # Slips and misses | Group Z swingtime <2 bilateral | 15 | 0.00 | 0.22 | 2.04 | 1.00 | 0.24 | 1.22 | -17 | 0.48 |
| | Group Z swingtime <2 unilateral | 6 | 1.00 | 0.06 | 1.94 | 0.50 | -0.19 | 1.52 | -3 | 0.50 |

**Supplementary Table 8** — with Figure 4 and Supplementary Figure 4

### Gait metrics with saline versus CNO in VGluT2 subcohorts

#### Footfall metrics

with Figure 4

**Group Z Swing Time<-3 (shuffle)**

| | | Sum of squares F test (alpha 0.001) F (DFn, DFd) | p | | Saline Average | Saline SD | CNO Average | CNO SD | paired, 2 tailed t-test (alpha 0.05) t(df) | p |
|---|---|---|---|---|---|---|---|---|---|---|
| N = 15 | Stride velocity (cm/s) | - | - | Stride velocity (cm/s) | 10.42 | 2.03 | 7.05 | 2.60 | t=3.777 df=14 | 0.002 |
| | Stride length (mm) | 84.18 (2, 906) | <0.0001 | Stride length (mm) | 40.74 | 5.47 | 28.22 | 7.09 | t=6.325 df=14 | <0.0001 |
| n Strides: | Swing time (s) | 63.13 (2, 906) | <0.0001 | Swing time (s) | 0.11 | 0.02 | 0.07 | 0.02 | t=11.87 df=14 | <0.0001 |
| saline n = 491 | Stance time (s) | 20.28 (4, 841) | <0.0001 | Stance time (s) | 0.36 | 0.06 | 0.43 | 0.12 | t=1.978 df=14 | 0.07 |
| CNO n = 631 | Cadence (Hz) | 49.5 (2, 886) | <0.0001 | Cadence (Hz) | 2.49 | 0.26 | 2.42 | 0.42 | t=0.4978 df=14 | 0.63 |
| | Swing speed (cm/s) | 176.9 (2, 1015) | <0.0001 | Swing speed (cm/s) | 40.19 | 6.62 | 38.57 | 7.56 | t=0.812 df=14 | 0.43 |

**Group -2>Z Swing Time >-3 (intermediate shuffle)**

| | | Sum of squares F test (alpha 0.001) F (DFn, DFd) | p | | Saline Average | Saline SD | CNO Average | CNO SD | paired, 2 tailed t-test (alpha 0.05) t(df) | p |
|---|---|---|---|---|---|---|---|---|---|---|
| N= 16 | Stride velocity (cm/s) | - | - | Stride velocity (cm/s) | 10.73 | 2.14 | 7.29 | 1.55 | t=6.11 df=15 | <0.0001 |
| | Stride length (mm) | 26.67 (2, 1120) | <0.0001 | Stride length (mm) | 40.50 | 5.91 | 30.61 | 2.77 | t=8.79 df=15 | <0.0001 |
| n Strides: | Swing time (s) | 5.26 (2, 1120) | 0.005 | Swing time (s) | 0.10 | 0.02 | 0.08 | 0.01 | t=7.98 df=15 | <0.0001 |
| saline: n = 656 | Stance time (s) | 9.43 (4, 1148) | <0.0001 | Stance time (s) | 0.35 | 0.07 | 0.45 | 0.16 | t=2.69 df=15 | 0.02 |
| CNO: n = 675 | Cadence (Hz) | 11.16 (2, 1070) | <0.0001 | Cadence (Hz) | 2.61 | 0.37 | 2.34 | 0.48 | t=2.10 df=15 | 0.05 |
| | Swing speed (cm/s) | 80.38 (2, 1196) | <0.0001 | Swing speed (cm/s) | 42.57 | 4.63 | 38.21 | 5.15 | t=6.28 df=15 | <0.0001 |

with Supplementary Figure 4

**Group Z Swing Time >-2 (non-shuffle)**

| | | Sum of squares F test (alpha 0.001) F (DFn, DFd) | p | | Saline Average | Saline SD | CNO Average | CNO SD | paired, 2 tailed t-test (alpha 0.05) t(df) | p |
|---|---|---|---|---|---|---|---|---|---|---|
| N = 14 | Stride velocity (cm/s) | - | - | Stride velocity (cm/s) | 12.32 | 3.82 | 8.90 | 3.06 | t=3.612 df=13 | 0.003 |
| | Stride length (mm) | 8.57 (2, 1009) | 0.0002 | Stride length (mm) | 41.78 | 6.22 | 33.04 | 7.50 | t=5.823 df=13 | <0.0001 |
| n Strides: | Swing time (s) | 2.23 (2, 1009) | 0.11 | Swing time (s) | 0.10 | 0.10 | 0.09 | 0.01 | t=8.376 df=13 | <0.0001 |
| saline n = 575 | Stance time (s) | 3.14 (4, 1126) | 0.01 | Stance time (s) | 0.34 | 0.15 | 0.40 | 0.11 | t=1.732 df=13 | 0.11 |
| CNO n = 806 | Cadence (Hz) | 2.77 (2, 968) | 0.06 | Cadence (Hz) | 2.85 | 0.59 | 2.59 | 0.49 | t=1.47 df=13 | 0.17 |
| | Swing speed (cm/s) | 41.28 (2, 1161) | <0.0001 | Swing speed (cm/s) | 44.86 | 5.79 | 38.61 | 8.49 | t=3.572 df=13 | 0.003 |

**WT control**

| | | Sum of squares F test (alpha 0.001) F (DFn, DFd) | p | | Saline Average | Saline SD | CNO Average | CNO SD | paired, 2 tailed t-test (alpha 0.05) t(df) | p |
|---|---|---|---|---|---|---|---|---|---|---|
| N= 6 | Stride velocity (cm/s) | - | - | Stride velocity (cm/s) | 11.24 | 1.94 | 10.44 | 2.75 | t=1.33 df=5 | 0.24 |
| | Stride length (mm) | 0.99 (2, 246) | 0.37 | Stride length (mm) | 48.58 | 6.76 | 45.47 | 7.36 | t=2.90 df=5 | 0.03 |
| n Strides: | Swing time (s) | 0.13 (2, 246) | 0.88 | Swing time (s) | 0.11 | 0.02 | 0.11 | 0.02 | t=0.27 df=5 | 0.80 |
| saline: n = 160 | Stance time (s) | 2.56 (4, 248) | 0.04 | Stance time (s) | 0.38 | 0.06 | 0.43 | 0.12 | t=1.24 df=5 | 0.27 |
| CNO: n = 160 | Cadence (Hz) | 2.97 (2, 187) | 0.05 | Cadence (Hz) | 2.27 | 0.21 | 2.19 | 0.42 | t=0.56 df=5 | 0.60 |
| | Swing speed (cm/s) | 1.47 (2, 283) | 0.23 | Swing speed (cm/s) | 44.11 | 6.36 | 42.35 | 5.57 | t=2.32 df=5 | 0.07 |

#### Temporal coupling

| | Swing Time | Comparison | saline r | CNO r | Watson (alpha 0.05) |
|---|---|---|---|---|---|
| with Figure 4 | Group Z swingtime <-3 (shuffle) | Back left-Back right | 0.74 | 0.60 | 0.00002 |
| | Group -3>Z swingtime>-2 (intermediate) | Back left-Back right | 0.73 | 0.60 | 0.002 |
| with Supplementary Figure 4 | Group Z swingtime > -2 (non-shuffle) | Back left-Back right | 0.69 | 0.60 | 0.03 |
| | WT control | Back left-Back right | 0.80 | 0.77 | 0.22 |

#### Postural metrics

| | | N | Mean | SD | Mean | SD | paired, 2 tailed t-test (alpha 0.05) t, df | p |
|---|---|---|---|---|---|---|---|---|
| with Figure 4 | **Group Z swing time<-3 (shuffle)** | | | | | | | |
| | **Height start swing** | | | | | | | |
| | Iliac crest to floor (mm) | 15 | 26.9 | 1.6 | 24.4 | 2.2 | t=4.131 df=14 | 0.001 |
| | Tail base to floor (mm) | 15 | 15.0 | 2.2 | 10.2 | 3.1 | t=5.514 df=14 | <0.0001 |
| | **Height end swing** | | | | | | | |
| | Iliac crest to floor (mm) | 15 | 27.5 | 1.7 | 24.1 | 2.0 | t=5.823 df=14 | <0.0001 |
| | Tail base to floor (mm) | 15 | 15.0 | 2.4 | 9.7 | 2.4 | t=7.008 df=14 | <0.0001 |
| | **Group Z swing time >-2 (non-shuffle)** | **N** | **Mean** | **SD** | **Mean** | **SD** | **t, df** | **p** |
| with Supplementary Figure 4 | **Height start swing** | | | | | | | |
| | Iliac crest to floor (mm) | 14 | 29.3 | 3.4 | 28.8 | 2.2 | t=0.6227 df=13 | 0.54 |
| | Tail base to floor (mm) | 14 | 14.5 | 4.1 | 12.8 | 3.0 | t=1.696 df=13 | 0.11 |
| | **Height end swing** | | | | | | | |
| | Iliac crest to floor (mm) | 14 | 29.6 | 3.8 | 28.6 | 2.0 | t=1.274 df=13 | 0.23 |
| | Tail base to floor (mm) | 14 | 14.3 | 4.3 | 12.0 | 3.1 | t=2.198 df=13 | 0.047 |

**Supplementary Table 9** | **Motor tests in VGluT2 sub-cohorts**

**Supplementary to Results**

| | | | **Saline** | | | **CNO** | | | **Wilcoxon matched-\| (alpha=0.05)** | |
|---|---|---|---|---|---|---|---|---|---|---|
| **Rotarod** | **Group (Z Swing Time)** | **N** | **Median** | **Lower 95% CI** | **Upper 95% CI** | **Median** | **Lower 95% CI** | **Upper 95% CI** | **W** | **p** |
| Latency to fall (sec) | Group Z<-3 (shuffle) | 10 | 107.50 | 101.10 | 120.00 | 116.30 | 100.10 | 136.70 | 25 | 0.23 |
| | Group -3<Z<-2 (intermediate) | 10 | 118.50 | 94.55 | 132.40 | 113.80 | 100.00 | 125.60 | 1 | >0.9999 |
| | Group Z>-2 (non-shuffle) | 13 | 122.30 | 89.43 | 140.00 | 117.00 | 76.89 | 140.70 | -11 | 0.74 |
| | WT control | 6 | 123.00 | 99.78 | 136.20 | 129.50 | 103.00 | 160.90 | 15 | 0.16 |
| **Beam** | **Group (Z Swing Time)** | **N** | **Median** | **Lower 95% CI** | **Upper 95% CI** | **Median** | **Lower 95% CI** | **Upper 95% CI** | **W** | **p** |
| # Slips and misses | Group Z<-3 (shuffle) | 10 | 7.00 | 3.12 | 12.48 | 4.50 | 1.32 | 23.08 | 15 | 0.33 |
| | Group -3<Z<-2 (intermediate) | 10 | 14.00 | 4.97 | 21.83 | 9.50 | 4.13 | 19.67 | -10 | 0.54 |
| | Group Z>-2 (non-shuffle) | 13 | 12.00 | 9.33 | 28.83 | 9.00 | 3.67 | 35.41 | -14 | 0.61 |
| | WT control | 6 | 18.00 | 0.20 | 50.14 | 18.00 | -4.71 | 48.71 | -5 | 0.66 |
| **Ladder** | **Group (Z Swing Time)** | **N** | **Median** | **Lower 95% CI** | **Upper 95% CI** | **Median** | **Lower 95% CI** | **Upper 95% CI** | **W** | **p** |
| # Slips and misses | Group Z<-3 (shuffle) | 10 | 1.50 | 0.39 | 1.81 | 2.00 | 0.69 | 3.31 | 14 | 0.19 |
| | Group -3<Z<-2 (intermediate) | 10 | 2.00 | 1.18 | 3.42 | 3.00 | 1.66 | 4.74 | 16 | 0.34 |
| | Group Z>-2 (non-shuffle) | 13 | 2.00 | 1.01 | 2.99 | 1.00 | 0.21 | 3.02 | -18 | 0.43 |
| | WT control | 6 | 2.50 | 0.49 | 5.17 | 2.00 | 1.13 | 2.21 | -7 | 0.38 |

**Supplementary Table 10**

with Supplementary Figure 5

Supplementary Figure 5

### Anterior Tibial EMG in VGluT2 subcohorts

| Walk | | Group (Z Swing Time) | N | Saline Average | SD | CNO Average | SD | Paired, two-tailed t test t, df | (alpha=0.05) p |
|---|---|---|---|---|---|---|---|---|---|
| **Duration** | burst (sec) | Z < -2 (shuffle) | 7 | 0.08 | 0.02 | 0.05 | 0.02 | t=4.324 df=6 | **0.005** |
| | | Z > -2 (non-shuffle) | 6 | 0.07 | 0.02 | 0.05 | 0.02 | t=3.367 df=5 | 0.02 |
| | interburst (sec) | Z < -2 (shuffle) | 7 | 0.34 | 0.11 | 0.39 | 0.11 | t=1.405 df=6 | 0.21 |
| | | Z > -2 (non-shuffle) | 6 | 0.33 | 0.08 | 0.36 | 0.06 | t=1.23 df=5 | 0.27 |
| **Peak amplitude** | burst | Z < -2 (shuffle) | 7 | 0.31 | 0.07 | 0.40 | 0.11 | t=3.307 df=6 | 0.02 |
| (Normalized) | | Z > -2 (non-shuffle) | 6 | 0.26 | 0.14 | 0.36 | 0.16 | t=3.044 df=5 | 0.03 |
| | interburst | Z < -2 (shuffle) | 7 | 0.44 | 0.15 | 0.62 | 0.20 | t=3.043 df=6 | 0.02 |
| | | Z > -2 (non-shuffle) | 6 | 0.43 | 0.23 | 0.71 | 0.39 | t=2.652 df=5 | 0.05 |
| **Integral** | burst | Z < -2 (shuffle) | 7 | 0.37 | 0.09 | 0.33 | 0.12 | t=1.642 df=6 | 0.15 |
| (Normalized) | | Z > -2 (non-shuffle) | 6 | 0.28 | 0.17 | 0.30 | 0.17 | t=1.142 df=5 | 0.31 |
| | interburst | Z < -2 (shuffle) | 7 | 1.09 | 0.52 | 1.86 | 1.12 | t=2.877 df=6 | 0.03 |
| | | Z > -2 (non-shuffle) | 6 | 1.22 | 0.70 | 1.74 | 0.81 | t=3.582 df=5 | 0.02 |

| Swim | | Group (Z Swing Time) | N | Saline Average | SD | CNO Average | SD | Paired, two-tailed t test t, df | (alpha=0.05) p |
|---|---|---|---|---|---|---|---|---|---|
| **Duration** | burst | Z < -2 (shuffle) | 7 | 0.06 | 0.013 | 0.06 | 0.010 | t=0.2822 df=6 | 0.79 |
| | | Z > -2 (non-shuffle) | 6 | 0.07 | 0.010 | 0.05 | 0.006 | t=3.408 df=5 | 0.02 |
| | interburst | Z < -2 (shuffle) | 7 | 0.11 | 0.011 | 0.11 | 0.013 | t=0.8298 df=6 | 0.44 |
| | | Z > -2 (non-shuffle) | 6 | 0.11 | 0.015 | 0.11 | 0.022 | t=0.6514 df=5 | 0.54 |

| | | Group (Z Swing Time) | N | Saline Median | Lower 95% CI | Upper 95% CI | CNO Median | Lower 95% CI | Upper 95% CI | Wilcoxon two-tailed paired ranks test W | p |
|---|---|---|---|---|---|---|---|---|---|---|---|
| **Frequency** | (Hz) | Z < -2 (shuffle) | 7 | 5.8 | 5.3 | 6.4 | 6.2 | 5.8 | 6.6 | 14 | 0.30 |
| | | Z > -2 (non-shuffle) | 6 | 5.7 | 5.2 | 6.3 | 6.4 | 5.8 | 7.5 | 21 | 0.03 |

### Gastrocnemius EMG in VGluT2 sub-cohorts

| Walk | | Group (Z Swing Time) | N | Saline Median | Lower 95% CI | Upper 95% CI | CNO Median | Lower 95% CI | Upper 95% CI | Wilcoxon two-tailed paired ranks test W | p |
|---|---|---|---|---|---|---|---|---|---|---|---|
| **Duration** | burst (sec) | Z < -2 (shuffle) | 5 | 0.30 | 0.20 | 0.45 | 0.43 | 0.28 | 0.55 | 11 | 0.19 |
| | | Z > -2 (non-shuffle) | 5 | 0.35 | 0.24 | 0.46 | 0.38 | 0.27 | 0.47 | 7 | 0.44 |
| | interburst (sec) | Z < -2 (shuffle) | 5 | 0.06 | 0.01 | 0.18 | 0.06 | 0.05 | 0.07 | -1 | >0.99 |
| | | Z > -2 (non-shuffle) | 5 | 0.06 | 0.03 | 0.10 | 0.05 | 0.03 | 0.08 | -9 | 0.31 |
| **Peak amplitude** | burst | Z < -2 (shuffle) | 5 | 0.19 | 0.14 | 0.33 | 0.23 | 0.13 | 0.45 | 13 | 0.13 |
| (Normalized) | | Z > -2 (non-shuffle) | 5 | 0.42 | 0.03 | 1.09 | 0.54 | -0.05 | 1.57 | 15 | 0.06 |
| | interburst | Z < -2 (shuffle) | 5 | 0.17 | 0.08 | 0.30 | 0.26 | 0.07 | 0.48 | 15 | 0.06 |
| | | Z > -2 (non-shuffle) | 5 | 0.41 | 0.04 | 1.11 | 0.46 | 0.09 | 1.04 | 5 | 0.63 |
| **Integral** | burst | Z < -2 (shuffle) | 5 | 1.03 | 0.67 | 1.71 | 1.54 | 0.88 | 2.13 | 11 | 0.19 |
| (Normalized) | | Z > -2 (non-shuffle) | 5 | 1.61 | 1.20 | 2.76 | 2.46 | 1.82 | 3.30 | 15 | 0.06 |
| | interburst | Z < -2 (shuffle) | 5 | 0.11 | -0.22 | 0.78 | 0.16 | 0.06 | 0.27 | 2 | 0.88 |
| | | Z > -2 (non-shuffle) | 5 | 0.33 | 0.11 | 0.58 | 0.39 | 0.11 | 0.70 | 8 | 0.25 |

| Swim | | Group (Z Swing Time) | N | Saline Median | Lower 95% CI | Upper 95% CI | CNO Median | Lower 95% CI | Upper 95% CI | W | p |
|---|---|---|---|---|---|---|---|---|---|---|---|
| **Duration** | burst | Z < -2 (shuffle) | 5 | 0.05 | 0.04 | 0.06 | 0.04 | 0.03 | 0.08 | 3 | 0.81 |
| | | Z > -2 (non-shuffle) | 5 | 0.05 | 0.01 | 0.11 | 0.05 | 0.03 | 0.07 | -5 | 0.63 |
| | interburst | Z < -2 (shuffle) | 5 | 0.12 | 0.09 | 0.14 | 0.11 | 0.09 | 0.14 | 1 | >0.99 |
| | | Z > -2 (non-shuffle) | 5 | 0.08 | 0.02 | 0.15 | 0.09 | 0.02 | 0.13 | -5 | 0.63 |

**Supplementary Table 11**
with Supplementary Figure 11 and results

### Does unilateral or bilateral activation drive results in VGluT2 sub-cohorts?

| Group Z swing time>-2 bilateral | | Sum of squares F test (alpha 0.001) F (DFn, DFd) | p |
|---|---|---|---|
| N = 6 | Stride velocity (cm/s) | - | - |
| | Stride length (mm) | 3.04 (2, 529) | 0.05 |
| n Strides: | Swing time (s) | 3.27 (2, 466) | 0.04 |
| saline n = 190 | Stance time (s) | 1.55 (4, 463) | 0.19 |
| CNO n = 273 | Cadence (Hz) | 0.09 (2, 455) | 0.91 |
| | Swing speed (mm) | 24.14 (2, 512) | <0.0001 |

| Group Z>-2 unilateral | | Sum of squares F test (alpha 0.001) F (DFn, DFd) | p |
|---|---|---|---|
| N= 8 | Stride velocity (cm/s) | - | - |
| | Stride length (mm) | 11.21 (2, 539) | <0.0001 |
| n Strides: | Swing time (s) | 4.26 (2, 539) | 0.01 |
| saline: n = 277 | Stance time (s) | 5.48 (4, 593) | 0.0002 |
| CNO: n = 454 | Cadence (Hz) | 7.69 (2, 509) | 0.0005 |
| | Swing speed (mm) | 21.85 (2, 645) | <0.0001 |

| Temporal coupling<br>Swing Time | | saline<br>r | CNO<br>r | Watson (alpha 0.05) |
|---|---|---|---|---|
| Group Z swing time >-2 bilateral | Back left-Back right | 0.64 | 0.59 | 0.78 |
| Group Z swing time >-2 unilateral | Back left-Back right | 0.75 | 0.63 | 0.0009 |

Supplementary Figure 11 **Subanalysis: Asymmetry** — **Gait metrics in ipsi- or contralateral hindlimbs: comparison between saline versus CNO (all Z swing time <-3)**

| Ipsilateral hindlimb | Saline vs CNO | Sum of squares F test (alpha 0.001) F (DFn, DFd) | p |
|---|---|---|---|
| N = 15 | Stride velocity (cm/s) | - | - |
| | Stride length (mm) | 34.68 (2, 448) | <0.0001 |
| Strides | Swing time (s) | 16.9 (2, 448) | <0.0001 |
| saline n = 249 | Stance time (s) | 8.861 (4, 419) | <0.0001 |
| CNO n = 313 | Cadence (Hz) | 16.57 (2, 436) | <0.0001 |
| | Swing speed (mm) | 55.27 (2, 497) | <0.0001 |

| Contralateral hindlimb | Saline vs CNO | Sum of squares F test (alpha 0.001) F (DFn, DFd) | p |
|---|---|---|---|
| N = 15 | Stride velocity (cm/s) | - | - |
| | Stride length (mm) | 41.22 (2, 453) | <0.0001 |
| Strides | Swing time (s) | 20.91 (2, 453) | <0.0001 |
| sal n = 249 | Stance time (s) | - | - |
| CNO n = 329 | Cadence (Hz) | 20.43 (2, 449) | <0.0001 |
| | Swing speed (mm) | 81.54 (2, 480) | <0.0001 |

Supplementary Figure 11 **Subanalysis: Asymmetry** — **Gait metrics in saline or CNO condition: comparison between ipsi- versus contralateral hindlimbs (all Z swing time <-3)**

| Saline | ipsi- vs contralateral hindlimb | Sum of squares F test (alpha 0.001) F (DFn, DFd) | p |
|---|---|---|---|
| N = 15 | Stride velocity (cm/s) | - | - |
| | Stride length (mm) | 0.09698 (2, 387) | 0.91 |
| Strides | Swing time (s) | 0.3077 (2, 387) | 0.74 |
| ipsilateral n = 249 | Stance time (s) | - | - |
| contralateral n = 249 | Cadence (Hz) | 0.03389 (2, 326) | 0.97 |
| | Swing speed (mm) | 3.135 (2, 435) | 0.04 |

| CNO | ipsi- vs contralateral hindlimb | Sum of squares F test (alpha 0.001) F (DFn, DFd) | p |
|---|---|---|---|
| N = 15 | Stride velocity (cm/s) | - | - |
| | Stride length (mm) | 0.9296 (2, 514) | 0.40 |
| Strides | Swing time (s) | 10.21 (2, 514) | <0.0001 |
| ipsilateral n = 313 | Stance time (s) | 0.5319 (4, 508) | 0.71 |
| contralateral n = 329 | Cadence (Hz) | 3.04 (2, 559) | 0.05 |
| | Swing speed (mm) | 7.704 (2, 542) | 0.0005 |

**Subanalysis: Motor tests in bilateral and unilateral cases (irrespective of Z swingtime score)**

| | | | Saline | | | CNO | | | Wilcoxon two-tailed paired ranks test | (alpha=0.05) |
|---|---|---|---|---|---|---|---|---|---|---|
| **Rotarod** | | N | Median | Lower 95% CI | Upper 95% CI | Median | Lower 95% CI | Upper 95% CI | W | p |
| Latency to fall (sec) | Unilateral | 20 | 115 | 102 | 127 | 116 | 100 | 129 | 9 | 0.88 |
| | Bilateral | 13 | 120 | 90 | 133 | 120 | 83 | 137 | -1 | >0.9999 |
| **Beam** | | N | Median | Lower 95% CI | Upper 95% CI | Median | Lower 95% CI | Upper 95% CI | W | p |
| # Slips and misses | Unilateral | 20 | 10 | 7 | 20 | 8 | 6 | 22 | 23 | 0.57 |
| | Bilateral | 13 | 12 | 7 | 23 | 9 | 3 | 30 | -20 | 0.46 |
| **Ladder** | | N | Median | Lower 95% CI | Upper 95% CI | Median | Lower 95% CI | Upper 95% CI | W | p |
| # Slips and misses | Unilateral | 20 | 1.5 | 1.02 | 2.48 | 2.00 | 2.03 | 4.17 | 79 | 0.04 |
| | Bilateral | 13 | 2 | 1.05 | 2.79 | 1.00 | 0.25 | 1.44 | -34 | 0.04 |

**Supplementary Table 12**
with Figure 5
and Supplementary Figure 6

### Gait metrics in the SerT cohort

#### Footfall metrics

| | high Z Stride Velocity (faster) | | Sum of squares F test (alpha 0.001) F (DFn, DFd) | p | | Saline Average | Saline SD | CNO Average | CNO SD | paired, 2 tailed t-test (alpha 0.05) t(df) | p |
|---|---|---|---|---|---|---|---|---|---|---|---|
| Figure 5 | N= 6 | Stride velocity (cm/s) | - | - | Stride velocity (cm/s) | 11.37 | 2.24 | 15.61 | 1.84 | t=8.82 df=5 | 0.0003 |
| | | Stride length (mm) | 0.00154 (2, 302) | 1.00 | Stride length (mm) | 41.26 | 4.35 | 49.18 | 4.42 | t=9.81 df=5 | 0.0002 |
| | n Strides: | Swing time (s) | 2.206 (2, 302) | 0.11 | Swing time (s) | 0.12 | 0.02 | 0.12 | 0.01 | t=0.098 df=5 | 0.9255 |
| | saline: n = 236 | Stance time (s) | 0.7058 (4, 298) | 0.59 | Stance time (s) | 0.31 | 0.05 | 0.23 | 0.04 | t=6.17 df=5 | 0.0016 |
| | CNO: n = 223 | Cadence (Hz) | 0.001025 (2, 226) | 1.00 | Cadence (Hz) | 2.72 | 0.35 | 3.10 | 0.17 | t=3.03 df=5 | 0.03 |
| | | Swing speed (mm) | 5.577 (2, 427) | 0.004 | Swing speed (cm/s) | 36.63 | 6.78 | 42.15 | 3.84 | t=4.43 df=5 | 0.01 |

| | intermediate Z Stride Velocity | | Sum of squares F test (alpha 0.001) F (DFn, DFd) | p | | Saline Average | Saline SD | CNO Average | CNO SD | paired, 2 tailed t-test (alpha 0.05) t(df) | p |
|---|---|---|---|---|---|---|---|---|---|---|---|
| | N= 6 | Stride velocity (cm/s) | - | - | Stride velocity (cm/s) | 11.97 | 2.14 | 12.71 | 2.31 | t=1.5 df=5 | 0.19 |
| | | Stride length (mm) | 1.02 (2, 371) | 0.36 | Stride length (mm) | 42.29 | 5.83 | 42.93 | 6.66 | t=0.36 df=5 | 0.73 |
| | n Strides: | Swing time (s) | 1.01 (2, 321) | 0.36 | Swing time (s) | 0.12 | 0.02 | 0.12 | 0.02 | t=0.31 df=5 | 0.77 |
| | saline: n = 229 | Stance time (s) | 1.73 (4, 367) | 0.14 | Stance time (s) | 0.27 | 0.05 | 0.24 | 0.03 | t=2.65 df=5 | 0.05 |
| | CNO: n = 251 | Cadence (Hz) | 1.61 (2, 271) | 0.20 | Cadence (Hz) | 2.83 | 0.34 | 3.02 | 0.23 | t=2.37 df=5 | 0.06 |
| | | Swing speed (mm) | 3.50 (2, 461) | 0.03 | Swing speed (cm/s) | 36.48 | 4.43 | 36.16 | 3.92 | t=0.34 df=5 | 0.75 |

| Supplementary Figure 6 | low Z Stride Velocity (slower) | | Sum of squares F test (alpha 0.001) F (DFn, DFd) | p | | Saline Average | Saline SD | CNO Average | CNO SD | paired, 2 tailed t-test (alpha 0.05) t(df) | p |
|---|---|---|---|---|---|---|---|---|---|---|---|
| | N= 6 | Stride velocity (cm/s) | - | - | Stride velocity (cm/s) | 12.79 | 4.27 | 8.63 | 1.67 | t=2.93 df=5 | 0.03 |
| | | Stride length (mm) | 2.94 (2, 369) | 0.05 | Stride length (mm) | 43.46 | 5.61 | 36.32 | 4.01 | t=3.95 df=5 | 0.01 |
| | n Strides: | Swing time (s) | 1.972 (2, 369) | 0.14 | Swing time (s) | 0.11 | 0.02 | 0.10 | 0.02 | t=2.33 df=5 | 0.07 |
| | saline: n = 229 | Stance time (s) | 2.264 (4, 370) | 0.06 | Stance time (s) | 0.29 | 0.11 | 0.39 | 0.04 | t=2.89 df=5 | 0.03 |
| | CNO: n = 241 | Cadence (Hz) | 2.55 (2, 319) | 0.08 | Cadence (Hz) | 2.92 | 0.59 | 2.35 | 0.18 | t=2.73 df=5 | 0.04 |
| | | Swing speed (mm) | 8.995 (2, 436) | 0.0001 | Swing speed (cm/s) | 39.72 | 5.08 | 35.46 | 3.82 | t=2.4 df=5 | 0.06 |

| | WT control | | Sum of squares F test (alpha 0.001) F (DFn, DFd) | p | | Saline Average | Saline SD | CNO Average | CNO SD | paired, 2 tailed t-test (alpha 0.05) t(df) | p |
|---|---|---|---|---|---|---|---|---|---|---|---|
| | N= 15 | Stride velocity (cm/s) | - | - | Stride velocity (cm/s) | 12.93 | 2.20 | 12.92 | 2.62 | t=0.003879 df=14 | 1.00 |
| | | Stride length (mm) | 0.45 (2, 829) | 0.64 | Stride length (mm) | 43.94 | 5.93 | 42.64 | 5.66 | t=0.7559 df=14 | 0.46 |
| | n Strides: | Swing time (s) | 0.19 (2, 829) | 0.82 | Swing time (s) | 0.12 | 0.02 | 0.11 | 0.01 | t=1.149 df=14 | 0.27 |
| | saline: n = 567 | Stance time (s) | 1.9 (4, 802) | 0.11 | Stance time (s) | 0.26 | 0.05 | 0.25 | 0.04 | t=0.2607 df=14 | 0.80 |
| | CNO: n = 604 | Cadence (Hz) | 0.38 (2, 622) | 0.69 | Cadence (Hz) | 2.97 | 0.38 | 3.04 | 0.30 | t=0.6387 df=14 | 0.53 |
| | | Swing speed (mm) | 3.5 (2, 1114) | 0.03 | Swing speed (cm/s) | 38.48 | 4.66 | 38.40 | 3.39 | t=0.0681 df=14 | 0.95 |

#### Temporal coupling

| | Stride Velocity | Comparison | saline r | CNO r | Watson (alpha 0.05) |
|---|---|---|---|---|---|
| Figure 5 | High Z (faster) | Back left-Back right | 0.67 | 0.72 | 0.30 |
| | Intermediate Z | Back left-Back right | 0.76 | 0.76 | 0.65 |
| Supplementary Figure 6 | Low Z (slower) | Back left-Back right | 0.65 | 0.56 | 0.43 |
| | WT control | Back left-Back right | 0.64 | 0.75 | 0.02 |

#### Postural metrics

| | | N | Mean | SD | Mean | SD | paired, 2 tailed t-test (alpha 0.05) t, df | p |
|---|---|---|---|---|---|---|---|---|
| Figure 5 | **High Z velocity (faster)** | | | | | | | |
| | **Height start swing** | | | | | | | |
| | Iliac crest to floor (mm) | 6 | 27.3 | 2.6 | 28.5 | 4.0 | t=1.21 df=5 | 0.28 |
| | Tail base to floor (mm) | 6 | 13.8 | 1.7 | 15.8 | 3.9 | t=1.774 df=5 | 0.14 |
| | **Height end swing** | | | | | | | |
| | Iliac crest to floor (mm) | 6 | 28.2 | 3.0 | 29.7 | 3.6 | t=1.487 df=5 | 0.20 |
| | Tail base to floor (mm) | 6 | 13.3 | 1.9 | 17.1 | 4.2 | t=2.464 df=5 | 0.06 |

| | | N | Mean | SD | Mean | SD | t, df | p |
|---|---|---|---|---|---|---|---|---|
| Supplementary Figure 6 | **Low Z velocity (slower)** | | | | | | | |
| | **Height start swing** | | | | | | | |
| | Iliac crest to floor (mm) | 6 | 27.3 | 1.6 | 26.7 | 2.7 | t=0.4799 df=5 | 0.65 |
| | Tail base to floor (mm) | 6 | 14.8 | 3.0 | 14.5 | 2.2 | t=0.1655 df=5 | 0.88 |
| | **Height end swing** | | | | | | | |
| | Iliac crest to floor (mm) | 6 | 28.5 | 2.3 | 27.4 | 2.6 | t=0.8578 df=5 | 0.43 |
| | Tail base to floor (mm) | 6 | 15.8 | 2.4 | 14.5 | 2.7 | t=0.7678 df=5 | 0.48 |

**Supplementary Table 13** with Supplementary Figure 7

### Anterior Tibial EMG in the SerT cohort

| Walk | | Group (Z Stride Velocity) | N | Saline Median | Lower 95% CI | Upper 95% CI | CNO Median | Lower 95% CI | Upper 95% CI | Wilcoxon matched-pa W | (alpha=0.05) p |
|---|---|---|---|---|---|---|---|---|---|---|---|
| **Duration** | burst (sec) | Z>0 (faster) | 5 | 0.07 | 0.04 | 0.08 | 0.07 | 0.05 | 0.09 | 9 | 0.31 |
| | | Z<0 (slower) | 3 | 0.09 | 0.06 | 0.14 | 0.12 | 0.09 | 0.14 | 4 | 0.50 |
| | | WT | 7 | 0.07 | 0.06 | 0.10 | 0.08 | 0.06 | 0.10 | 6 | 0.69 |
| | interburst (sec) | Z>0 (faster) | 5 | 0.30 | 0.26 | 0.32 | 0.35 | 0.21 | 0.47 | 7 | 0.44 |
| | | Z<0 (slower) | 3 | 0.23 | 0.10 | 0.42 | 0.25 | 0.04 | 0.50 | 0 | >0.9999 |
| | | WT | 7 | 0.28 | 0.24 | 0.32 | 0.28 | 0.21 | 0.43 | 14 | 0.30 |
| **Peak amplitude** (Normalized) | burst | Z>0 (faster) | 5 | 0.29 | 0.10 | 0.37 | 0.30 | 0.12 | 0.42 | 15 | 0.06 |
| | | Z<0 (slower) | 3 | 0.44 | 0.06 | 0.69 | 0.46 | 0.10 | 0.70 | 6 | 0.25 |
| | | WT | 7 | 0.34 | 0.23 | 0.42 | 0.37 | 0.25 | 0.42 | 6 | 0.69 |
| | interburst | Z>0 (faster) | 5 | 0.27 | 0.10 | 0.38 | 0.30 | 0.10 | 0.48 | 15 | 0.06 |
| | | Z<0 (slower) | 3 | 0.34 | -0.04 | 0.87 | 0.38 | 0.11 | 0.70 | 0 | >0.9999 |
| | | WT | 7 | 0.40 | 0.33 | 0.48 | 0.40 | 0.31 | 0.57 | 10 | 0.47 |
| **Integral** (Normalized) | burst | Z>0 (faster) | 5 | 0.27 | 0.17 | 0.38 | 0.35 | 0.24 | 0.45 | 15 | 0.06 |
| | | Z<0 (slower) | 3 | 0.69 | 0.06 | 1.10 | 0.80 | 0.19 | 1.20 | 6 | 0.25 |
| | | WT | 7 | 0.45 | 0.28 | 0.59 | 0.46 | 0.34 | 0.55 | 4 | 0.81 |
| | interburst | Z>0 (faster) | 5 | 0.42 | 0.24 | 0.84 | 0.69 | 0.34 | 0.98 | 9 | 0.31 |
| | | Z<0 (slower) | 3 | 0.85 | 0.49 | 1.25 | 0.90 | 0.11 | 1.59 | 0 | >0.9999 |
| | | WT | 7 | 1.15 | 0.90 | 1.24 | 1.17 | 0.77 | 1.93 | 18 | 0.16 |

| Swim | | Group (Z Stride Velocity) | N | Saline Median | Lower 95% CI | Upper 95% CI | CNO Median | Lower 95% CI | Upper 95% CI | Wilcoxon matched-pa W | (alpha=0.05) p |
|---|---|---|---|---|---|---|---|---|---|---|---|
| **Duration** | burst (sec) | Z>0 (faster) | 5 | 0.04 | 0.04 | 0.06 | 0.05 | 0.03 | 0.06 | 1 | >0.9999 |
| | | Z<0 (slower) | 3 | 0.07 | 0.05 | 0.08 | 0.06 | 0.00 | 0.14 | 0 | >0.9999 |
| | | WT | 7 | 0.06 | 0.04 | 0.08 | 0.04 | 0.03 | 0.08 | -8 | 0.58 |
| | interburst (sec) | Z>0 (faster) | 5 | 0.11 | 0.09 | 0.18 | 0.11 | 0.08 | 0.14 | -13 | 0.13 |
| | | Z<0 (slower) | 3 | 0.11 | 0.06 | 0.15 | 0.09 | 0.04 | 0.13 | -4 | 0.50 |
| | | WT | 7 | 0.10 | 0.07 | 0.13 | 0.10 | 0.09 | 0.12 | 0 | >0.9999 |

| | | Group (Z Stride Velocity) | N | Saline Median | Lower 95% CI | Upper 95% CI | CNO Median | Lower 95% CI | Upper 95% CI | Wilcoxon two-tailed paired ranks test W | p |
|---|---|---|---|---|---|---|---|---|---|---|---|
| **Frequency** | (Hz) | Z>0 (faster) | 5 | 6.1 | 5.1 | 6.9 | 6.5 | 5.9 | 6.9 | 15 | 0.06 |
| | | Z<0 (slower) | 3 | 6.3 | 5.3 | 7.1 | 6.9 | 5.5 | 7.8 | 6 | 0.25 |
| | | WT | 7 | 6.4 | 6.1 | 6.8 | 6.6 | 6.0 | 7.2 | 8 | 0.58 |

### Gastrocnemius EMG in the SerT cohort

| Walk | | Group (Z Swing Time) | N | Saline Median | Lower 95% CI | Upper 95% CI | CNO Median | Lower 95% CI | Upper 95% CI | Wilcoxon two-tailed paired ranks test W | p |
|---|---|---|---|---|---|---|---|---|---|---|---|
| **Duration** | burst (sec) | Z>0 (faster) | 5 | 0.24 | 0.21 | 0.27 | 0.30 | 0.22 | 0.35 | 11 | 0.19 |
| | | Z<0 (slower) | 3 | 0.28 | 0.23 | 0.34 | 0.28 | 0.13 | 0.50 | 4 | 0.50 |
| | interburst (sec) | Z>0 (faster) | 5 | 0.09 | 0.07 | 0.14 | 0.08 | 0.04 | 0.20 | 7 | 0.44 |
| | | Z<0 (slower) | 3 | 0.06 | 0.0003 | 0.15 | 0.07 | 0.04 | 0.10 | -2 | 0.75 |
| **Peak amplitude** (Normalized) | burst | Z>0 (faster) | 5 | 0.21 | 0.09 | 0.49 | 0.23 | -0.18 | 1.09 | 5 | 0.63 |
| | | Z<0 (slower) | 3 | NA | | | NA | | | | |
| | interburst | Z>0 (faster) | 5 | 0.30 | 0.20 | 0.43 | 0.24 | -0.03 | 0.81 | -1 | >0.99 |
| | | Z<0 (slower) | 2 | NA | | | NA | | | | |
| **Integral** (Normalized) | burst | Z>0 (faster) | 5 | 0.93 | 0.72 | 1.21 | 1.26 | 0.83 | 1.74 | 13 | 0.13 |
| | | Z<0 (slower) | 3 | NA | | | NA | | | | |
| | interburst | Z>0 (faster) | 5 | 0.33 | 0.08 | 0.81 | 0.49 | 0.28 | 0.65 | 5 | 0.63 |
| | | Z<0 (slower) | 2 | NA | | | NA | | | | |

| Swim | | Group (Z Swing Time) | N | Saline Median | Lower 95% CI | Upper 95% CI | CNO Median | Lower 95% CI | Upper 95% CI | Wilcoxon two-tailed paired ranks test W | p |
|---|---|---|---|---|---|---|---|---|---|---|---|
| **Duration** | burst | Z>0 (faster) | 5 | 0.06 | 0.03 | 0.12 | 0.05 | 0.01 | 0.12 | -5 | 0.63 |
| | | Z<0 (slower) | 3 | 0.09 | 0.04 | 0.12 | 0.10 | 0.03 | 0.15 | 4 | 0.50 |
| | interburst | Z>0 (faster) | 5 | 0.10 | 0.06 | 0.13 | 0.11 | 0.06 | 0.13 | 3 | 0.81 |
| | | Z<0 (slower) | 3 | 0.07 | 0.02 | 0.14 | 0.05 | -0.02 | 0.15 | -6 | 0.25 |

**Supplementary Table 14** **Motor tests in the SerT cohort**

| | | | Saline | | | CNO | | | Wilcoxon matched-| (alpha=0.05) |
|---|---|---|---|---|---|---|---|---|---|---|
| **Rotarod** | **Group (Z Stride Velocity)** | **N** | **Median** | **Lower 95% CI** | **Upper 95% CI** | **Median** | **Lower 95% CI** | **Upper 95% CI** | **W** | **p** |
| Latency to fall (sec) | High Z stride velocity (faster) | 6 | 107.3 | 75.44 | 132 | 105.3 | 87.87 | 132 | 9 | 0.44 |
| | Intermediate Z stride velocity | 6 | 91.33 | 55.31 | 123.9 | 100.5 | 68.66 | 136.1 | 13 | 0.22 |
| | Low Z stride velocity (slower) | 6 | 89.5 | 53.4 | 111.8 | 96 | 60.25 | 137 | 15 | 0.16 |
| | WT control | 15 | 113.7 | 96.92 | 127.5 | 99.33 | 92.02 | 113.6 | -66 | 0.06 |
| **Beam** | **Group (Z Stride Velocity)** | **N** | **Median** | **Lower 95% CI** | **Upper 95% CI** | **Median** | **Lower 95% CI** | **Upper 95% CI** | **W** | **p** |
| Slips and misses | High Z stride velocity (faster) | 6 | 5 | 0.9911 | 8.342 | 3 | 0.924 | 5.409 | -9 | 0.47 |
| | Intermediate Z stride velocity | 6 | 7.5 | 3.127 | 14.21 | 5 | 1.215 | 10.45 | -8 | 0.47 |
| | Low Z stride velocity (slower) | 6 | 9.5 | 1.485 | 25.18 | 10.5 | 2.599 | 20.73 | -4 | 0.72 |
| | WT control | 15 | 5 | 3.684 | 7.916 | 4 | 3.347 | 6.253 | -19 | 0.61 |
| **Ladder** | **Group (Z Stride Velocity)** | **N** | **Median** | **Lower 95% CI** | **Upper 95% CI** | **Median** | **Lower 95% CI** | **Upper 95% CI** | **W** | **p** |
| Slips and misses | High Z stride velocity (faster) | 6 | 0.5 | -0.5146 | 2.848 | 0.5 | -0.1902 | 1.524 | -3 | 0.81 |
| | Intermediate Z stride velocity | 6 | 1.5 | 0.2329 | 3.1 | 0.5 | -0.6955 | 4.029 | -2 | >0.9999 |
| | Low Z stride velocity (slower) | 6 | 0.5 | -0.1902 | 1.524 | 1.5 | -0.8266 | 5.493 | 7 | 0.38 |
| | WT control | 15 | 0 | 0.3992 | 2.934 | 1 | 0.2659 | 1.067 | -27 | 0.14 |

**Supplementary Table 15** | **Quantification of synaptophysin labeled boutons onto spinal profiles**

**With Supplementary Figure 15**

Fig. 15 c **Boutons in apposition to ChAT-IR motoneuron soma, dendrite or C bouton, per genotype**

| Ordinary One Way Anova | |
|---|---|
| **F (DFn, DFd)** | **P value** |
| F (5, 30) = 14.69 | <0.0001 |

| Sidak's multiple comparisons test | Mean Diff. | 95.00% CI of diff. | Summary | Adjusted P Value |
|---|---|---|---|---|
| VGluT2 Soma vs. VGat Soma | -0.5 | -26.80 to 25.80 | ns | >0.9999 |
| VGluT2 Dendrite vs. VGaT Dendrite | 0 | -26.30 to 26.30 | ns | >0.9999 |
| VGluT2 C-bouton vs. VGaT C-bouton | 28.5 | 2.195 to 54.80 | * | 0.03 |
| VGluT2 Soma vs. VGluT2 Dendrite | -46.5 | -72.80 to -20.20 | **** | <0.0001 |
| VGluT2 Soma vs. VGluT2 C-bouton | -22.17 | -48.47 to 4.138 | ns | 0.15 |
| VGluT2 Dendrite vs. VGluT2 C-bouton | 24.33 | -1.971 to 50.64 | ns | 0.09 |
| VGat Soma vs. VGaT Dendrite | -46 | -72.30 to -19.70 | *** | 0.0001 |
| VGat Soma vs. VGaT C-bouton | 6.833 | -19.47 to 33.14 | ns | 0.995 |
| VGaT Dendrite vs. VGaT C-bouton | 52.83 | 26.53 to 79.14 | **** | <0.0001 |

Fig. 15 d **Boutons in lamina IX (standardized volume)**

| Ordinary One Way Anova | |
|---|---|
| **F (DFn, DFd)** | **P value** |
| F (3, 23) = 10.70 | 0.0001 |

| Tukey's multiple comparisons test | Mean Diff. | 95.00% CI of diff. | Summary | Adjusted P Value |
|---|---|---|---|---|
| VGluT2 rostral vs. VGluT2 intermediate | 25.33 | -10.26 to 60.93 | ns | 0.23 |
| VGluT2 rostral vs. VGaT rostral | 50 | 16.44 to 83.56 | ** | 0.002 |
| VGluT2 rostral vs. VGaT intermediate | 6.333 | -27.23 to 39.90 | ns | 0.95 |
| VGluT intermediate vs. VGaT rostral | 24.67 | -1.867 to 51.20 | ns | 0.07 |
| VGluT intermediate vs. VGaT intermediate | -19 | -45.53 to 7.533 | ns | 0.22 |
| VGaT rostral vs. VGaT intermediate | -43.67 | -67.40 to -19.93 | *** | 0.0002 |

Fig. 15 f **Boutons in area X (standardized volume)**

| Kruskal-Wallis test | P value |
|---|---|
| KW 9.482 | 0.0009 |

| Dunn's multiple comparisons test | Mean rank diff. | Summary | Adjusted P Value |
|---|---|---|---|
| VGluT2 rostral vs. VGluT2 intermediate | -1.667 | ns | >0.9999 |
| VGluT2 rostral vs. VGaT rostral | 6.5 | ns | 0.16 |
| VGluT2 rostral vs. VGaT intermediate | 3.833 | ns | >0.9999 |
| VGluT intermediate vs. VGaT rostral | 8.167 | * | 0.03 |
| VGluT intermediate vs. VGaT intermediate | 5.5 | ns | 0.37 |
| VGaT rostral vs. VGaT intermediate | -2.667 | ns | >0.9999 |

Fig. 15 e **Boutons in apposition to CTb labeled TA, G, Glut motoneuron soma or proximal dendrite (standardized volume)**

| Kruskal-Wallis test | P value |
|---|---|
| KW 14.30 | 0.003 |

| Dunn's multiple comparisons test | Mean rank diff. | Summary | Adjusted P Value |
|---|---|---|---|
| VGluT2 rostral vs. VGluT2 intermediate | 9.35 | ns | 0.47 |
| VGluT2 rostral vs. VGaT rostral | 6.156 | ns | 0.98 |
| VGluT2 rostral vs. VGaT intermediate | -5.511 | ns | >0.9999 |
| VGluT intermediate vs. VGaT rostral | -3.194 | ns | >0.9999 |
| VGluT intermediate vs. VGaT intermediate | -14.86 | * | 0.01 |
| VGaT rostral vs. VGaT intermediate | -11.67 | * | 0.01 |

Fig. 15 g **Boutons in area X in apposition to ChAT-IR profiles (standardized volume)**

| Kruskal-Wallis test | P value |
|---|---|
| KW 7.410 | 0.03 |

| Dunn's multiple comparisons test | Mean rank diff. | Summary | Adjusted P Value |
|---|---|---|---|
| VGluT2 rostral vs. VGluT2 intermediate | 0 | ns | >0.9999 |
| VGluT2 rostral vs. VGaT rostral | 4.333 | ns | 0.85 |
| VGluT2 rostral vs. VGaT intermediate | -3.667 | ns | >0.9999 |
| VGluT intermediate vs. VGaT rostral | 4.333 | ns | 0.85 |
| VGluT intermediate vs. VGaT intermediate | -3.667 | ns | >0.9999 |
| VGaT rostral vs. VGaT intermediate | -8 | * | 0.04 |

**Supplementary Table 16** | **Bouton densities in the lumbar gray matter of VGaT mice with high versus low Z swing time**

**With Figure 8**

**High Z (slomo; 2.6-5.7; N=5) versus Low Z swing time scores (non-slomo); 0.2-1.9; N=5)**

| 2 way ANOVA | SS | n DF | MS | F (DFn, DFd) | P value |
|---|---|---|---|---|---|
| Z score | 1441 | 1 | 1441 | F (1, 48) = 6.227 | 0.02 |
| Region | 12199 | 5 | 2440 | F (5, 48) = 10.55 | <0.0001 |
| Interaction | 1807 | 5 | 361.4 | F (5, 48) = 1.562 | 0.19 |

| Sidak's multiple compariso | P Value, α 0.05 |
|---|---|
| lamina I-II | 1.00 |
| lamina III | 0.02 |
| lat. Lamina V | 0.54 |
| area X | 0.86 |
| lamina VIII | > 0.99 |
| lamina IX | > 0.99 |

**Correlation between swing Time Z score and bouton density/region**

N=10

| Pearson Correlation | r | ɔ5% confidence interva | R squared | P (two-tailed) |
|---|---|---|---|---|
| lamina I-II | 0.30 | -0.4069 to 0.7817 | 0.09 | 0.40 |
| lamina III | 0.65 | 0.02904 to 0.9071 | 0.42 | 0.04 |
| lat. Lamina V | 0.58 | -0.07162 to 0.8875 | 0.34 | 0.08 |
| area X | 0.53 | -0.1531 to 0.8686 | 0.28 | 0.12 |
| medial lamina VII | 0.12 | -0.5491 to 0.6986 | 0.02 | 0.73 |
| lamina IX | -0.33 | -0.7942 to 0.3789 | 0.11 | 0.35 |

**Supplementary Table 17** | **Sensory tests in sub-groups of the VGaT cohort**

**With Figure 8**

**Von Frey**

| Group | N | Source of Variation | % of total variation | F (DFn, DFd) | P value |
|---|---|---|---|---|---|
| Z swingtime >2 (slomo: N=9 out of 14 mice that underwent von Frey testing) | 9 | Filament | 31.35 | F (3, 24) = 18.19 | < 0.0001 |
| | | Treatment (Saline/CNO | 15.31 | F (1, 8) = 8.490 | 0.02 |
| | | Interaction | 6.719 | F (3, 24) = 4.344 | 0.01 |

2 way RM ANOVA (above)

Sidak's multiple comparisons test

| Filament # | Mean Diff. | 95% CI of diff. | Summary |
|---|---|---|---|
| 0.4 | 0.1111 | -1.480 to 1.703 | ns |
| 0.6 | 1.111 | -0.4804 to 2.703 | ns |
| 1 | 2.444 | 0.8530 to 4.036 | ** |
| 2 | 2.778 | 1.186 to 4.369 | *** |

2 way RM ANOVA

| Group | N | Source of Variation | % of total variation | F (DFn, DFd) | P value |
|---|---|---|---|---|---|
| WT control (all Z<2) | 8 | Filament | 61.32 | F (3, 21) = 45.09 | < 0.0001 |
| | | Treatment | 0 | F (1, 7) = 0.0 | > 0.9999 |
| | | Interaction | 0.7719 | F (3, 21) = 0.4468 | 0.72 |

Sidak's multiple comparisons test

| Filament # | Mean Diff. | 95% CI of diff. | Summary |
|---|---|---|---|
| 0.4 | 0 | -1.247 to 1.247 | ns |
| 0.6 | -0.375 | -1.622 to 0.8723 | ns |
| 1 | 0 | -1.247 to 1.247 | ns |
| 2 | 0.375 | -0.8723 to 1.622 | ns |

**Pin**

| Group | N | Saline Median | CNO Median | o tailed Wilcoxon Rank test Sum of signed ranks (W) | P value |
|---|---|---|---|---|---|
| Z>2 (slomo): 6 out of 10 mice with Pin testing | 6 | 0.225 | 0.2 | -6 | 0.63 |
| WT control (all Z<2) | 8 | 0.235 | 0.22 | -19 | 0.21 |

**Hotplate**

| Group | N | Saline Median | CNO Median | o tailed Wilcoxon Rank test Sum of signed ranks (W) | P value |
|---|---|---|---|---|---|
| Z>2 (slomo): 6 out of 10 mice with pin testing | 6 | 7.55 | 10.35 | 11 | 0.31 |
| WT control (all Z<2) | 8 | 8.55 | 8.45 | -2 | 0.95 |

**Supplementary Table 18** with Figure 9

**Optogenetic analysis of Phase Specificity of VGaT mRF sub-regions**

Figure 9, d-i

**Slow-motion hotspot (rostral venteral mRI**

| Timing | TA bursts n | TA burst Duration (s) Mean | SD | TA Integral Mean | SD | TA Peak Amplitude Mean | SD |
|---|---|---|---|---|---|---|---|
| Pulse >0.1s prior to TA burst | 18 | 0.95 | 0.21 | 1.12 | 0.48 | 1.08 | 0.27 |
| Pulse <0.1s prior to TA burst | 30 | 0.96 | 0.20 | 1.21 | 0.43 | 1.15 | 0.31 |
| Pulse during TA burst | 42 | 1.09 | 0.30 | 1.34 | 0.41 | 1.14 | 0.22 |
| Pulse after TA burst | 62 | 0.85 | 0.20 | 1.06 | 0.26 | 1.12 | 0.28 |

**Two-way ANOVA**

| Source of Variation | % of total variation | F (DFn, DFd) | P value |
|---|---|---|---|
| Interaction | 1.46 | F (6, 444) = 1.234 | 0.29 |
| Timing | 3.71 | F (3, 444) = 6.264 | 0.0004 |
| Metric | 7.16 | F (2, 444) = 18.12 | <0.0001 |

**Duration**

| Tukey's multiple comparisons test | Mean Diff. | 95.00% CI of diff. | Adjusted P Value |
|---|---|---|---|
| Pre far vs. Pre close | -0.0184 | -0.246 to 0.2092 | 1.00 |
| Pre far vs. During | -0.1412 | -0.3562 to 0.07382 | 0.33 |
| Pre far vs. Post | 0.0922 | -0.1121 to 0.2965 | 0.65 |
| Pre close vs. During | -0.1228 | -0.3052 to 0.05965 | 0.31 |
| Pre close vs. Post | 0.1106 | -0.05914 to 0.2803 | 0.34 |
| During vs. Post | 0.2334 | 0.08087 to 0.3859 | **0.001** |

**Integral**

| Tukey's multiple comp | Mean Diff. | 95.00% CI of diff. | Adjusted P Value |
|---|---|---|---|
| Pre far vs. Pre close | -0.085 | -0.3126 to 0.1426 | 0.77 |
| Pre far vs. During | -0.215 | -0.43 to 1.72e-005 | 0.05 |
| Pre far vs. Post | 0.065 | -0.1393 to 0.2693 | 0.84 |
| Pre close vs. During | -0.13 | -0.3124 to 0.05245 | 0.26 |
| Pre close vs. Post | 0.15 | -0.01974 to 0.3197 | 0.10 |
| During vs. Post | 0.28 | 0.1275 to 0.4325 | <0.0001 |

**Amplitude**

| Tukey's multiple comparisoı | Mean Diff. | 95.00% CI of diff. | Adjusted P Value |
|---|---|---|---|
| Pre far vs. Pre close | -0.063 | -0.2906 to 0.1646 | 0.89 |
| Pre far vs. During | -0.056 | -0.271 to 0.159 | 0.91 |
| Pre far vs. Post | -0.038 | -0.2423 to 0.1663 | 0.96 |
| Pre close vs. During | 0.007 | -0.1754 to 0.1894 | 1.00 |
| Pre close vs. Post | 0.025 | -0.1447 to 0.1947 | 0.98 |
| During vs. Post | 0.018 | -0.1345 to 0.1705 | 0.99 |

Figure 9, d-ii

**Caudal ventral mRF**

| Timing | TA bursts n | TA burst Duration Mean | SD | TA Integral Mean | SD | TA Peak Amplitude Mean | SD |
|---|---|---|---|---|---|---|---|
| Pulse >0.1s prior to TA burst | 29 | 1.07 | 0.27 | 1.23 | 0.37 | 1.18 | 0.29 |
| Pulse <0.1s prior to TA burst | 67 | 1.08 | 0.26 | 1.37 | 0.40 | 1.32 | 0.38 |
| Pulse during TA burst | 74 | 0.98 | 0.27 | 1.47 | 0.40 | 1.53 | 0.41 |
| Pulse after TA burst | 55 | 0.96 | 0.25 | 1.19 | 0.33 | 1.27 | 0.28 |

**Two-way ANOVA**

| Source of Variation | % of total variation | F (DFn, DFd) | P value |
|---|---|---|---|
| Interaction | 3.06 | F (6, 663) = 4.173 | 0.0004 |
| Timing | 3.61 | F (3, 663) = 9.828 | <0.0001 |
| Metric | 12.20 | F (2, 663) = 49.84 | <0.0001 |

**Duration**

| Tukey's multiple comparisons test | Mean Diff. | 95.00% CI of diff. | Adjusted P Value |
|---|---|---|---|
| Pre far vs. Pre close | -0.007 | -0.2013 to 0.1873 | 1.00 |
| Pre far vs. During | 0.0874 | -0.1041 to 0.2789 | 0.64 |
| Pre far vs. Post | 0.1077 | -0.09289 to 0.3083 | 0.51 |
| Pre close vs. During | 0.0944 | -0.053 to 0.2418 | 0.35 |
| Pre close vs. Post | 0.1147 | -0.04434 to 0.2737 | 0.25 |
| During vs. Post | 0.0203 | -0.1353 to 0.1759 | 0.99 |

**Integral**

| Tukey's multiple comp | Mean Diff. | 95.00% CI of diff. | Adjusted P Value |
|---|---|---|---|
| Pre far vs. Pre close | -0.143 | -0.3373 to 0.05129 | 0.23 |
| Pre far vs. During | -0.238 | -0.4295 to -0.04651 | 0.01 |
| Pre far vs. Post | 0.036 | -0.1646 to 0.2366 | 0.97 |
| Pre close vs. During | -0.095 | -0.2424 to 0.0524 | 0.35 |
| Pre close vs. Post | 0.179 | 0.01996 to 0.338 | 0.02 |
| During vs. Post | 0.274 | 0.1184 to 0.4296 | <0.0001 |

**Amplitude**

| Tukey's multiple comparisoı | Mean Diff. | 95.00% CI of diff. | Adjusted P Value |
|---|---|---|---|
| Pre far vs. Pre close | -0.145 | -0.3393 to 0.04929 | 0.22 |
| Pre far vs. During | -0.355 | -0.5465 to -0.1635 | <0.0001 |
| Pre far vs. Post | -0.089 | -0.2896 to 0.1116 | 0.66 |
| Pre close vs. During | -0.21 | -0.3574 to -0.0626 | 0.002 |
| Pre close vs. Post | 0.056 | -0.103 to 0.215 | 0.80 |
| During vs. Post | 0.266 | 0.1104 to 0.4216 | <0.0001 |

Figure 9, d-iii

**Avoiding ventral mRF**

| Timing | TA bursts n | TA burst Duration Mean | SD | TA Integral Mean | SD | TA Peak Amplitude Mean | SD |
|---|---|---|---|---|---|---|---|
| Pulse >0.1s prior to TA burst | 28 | 1.04 | 0.27 | 1.18 | 0.37 | 1.18 | 0.39 |
| Pulse <0.1s prior to TA burst | 44 | 1.02 | 0.18 | 1.22 | 0.32 | 1.21 | 0.33 |
| Pulse during TA burst | 58 | 1.11 | 0.24 | 1.14 | 0.31 | 1.08 | 0.32 |
| Pulse after TA burst | 60 | 1.07 | 0.23 | 1.20 | 0.33 | 1.18 | 0.35 |

**Two-way ANOVA**

| Source of Variation | % of total variation | F (DFn, DFd) | P value |
|---|---|---|---|
| Interaction | 1.14 | F (6, 558) = 1.104 | 0.36 |
| Timing | 0.25 | F (3, 558) = 0.4777 | 0.70 |
| Metric | 2.89 | F (2, 558) = 8.419 | 0.0002 |

**Duration**

| Tukey's multiple comparisons test | Mean Diff. | 95.00% CI of diff. | Adjusted P Value |
|---|---|---|---|
| Pre far vs. Pre close | 0.016 | -0.1742 to 0.2062 | 1.00 |
| Pre far vs. During | -0.073 | -0.254 to 0.108 | 0.73 |
| Pre far vs. Post | -0.034 | -0.214 to 0.146 | 0.96 |
| Pre close vs. During | -0.089 | -0.2463 to 0.06825 | 0.46 |
| Pre close vs. Post | -0.05 | -0.2061 to 0.1061 | 0.84 |
| During vs. Post | 0.039 | -0.1058 to 0.1838 | 0.90 |

**Integral**

| Tukey's multiple comp | Mean Diff. | 95.00% CI of diff. | Adjusted P Value |
|---|---|---|---|
| Pre far vs. Pre close | -0.035 | -0.2252 to 0.1552 | 0.96 |
| Pre far vs. During | 0.044 | -0.137 to 0.225 | 0.92 |
| Pre far vs. Post | -0.013 | -0.193 to 0.167 | 1.00 |
| Pre close vs. During | 0.079 | -0.07825 to 0.2363 | 0.57 |
| Pre close vs. Post | 0.022 | -0.1341 to 0.1781 | 0.98 |
| During vs. Post | -0.057 | -0.2018 to 0.08784 | 0.74 |

**Amplitude**

| Tukey's multiple comparisoı | Mean Diff. | 95.00% CI of diff. | Adjusted P Value |
|---|---|---|---|
| Pre far vs. Pre close | -0.039 | -0.2292 to 0.1512 | 0.95 |
| Pre far vs. During | 0.092 | -0.08901 to 0.273 | 0.56 |
| Pre far vs. Post | -0.002 | -0.182 to 0.178 | >0.9999 |
| Pre close vs. During | 0.131 | -0.02625 to 0.2883 | 0.14 |
| Pre close vs. Post | 0.037 | -0.1191 to 0.1931 | 0.93 |
| During vs. Post | -0.094 | -0.2388 to 0.05084 | 0.34 |
